# Collagen-based scaffolds loaded with iron oxide nanoparticles promote functional sensorimotor recovery in spinal cord injury

**DOI:** 10.64898/2026.05.23.727171

**Authors:** Verónica Barranco-Maresca, Julia Martínez-Ramírez, María de Lamo-Atencia, Yasmina Hernández-Martín, Marina Sánchez-Petidier, Esther Benayas, Víctor Caz, Cristina Rosas, Raquel Madroñero-Mariscal, Elena Alonso-Calviño, Elisa López-Dolado, Juan Aguilar, María C. Serrano, Juliana M. Rosa

**Author notes:** **Authorship note:** VB-M and JM-R contributed equally to this work. **Corresponding authors:** Juliana M Rosa, Hospital Nacional de Parapléjicos, IDISCAM, Finca La Peraleda s/n, Neuronal Circuits and Behaviour Lab, planta 1, lab 5. Toledo, Spain. María C. Serrano, Instituto de Ciencia de Materiales de Madrid, CSIC, Calle Sor Juana Inés de la Cruz 3, 28049-Madrid, Spain.

## Abstract

Spinal cord injury disrupts sensorimotor circuits, leading to chronic deficits that require coordinated repair of both spinal and supraspinal circuits. Here, we developed and evaluated a hybrid collagen hydrogel containing chitosan-functionalized iron oxide nanoparticles as a therapeutic scaffold to promote multi-level recovery in a C6 hemisection model in rats. *In vitro*, both the nanoparticles and the resulting hybrid scaffold show preserved neuronal viability, excitability, and network connectivity. *In vivo*, scaffold-implanted rats demonstrate significant improvements in gross motor function and postural control, as well as recovery of fine motor skills, forelimb dexterity and grip strength. Sensory evaluations show preserved hindlimb tactile responses accompanied by plasticity within the somatosensory cortex, indicating functional recovery of the different tracts related to sensory and motor functions. At the lesion site, the scaffold enhances neurite outgrowth and modulates the inflammatory milieu, providing a permissive environment for neural repair. These findings indicate that these hybrid collagen scaffolds support relevant integrated structural and functional recovery features after SCI and represent a promising platform for further optimization toward the effective release of therapeutics at the injured spinal cord.

## INTRODUCTION

Spinal cord injury (SCI) disrupts the bidirectional flow of neural information between the brain and the spinal cord, leading to often irreversible deficits in sensory, motor and autonomic functions^1^. Functional recovery after SCI, particularly the restoration of skilled movements and adaptive sensorimotor control, requires the re-establishment of coherent interactions between ascending sensory feedback and descending motor commands^2,3^. Achieving this remains a central challenge as functional repair depends not only on axon regeneration across the lesion but also on the reorganization of supraspinal networks, including sensorimotor cortex, brainstem, and cerebellum^4–8^.

Over the past decades, biomaterial-based strategies have emerged as promising tools to guide and enhance such complex regenerative processes by providing structural support and modulating the local injury environment, thereby limiting secondary degeneration, promoting axonal sprouting, and generating permissive conditions for circuit-level reorganization^9,10^. Their capacity to deliver therapeutic cues directly at the lesion site further expands their utility, enabling local delivery of growth factors, gene therapeutics, and neuroprotective molecules^11,12^. Among available biomaterials, collagen hydrogels have attracted interest as scaffolds for SCI repair due to their ability to mimic the physiological extracellular matrix and support cell adhesion, proliferation, differentiation, and modulation of the inflammatory response^13,14^. For instance, fibroin-collagen hydrogels delivering induced neural stem cells enhanced survival, axon growth, and functional recovery^15^. Similarly, graphene-collagen cryogels reduced pro-inflammatory cytokines and microglial activation, improving the neuroinflammatory milieu post-SCI in rodent models^16^. In another study, electrospun collagen nanofibers were shown to guide neurite outgrowth and attenuate astroglial scarring in rat models, fostering neural sprouting and tissue organization^17^. In more complex settings, collagen hydrogels and conduits have been combined with bioactive cargos to boost cell viability and recovery (for review see^18^). Within this context, the incorporation of inorganic nanoparticles, like iron oxide nanoparticles (IONPs), in collagen hydrogels is gaining attention as an attractive strategy to enable SCI microenvironment modulation and magnetic-responsive therapies. Some studies showed promising results such as the promotion of neural elongation^19,20^ and peripheral nerve regeneration in rats through neurotrophic factor delivery from magnetic fibrin-collagen scaffolds^21^. In addition, IONPs can also mitigate free-radical–induced damage, improve the microenvironment for neural repair, enhance axonal regrowth, reduce lesion volume, and improve functional recovery, as shown in preclinical animal models^22,23^. Despite these encouraging findings, the synergistic potential of combining collagen hydrogels with IONPs remains incompletely characterized *in vivo*. Whether this hybrid strategy can amplify regenerative signaling, reinforce scaffold-mediated biomechanics in the paraplegic animal, or deliver superior neuroprotection and functional recovery compared with either component alone is still unknown.

In this study, we developed and evaluated the therapeutic efficacy of a hybrid collagen hydrogel containing chitosan-functionalized iron oxide nanoparticles (COLCHI) designed to enhance structural and functional repair following incomplete SCI. We first validated the biocompatibility and network-preserving properties of both the nanoparticles and the resulting hybrid scaffold *in vitro*. Following implantation in a rat model of SCI (C6 hemisection), our results revealed that COLCHI scaffolds not only promoted neurite growth at the lesion site but also accelerated the recovery of coordinated sensorimotor function. Importantly, statistical and network analyses indicated that this therapeutic approach strengthens adaptive cortical reorganization, suggesting that supraspinal circuit plasticity constitutes a key mechanism underlying the observed functional improvements.

## RESULTS

### Characterization of chitosan-functionalized iron-oxide nanoparticles

Nanoparticle formulations prepared for further loading into collagen hydrogels comprised uniform, single core iron oxide nanoparticles (IONPs) of 8 nm in diameter (**Figure 1A**, Bare). Given the colloidal instability at physiological pH values of the bare nanoparticles fabricated, we selected chitosan as a stabilizing and bioactive natural molecule, since it can provide neuroprotective properties to the resulting nanosystem^24,25^. Transmission electron microscopy (TEM) studies confirmed that IONPs functionalized with chitosan (*herein* NPCHI) preserved their spherical morphology after polymer coating (**Figure 1A**, NPCHI). At pH 7, dynamic light scattering (DLS) measures revealed a hydrodynamic size of around 46 nm for these coated nanoparticles (**Figure 1B**), which also displayed a narrower size distribution than their uncoated counterparts as they severely aggregated (reason why they cannot be shown in the graph). Surface charge measurements by DLS showed NPCHI yielding a slightly more negative charge (-8 mV) than their bare counterparts owing to the tripolyphosphate crosslinking method used for CHI coating (**Figure 1C**). TGA results indicated around 23 % of weight loss assigned to physically adsorbed water (up to 100 °C) and around 13 % of weight loss above 100 °C corresponding to the chitosan coating (**Figure 1D**). The heating capacity of NPCHI measured by the specific absorption rate (SAR) at 1 mg Fe/ml (**Figure 1E**) and corresponding magnetization curves (**Figure 1F**) remained unaffected by the chitosan coating. Specifically, both bare and NPCHI nanoparticles exhibited a superparamagnetic behaviour at room temperature, with saturation magnetization values around 60 emu/g for both of them and no hysteresis.

**Figure 1.**
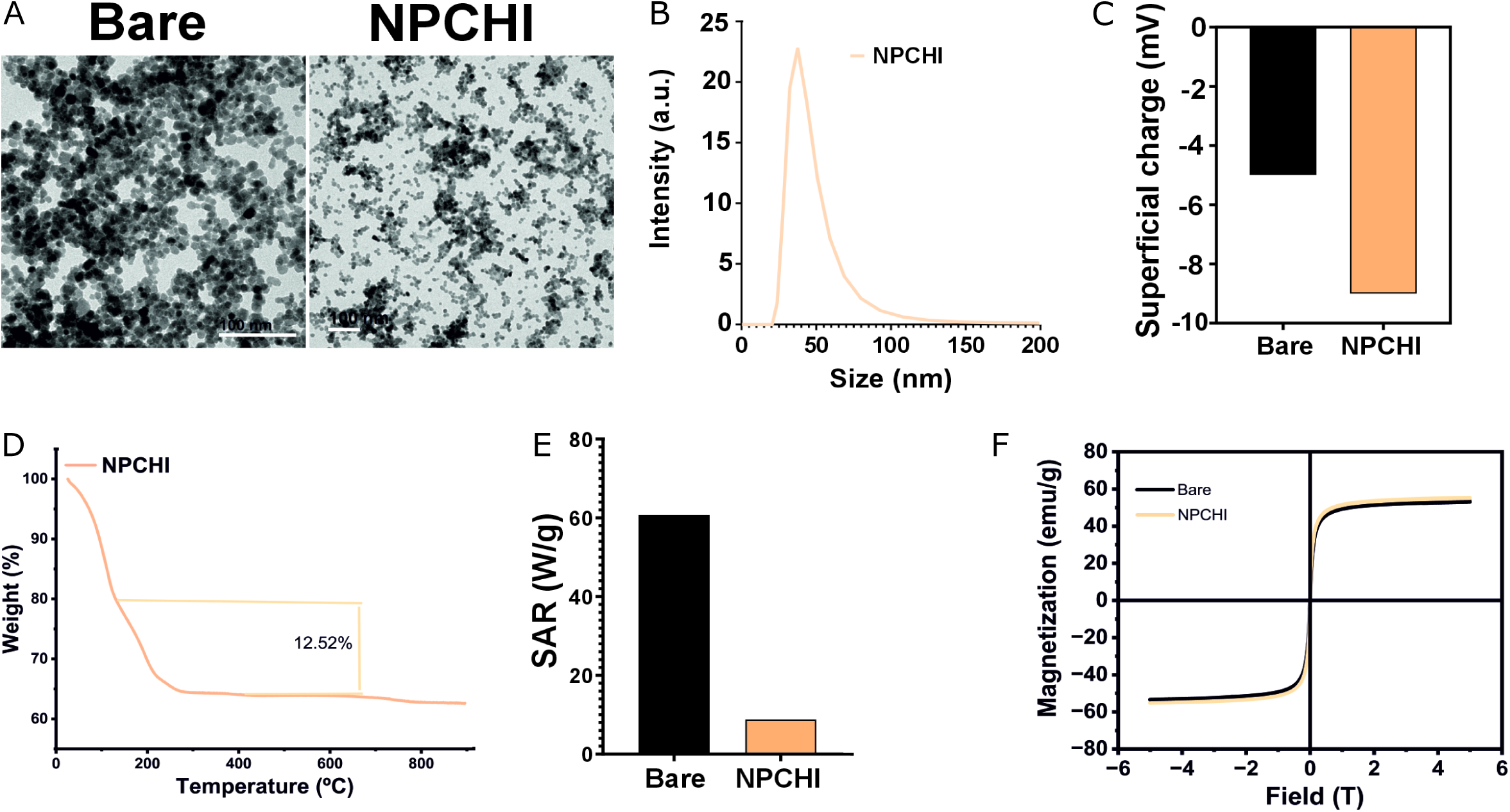
Physicochemical and biological characterization of chitosan-coated iron oxide nanoparticles (NPCHI). **A)** Representative TEM micrographs. Scale bar: 100 mm. **B)** Hydrodynamic size distribution in intensity measured by DLS at pH 7. **C)** Surface charge measured by DLS at pH 7. **D)** Thermogravimetric analyses (TGA). **E)** Specific absorption rate (SAR). **F)** Magnetization curves. IONP formulations tested: Bare IONPs (uncoated, black) and NPCHI (pale orange).

### NPCHI IONPs preserve cell viability, synaptic formation and network connectivity

After physical-chemical characterization of NPCHI, we evaluated their biocompatibility *in vitro*. Cultured embryonic neural progenitor cells (ENPCs) extracted from E17 rat embryos were exposed to two different NPCHI concentrations (0.01 mg Fe/mL and 0.05 mg Fe/mL) (**Figure 2A**). Cell viability, measured as calcein and ethidium homodimer 1 (EthD-1) positive area, was preserved in all conditions tested (**Figure 2A,B**), with no statistical differences with respect to untreated cells (ANOVA p > 0.05). To assess neuronal differentiation, we performed immunostaining for the vesicular glutamate transporter 1 (VGlut1) and glutamate decarboxylase 67 (GAD67) to identify excitatory (E) and inhibitory (I) neurons, respectively. Under control conditions, ENPCs spontaneously differentiated into E and I neurons at an approximate ratio of 8:2 (**Figure 2C**). NPCHI did not alter this E/I ratio or affect the number of primary neurites extending from neuronal soma (**Figure 2C**). Similarly, quantification of presynaptic (Homer) and postsynaptic (Bassoon) protein puncta revealed no significant differences in protein density or apposition/overlap synapses between NPCHI-treated and control ENPCs (**Figure 2D**). We next examined whether NPCHI affects neuronal intrinsic excitability and network connectivity *in vitro*. At day *in vitro* (DIV) 16^th^, when neuronal cultures have reached maturity and established initial network connections^26^, ENPCs were loaded with the membrane-permeable calcium indicator Fluo-4 AM to monitor intracellular Ca²⁺ dynamics as a proxy for neuronal activity (**Figure 2E**). NPCHI-treated ENPCs exhibited a modest increase in Ca²⁺ event frequency compared to controls (**Figure 2F**); however, network connectivity, as measured by mean pairwise correlation, remained unchanged. Together, these results demonstrate that NPCHI supports normal synaptogenesis and enhances neuronal excitability without compromising cell viability or network connectivity.

**Figure 2.**
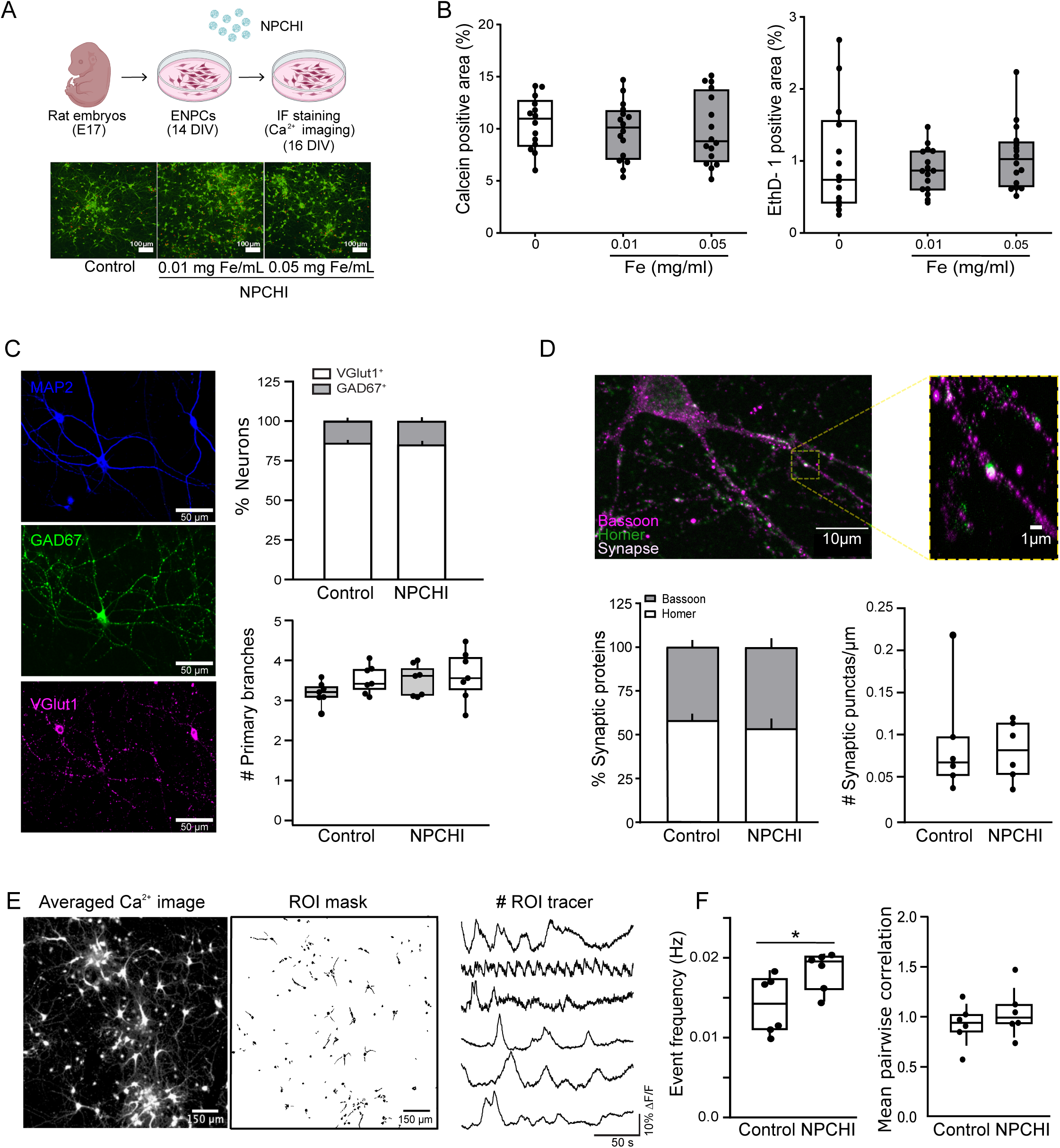
NPCHI maintains neural cell viability, differentiation, and neuronal synaptic and network connectivity. **A)** Top: Schematic of the *in vitro* experimental design used to assess the effects of NPCHI on primary neural cultures. Bottom: Representative viability confocal images of ENPCs exposed to NPCHI. Control corresponds to non-treated cells. Scale bars: 100 μm. **B)** Positive area for calcein (live/green cells in panel A) and ethidium homodimer 1 (EthD-1, dead/red cells in panel A) upon different NPCHI concentrations (0.01 mg Fe/mL and 0.05 mg Fe/mL). **C)** Left: Representative confocal images of ENPC cells labeled for MAP-2 (mature neurons, blue), GAD67 (GABAergic neurons, green) and VGlut1 (glutamatergic neurons, magenta). Right: Proportion of GABAergic and glutamatergic neurons expressed as a percentage of total MAP-2^+^ neurons and number of primary branches from both cell types (N = 7, n = 42). **D)** Representative images showing presynaptic marker Bassoon (magenta) and postsynaptic marker Homer (green). Colocalization (white) indicates functional synaptic punctas. Inset shows a magnified view of the area highlighted by the yellow box. Quantification of colocalized synapses per µm (N = 6, n = 36). **E)** ENPCs loaded with Fluo4-AM and the respective pseudo-automated segmentation showing the different regions of interest (ROIs). Examples of fluorescence changes denoting spontaneous activity of the ENPCs from 6 random ROIs. **F)** Frequency of spontaneous events (in Hz) and the mean pairwise correlation values from both conditions (N= 6, n = 3 per condition). Unpaired t-test * p < 0.05.

Next, we homogeneously dispersed NPCHI nanoparticles within a collagen I matrix forming hybrid hydrogels (*herein* COLCHI scaffolds, **Sup Figure 1**). Scanning electron microscopy (SEM) revealed soft, randomly porous, and fibrous architectures resulting from the freeze-casting methodology used (**Sup Figure 1A**). Transmission electron micrographs (TEM) confirmed the homogeneous NPCHI distribution throughout the hydrogel (**Sup Figure 1B**). *In vitro* cell studies revealed, as for the nanoparticles alone, a high cell viability on the matrix (**Sup Figure 1C**) and a predominant differentiation toward neuronal phenotypes (**Sup Figure 1D**). Together, the formulated hybrid hydrogels preserved cell viability and differentiation, fostering their exploration *in vivo* in animal SCI models.

### Hybrid collagen hydrogels boost sensorimotor integration in injured rats

Next, hybrid hydrogels were fabricated for *in vivo* implantation into a right C6 hemisection model of SCI in rats, enabling chronic evaluation of its therapeutic potential (**Figure 3A, Sup Figure 2A**). We then conducted longitudinal behavioral assessments using a battery of tests specifically designed to measure sensorimotor recovery. We first employed the horizontal ladder walking test, a well-established assay for evaluating sensorimotor integration (Metz & Whishaw, 2002) (**Figure 3B-H**). Under regular rung spacing (regular pattern, RP), which assesses rhythmic locomotion, a marked functional deficit was observed following SCI. These deficits included a significantly increased time to cross the ladder (**Figure 3C**) and a greater number of stepping errors (**Sup Figure 2C)** observed as a higher percentage of total miss paw placements and a corresponding decrease in correct/partial steps (**Figure 3D, Sup Table 1**). These impairments were most pronounced in ipsilateral forelimbs (FL) at 30 DPI, with a partial spontaneous recovery at 120 DPI. Ipsilateral hindlimbs (HL) also exhibited an increase of step errors, but less pronounced than in FL given the level of the SCI at C6 (**Figure 3E, Sup Figure 2C**). Notably, COLCHI-implanted rats showed values of crossing time similar to pre-injury time point (**Figure 3C-D**) and enhanced ipsilateral forelimb accuracy as early as 30 DPI (**Figure 3D**, **Sup Table 1**). Therefore, the enhanced rhythmic stepping and interlimb coordination observed in COLCHI rats suggest partial functional recovery largely mediated by propriospinal and reticulospinal circuits.

**Figure 3.**
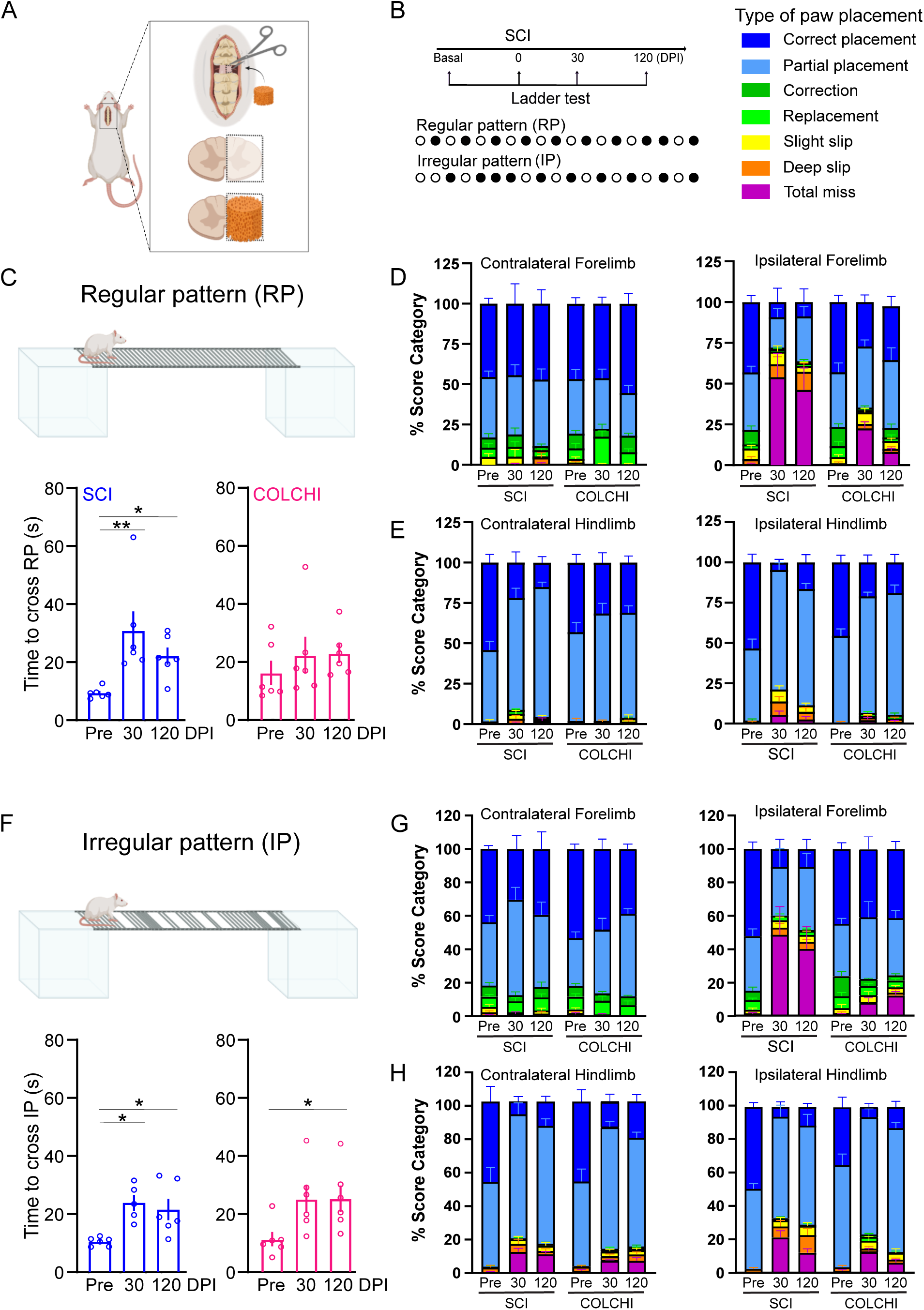
COLCHI scaffolds improve sensorimotor performance in the ladder test following SCI. **A)** Schematic representation of COLCHI scaffold implantation in rats with a right-sided C6 hemisection. **B)** Longitudinal ladder walking assessment using regular rung patterns (RP) or irregular rung patterns (IP). Color scale showing the types of paw placement used for the score category analysis. **C)** Time to cross the ladder under a regular pattern. **(D-E)** Percentage of paw placement categories during RP for FL and HL, respectively. **F)** Time to cross the ladder under an irregular pattern. **(G-H)** Percentage of paw placement categories during IP for FL and HL, respectively. FL: forelimb; HL: hindlimb. See **Sup. Table 1** for complete statistics. (N = 6 per group). Kruskal-Wallis test followed by Dunn’s post hoc, * p < 0.05, ** p < 0.01.

We next challenged the animals to an irregular rung spacing (irregular pattern, IP) to assess functions requiring forebrain circuits including step-by-step adjustment and body coordination^27^. Under IP conditions, COLCHI-implanted rats did not improve crossing times with respect to pre-SCI values, and functional deficits were observed at 120 DPI (p = 0.048) with a tendency at 30 DPI (p = 0.0502) (**Figure 3F-H**). Nevertheless, COLCHI rats showed a significant reduction in step errors **(Sup. Figure 2D)** and improved foot placement accuracy, indicating enhanced coordination and corrective control of ipsilateral FL and HL (**Figure 3G, Sup. Table 1**). In contrast to RP performance, HL displayed a higher number of step errors under IP conditions, likely reflecting compensatory weight-support strategies to correct foot misplacement (**Figure 3H; Sup. Figure 2D**). Importantly, control experiments in age-matched, uninjured rats confirmed that ladder test performance was not affected by aging *per se*, indicating a specific effect of the therapy (**Sup. Figure 3**). Taken together, COLCHI scaffolds enhance sensorimotor integration evidenced by improved mechanical and rhythmic stepping (under RP) and adaptive control (under IP), especially at the ipsilateral side.

### COLCHI scaffolds differentially affect motor and sensory pathways

Following the positive outcomes in the ladder test, we next used complementary behavioral assessments to dissect individual patterns of motor and sensory functional recovery (**Figure 4** and **Figure 5**). We started by placing animals into an open field arena to measure parameters of overall global activity. After SCI, rats significantly decreased their walking activity (**Figure 4Bi).** COLCHI rats spent significantly more time in spontaneous walking than their counterparts without hydrogel (group effect F_(1,10)_ = 5.33, p = 0.043) with distinct recovery trajectories between groups (time x group F_(5,50)_ = 3.114, p = 0.015). Overall, active behavior remained similar to pre-injury values at all time points except 120 DPI, where a decrease was observed which was more pronounced in the SCI group (pre vs. 120 DPI: p = 0.022) than in the COLCHI group (pre vs. 120 DPI: p = 0.097) (**Figure 4Bii**). We then measured the ability of animals to lean on their FL during stand-up movements (**Figure 4C**). Both injured groups reduced the frequency of stand-ups using both FL (**Figure 4Ci**) or the ipsilateral forelimb alone (**Figure 4Cii**), while increasing the use of the contralateral (unaffected) forelimb (**Figure 4Ciii**). COLCHI outperformed SCI rats, using both FL more frequently to lean on the cage, an effect that emerged as early as 14 DPI indicating partial recovery at later time points. Additionally, COLCHI animals maintained pre-injury levels of time devoted to grooming after the injury (**Figure 4Biii**), whereas SCI only showed altered grooming patterns (time × group F_(5,50)_ = 3.285, p = 0.012). Spontaneous grooming, while representing a self-care activity, can indicate the presence of dysesthesia or neuropathic pain induced by hyperexcitability and sensory disorganization after injury^28–30^. The preservation of normal grooming patterns in COLCHI animals may therefore suggest mitigation of abnormal somatic signaling following SCI. To further clarify the magnitude of these differences, grooming at the different head reaching positions was investigated (**Figure 4D, Sup Figure 4**). A two-way ANOVA analysis from the use of contralateral forelimb to reach position 3 (between eyes and ears) revealed a significant group x time interaction (F_(5,50)_ = 3.393, p < 0.05), indicating COLCHI increased the use of the unaffected forelimb (**Figure 4Di**). No significant differences were found in the use of contralateral or ipsilateral FL to reach ears’ (position 4) and beyond ears’ (position 5), which require greater forelimb extension and postural demand (**Figure 4Dii-iii, Sup Figure 4**). These results suggest that COLCHI hydrogel facilitates the compensatory use of the unaffected forelimb for mid-range grooming movements, while higher-demand reaching positions remain persistently impaired in both lesioned groups.

**Figure 4.**
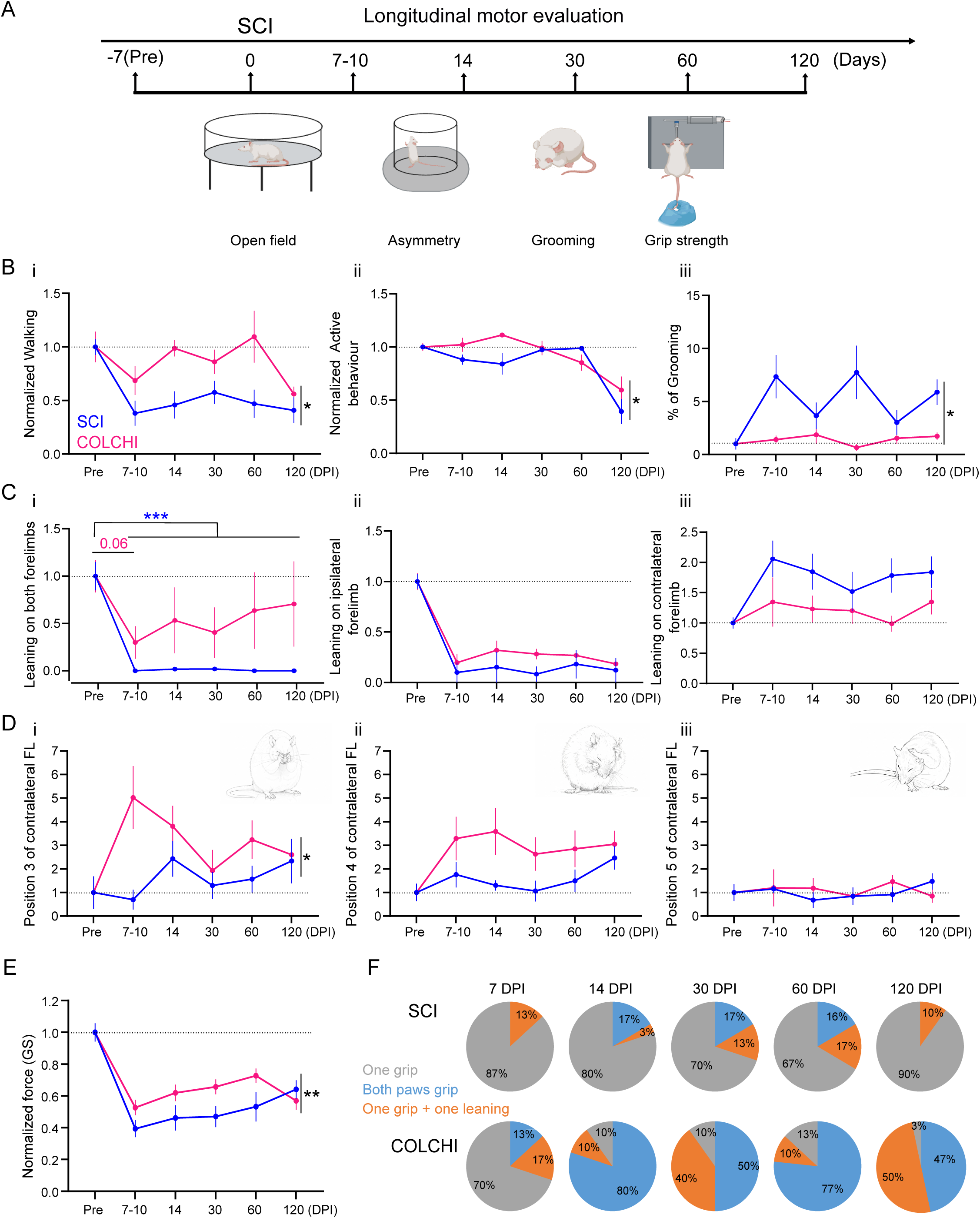
Motor behavioural assessment of COLCHI-implanted tetraplegic rats. **A)** Schematic of the longitudinal protocol used to evaluate motor recovery. **B)** Open field examination of rat’s walking performance (i), active behaviour (ii) and grooming (iii). **C)** Asymmetry test showing the ability of tetraplegic rats to lean on both FLs (i), the ipsilateral FL (ii) or the contralateral FL (iii) while standing up. **D)** Grooming assessment with focus on the use of the contralateral FL to reach positions 3 (eyes’ area), 4 (ears’ area) and 5 (beyond ears’ area). **E)** Grip strength assessment of FL muscle strength. **F)** Percentage of FL used to pull the bar during grip strength test. (N = 6 per group for all behavioral tests). Two-way repeated measures ANOVA (panels C_iii_, D_ii_) or linear mixed model. FL: forelimb. See **Sup. Table 2** for complete statistics. For clarity, asterisks denote time x group interaction, except on Panel Ci where it indicates differences between time post SCI to pre-SCI values (* p < 0.05; ** p < 0.01; *** p < 0.001).

**Figure 5.**
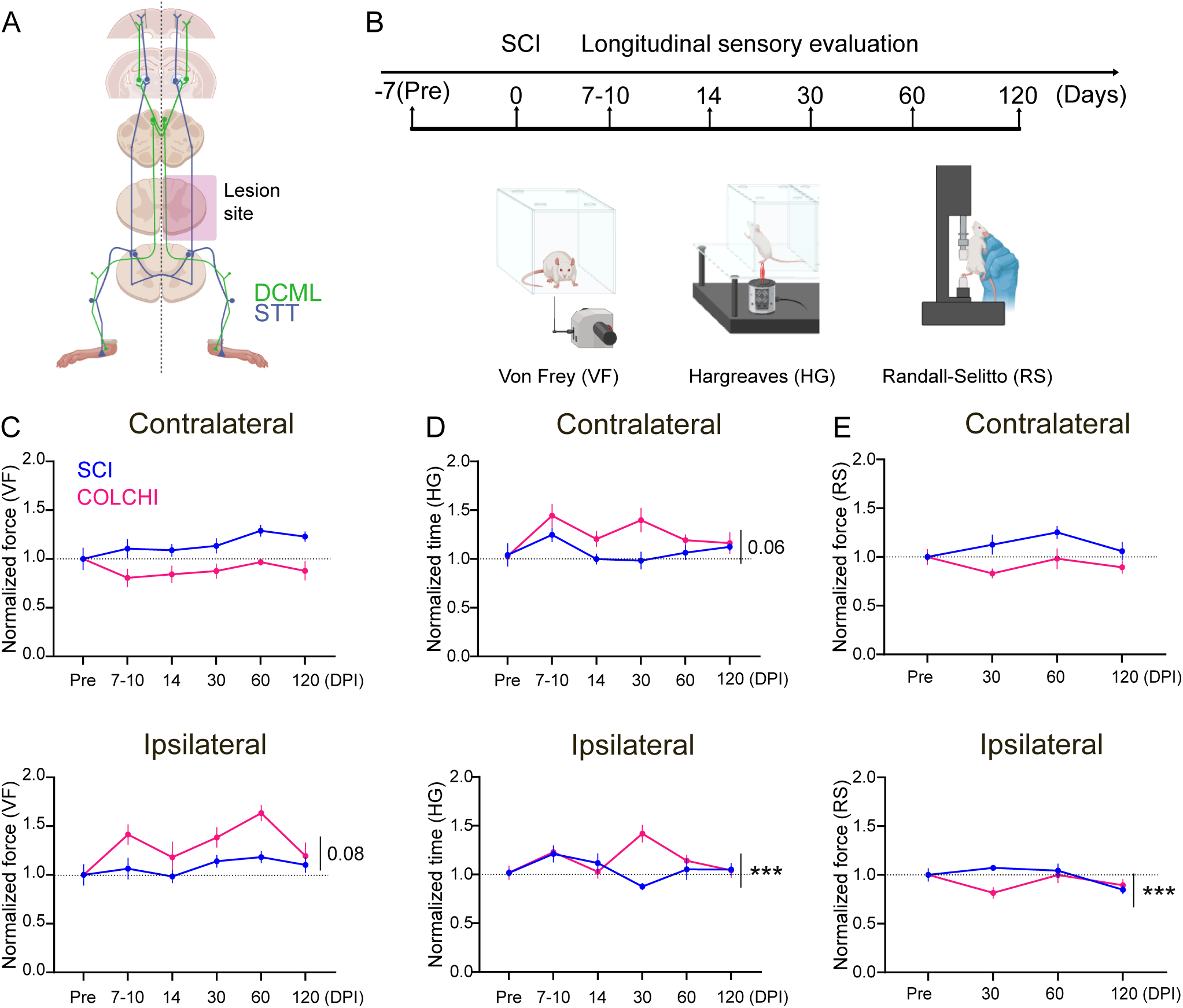
COLCHI scaffolds produce changes in sensory circuitry in longitudinal behavioral studies. **A)** Diagram of the ascending sensory pathways affected by the C6 hemisection. **B)** Schematic of the different sensory tests (Von Frey, Hargreaves and Randall-Selitto) showing the time points of longitudinal assessments. **C)** Normalized force to induce HL withdrawal in the dynamic Von frey stimulation of the contralateral (above) and ipsilateral (below) hindlimb. **D)** Normalized time to Same as C but using an infrared beam in the Hargreaves test. Normalized response (thermal stimulation using the Hargreaves plantar test **(D)**, and to the force applied using the Randall-Selitto test for nociception **(E)**. See **Sup. Table 3** for complete statistics. N = 6 per group. Two-way repeated measures ANOVA (panel E) or linear mixed model. For clarity, asterisks denote time x group interaction. *** p < 0.001.

Next, we assessed voluntary forelimb dexterity and muscle strength using the grip strength test^31,32^. Injured rats, in both groups, exhibited a marked loss of forelimb muscle strength after SCI that gradually recovered over time (F_(3.4,34,02)_ = 40.36, p < 0.0001, **Figure 4E**) with a significant group × time interaction (F_(5,50)_ = 3.418, p = 0.009), indicating that both groups differed in their recovery trajectory. This difference was observed at 60 DPI where COLCHI rats presented a higher recovery when compared to 7-10 DPI time point (**Sup Table 1**). Moreover, animals with COLCHI scaffold - but not SCI only - showed a drastic and progressively increased use of the affected ipsilateral paw to pull the bar, starting from 7 DPI and sustained through the end of the study (**Figure 4F**, time x type of gripping F_(8,60)_ = 8.459, p < 0.0001 for COLCHI; time x type of gripping F_(8,60)_ = 1.42, p = 0.207 for SCI). A late decline in bilateral paws use was observed at 120 DPI in both groups, consistent with the reduction in walking activity, active behaviour and gripping force described above. Notably, the remaining forelimb dexterity for COLCHI-implanted rats at 120 DPI was superior to that of SCI-only animals, indicating COLCHI improvement over the SCI condition. Globally, our results obtained from the motor tests applied indicate that COLCHI improves descending motor pathways and spinal circuits leading to increased global mobility, forelimb coordination, muscle strength and voluntary motor output after SCI as early as 7 DPI and up to 120 DPI for most of the parameters investigated.

To determine whether COLCHI scaffolds affected sensory circuits, we longitudinally evaluated the functions of the two main ascending pathways (*i.e.,* dorsal column-medial lemniscal (DCML) and spinothalamic tracts (STT)) using complementary behavioral tests (**Figure 5A,B**). DCML-mediated mechanical sensitivity was studied by using the dynamic Von Frey applied to HL. This approach allowed us to distinguish between the preserved contralateral DCML tract and the injured ipsilateral tract, providing insight into both compensatory mechanisms and lesion-specific deficits (**Figure 5A**). Following contralateral stimulation, a strong effect of the group was observed (F_(1,10)_ = 24.33, p < 0.001), with COLCHI animals exhibiting consistently lower withdrawal thresholds (**Figure 5C**). This effect was stable across time points, as neither a main effect of time nor group x time interaction was detected (**Sup Table 2**). In contrast, following ipsilateral stimulation, significant main effects of both group (F_(1, 10)_ = 5,462, p = 0.041) and time (F_(5,50)_ = 6,301, p = 0.0019) were observed, without interaction. In this case, COLCHI rats exhibited higher withdrawal thresholds over time, consistent with reduced mechanical hypersensitivity. The absence of interaction suggests that the temporal evolution was similar between groups, with COLCHI rats maintaining an overall decreased mechanical sensitivity. These results indicate that COLCHI induced side-specific effects on mechanical sensitivity, exacerbating contralateral hypersensitivity while attenuating ipsilateral mechanical responsiveness, pointing to distinct lateralized mechanisms of sensory modulation.

Thermal responsiveness was next assessed using the Hargreaves test set to 38 °C, a temperature used to activate thermal afferents without reaching the nociceptive threshold (**Figure 5D**). In this case, stimulation of the ipsilateral HL relies on preserved ascending pathways - given that the STT decussates shortly after entering the spinal cord, while contralateral HL activates injured STT (**Figure 5A**). Our results revealed that COLCHI significantly altered the thermal responsiveness to ipsilateral stimulation (*i.e.* spared STT) following SCI (group x time F_(5,50)_ = 6.981, p < 0.0001). *Post-hoc* analysis localized this effect to 30 DPI, a time point where secondary injury processes and pain following SCI has been described^33,34^. In contrast, no significant interaction was observed in the contralateral paw (F_(5,50)_ = 2.233, p = 0.065) indicating that COLCHI modulates ipsilateral thermal responsiveness, likely by attenuating secondary injury or promoting adaptive plasticity within spared sensory circuits during the critical subacute-to-chronic transition of SCI.

Finally, we assessed mechanical nociception on distal limbs using the Randall-Selitto test (**Figure 5E**). In this case, early post-injury time points were deliberately skipped as the acute inflammatory milieu can profoundly lower mechanical nociceptive thresholds in a non-specific manner, thereby confounding the interpretation of the results. Following contralateral HL stimulation, the temporal profile of the response did not differ significantly between SCI and COLCHI rats (F_(3,30)_ = 2.177, p = 0.111), indicating comparable dysfunction of the directly injured pathway across groups. In contrast, ipsilateral stimulation showed a significant group x time interaction (F_(3, 30)_ = 4,928, p = 0.0067). *Post-hoc* analysis localized this effect to 30 DPI (p = 0.0205), where COLCHI rats exhibited lower mechanical withdrawal thresholds (i.e., increased sensitivity) compared to SCI group. Interestingly, our data showed mechanical hypersensitivity in the ipsilateral paw at 30 DPI that stands in apparent contrast to the thermal hypoalgesia observed at the same time point in the Hargreaves test, pointing to a modality-specific reorganization of STT function. The opposite effects of COLCHI on these two sub-modalities suggest differential modulation of dorsal horn circuitry or fiber-type-specific changes in excitability within spared ascending pathways, rather than a uniform effect on STT function.

### Cortical plasticity of somatosensory regions in COLCHI implanted rats

Our data so far demonstrates that COLCHI scaffolds promote the recovery of sensorimotor functions, which may arise from spinal circuit plasticity (with or without neuronal regeneration) or through changes in supraspinal regions involved in sensorimotor integration. To investigate this, we performed *in vivo* electrophysiological recordings from the HL coordinates of the primary sensorimotor cortex (HLCx) to assess sensory-evoked potentials (SEPs) following HL stimulation at 120 DPI (**Figure 6A)**. In SCI-only animals, SEPs recorded in the contralateral HLCx in response to ipsilateral HL stimulation exhibited a rightward shift in input–output recruitment curve with a marked increase in the Ihalf values corresponding to the input needed to induce a measurable response (**Figure 6B)**. This shift in the dynamic of cortical recruitment indicates impaired ascending signal transmission and reduced cortical responsiveness. On the contrary, recordings from the ipsilateral HLCx in response to contralateral HL stimulation did not show significant changes in the recruitment curve (**Figure 6C**), denoting preservation of low-threshold ascending fibers. Notably, COLCHI significantly increased cortical excitability, as evidenced by enhanced SEP responses at lower intensities and a leftward shift of the recruitment curve towards control values (group x intensity interaction F_(22,100)_ = 4.651, p < 0.0001; I_half_ SCI x COLCHI p = 0.002). This effect was selective to the left hemisphere, as SEP responses recorded from the right (contralateral, largely spared) hemisphere were comparable across SCI and COLCHI groups (group x intensity interaction F_(22,95)_ = 0.965, p = 0.513). Together, these data indicate that COLCHI scaffolds induce long-term plastic changes within supraspinal sensory circuits, enhancing the efficacy of ascending input to the sensorimotor cortex and providing a cortical-level substrate for the observed improvements in sensorimotor function.

**Figure 6.**
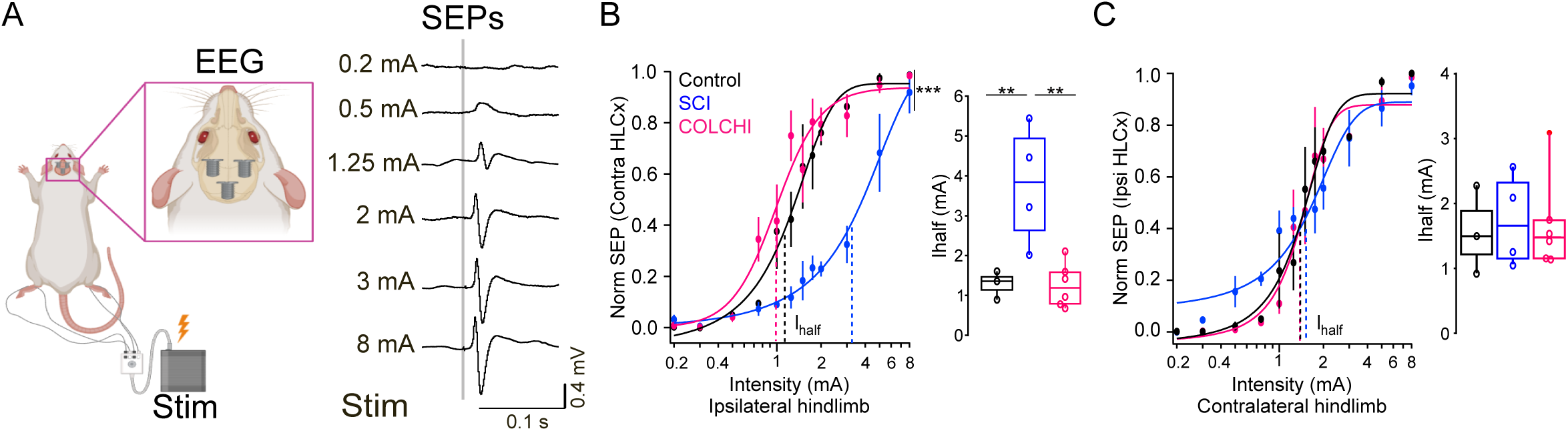
COLCHI scaffolds increase cortical excitability after SCI. **A)** Left: Scheme of the bilateral EEG recordings obtained from the HLCx in anesthetized animals in response to increasing HL electrical stimulation. Right: Example traces showing SEPs to different intensities. **B-C)** Input:output recruitment curves showing the population response for control healthy animals (black), injured animals without scaffold (SCI, blue) and COLCHI animals (magenta) when stimulating the ipsilateral HL (B) or contralateral HL (C). Dots represent the avg ± SEM value from 30-50 stimuli for each intensity. Dashed lines represent the I_half_ value obtained from Hill equation fitting. Left: **I**_half_ obtained from each animal within the respective group. Mixed-effects analysis (for the I:O curves) or one-way ANOVA (Ihalf) and *post hoc* Tukey’s test. ** p < 0.01, *** p < 0.001.

### Multifactorial analysis shows that COLCHI improves functional recovery at long term

Our data indicates that COLCHI modulates both spinal and supraspinal circuits, with differential contributions across ascending and descending pathways. To integrate these distributed effects at the level of global functional recovery - rather than isolated outcomes - we performed a multidimensional principal component analysis (PCA) using endpoints metrics from individual animals (**Figure 7**). In the PC1–PC2 space (**Figure 7A**), control animals clustered at positive PC1 values, reflecting strong contributions from motor-related variables, while SCI animals were shifted toward negative PC1 and PC2 values. COLCHI animals occupied an intermediate position along PC1 and were clearly separated from SCI along PC2. Loadings analysis (**Sup Figure 5A-B**) indicated that PC1 was primarily driven by motor performance variables, whereas PC2 loaded preferentially on cortical excitability. Thus, the rightward displacement of the COLCHI relative to SCI along PC1 indicates partial recovery of motor-related functions, while the upward shift along PC2 reflects enhanced cortical processing. A similar organization was observed in the PC1–PC3 projection (**Figure 7B**). In this case, SCI animals formed a compact cluster at negative PC1 values, whereas control animals clustered at positive PC1 with similar PC3 contributions. COLCHI animals again occupied an intermediate position along PC1, with PC3 values partially overlapping those of SCI animals. Variables loading onto PC3 included sensory responses (**Sup Figure 5A-B**), suggesting partial restoration of sensory functions in COLCHI-implanted animals.

**Figure 7.**
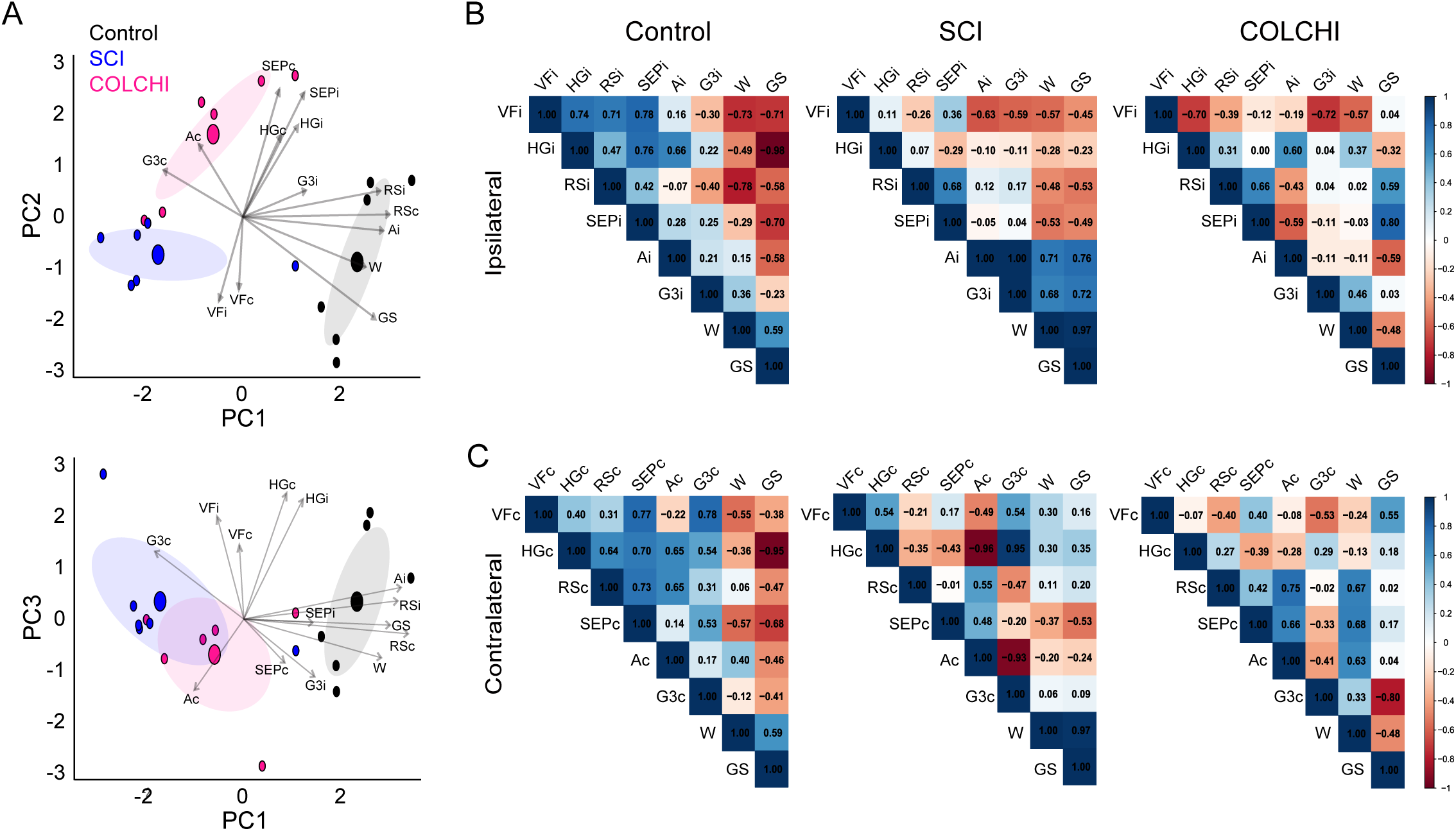
COLCHI promotes the reorganization of the sensorimotor functional network after SCI. **A)** Biplot of the PCA showing the distribution of the control (black), SCI (blue) and COLCHI (magenta) groups and the contribution of sensory and motor variables including contralateral and ipsilateral data. **B-C)** Correlation heatmaps of the different groups from the contralateral **(B)** and ipsilateral **(C)** sides. Legend: SEP: sensory-evoked potentials; VF: Von Frey; HG: Hargreaves; RS: Randall-Selitto; GS: grip strength; A: asymmetry; G3: grooming position 3; W: walking activity (c = contralateral, i = ipsilateral).

Across both projections, SCI showed decreased dispersion along PC1 and a displacement away from control loadings, whereas COLCHI treatment restored separation across principal dimensions and better re-aligned key sensory and motor variables toward control-associated directions. Notably, COLCHI animals did not simply overlap with controls, but instead formed a distinct cluster characterized by intermediate motor recovery and enhanced sensory–cortical contributions, consistent with partial but coordinated functional restoration. Together, these results demonstrate that COLCHI scaffolds promote long-term improvement at the systems level by shifting the global functional state away from the SCI-associated domain and toward a control-like organization that integrates motor performance with sensory and cortical excitability measures.

To further assess system-level recovery, we analyzed pairwise correlations among cortical, sensory and motor variables for each experimental group (contra and ipsilateral) (**Figure 7B-C**). In control animals, sensory and motor responses showed weak correlations, consistent with largely independent functional domains. After SCI, correlations derived from the injured ipsilateral side were significantly reduced, whereas contralateral responses showed increased correlation between sensory and motor variables. Notably, this cross-domain coupling was further modulated in COLCHI-treated animals, which displayed slight decrease in correlations between some of the tests with significant changes in our study such as walking, grip strength or grooming. Among the measured features, SEPs displayed weak correlation with motor performance in control and SCI groups in both hemispheres. In COLCHI-implanted animals, these correlations were increased in the contralateral side, which could explain some of the beneficial effects observed at motor behaviour. Therefore, COLCHI treatment not only improves individual functional measures but also enhances coordinated coupling between cortical excitability and sensorimotor performance. Together, these data support a role for COLCHI-induced cortical plasticity in promoting integrated functional recovery following SCI.

### COLCHI scaffolds modulate the spinal cord tissue response

After extensive behavioral and electrophysiological assessments, we next studied the spinal tissue at the lesion site. Representative intra-surgical images have been included for visualization of the injury done at C6 and the appropriate positioning of the COLCHI hydrogel inside the cavity created (**Sup Figure 2A**). Over the 120 DPI of examination, all rats showed weight gain, with COLCHI rats not differing from control rats (ANOVA, p = 0.985) and SCI-alone rats exhibiting a less pronounced weight gain (ANOVA, p = 0.026 with respect to control rats and p = 0.019 with respect to COLCHI rats). Trichrome staining revealed abundant collagen deposition within the lesion site, particularly at the tissue interfaces, with more pronounced fibrosis in SCI rats without COLCHI scaffolds (**Sup. Figure 6A-B**). Curiously, COLCHI rats showed numerous round and brownish cells within the lesion, consistent with macrophages that migrated to phagocyte the IONPs contained in the hydrogel and stayed chronically at the lesion site. Importantly, an almost complete biodegradation of the COLCHI scaffold was evident at this chronic time point, with a total area of tissue cavities of 24 ± 6 %, which was slightly larger than that found in SCI rats (16 ± 8 %; ANOVA, p = 0.123) and significantly larger than that in control rats (2 ± 1 %; ANOVA, p < 0.005).

Following gross inspection, the spinal cord was analyzed by immunofluorescence (**Figure 8, Sup. Figure 7**). Injured animals showed a marked increase in cell density at the lesion site and its interfaces compared with perilesional (PL12), left hemicord (LH), and control regions (ANOVA, p < 0.005; **Figure 8A**), consistent with injury-induced cell migration and proliferation. COLCHI-treated rats exhibited an additional increase in cell density in PL12 and the rostral interface (RIF) compared with both SCI and control groups (ANOVA, p < 0.005), indicating a more populated lesion environment. When focused on inflammatory markers, macrophages notably invaded the lesion and corresponding interface tissue (ANOVA, p < 0.005; **Figure 8B**). Indeed, the abundance of ED1^+^ cells at the lesion site was significantly higher than at the rostral interface for both injured groups (ANOVA, p = 0.035 and p = 0.032 with respect to lesion site for SCI and COLCHI, respectively), but not for the caudal interface (ANOVA, p > 0.05). At PL12, injured rats without hydrogel showed a significantly higher abundance of macrophages than control rats (ANOVA, p = 0.046). Contrarily, COLCHI rats did not show such differences with control rats at those perilesional regions. Vimentin⁺ cells were similarly enriched at the lesion site and its interfaces in both injured groups, independent of COLCHI implantation (ANOVA, p < 0.005, **Figure 8C**). Likewise, GFAP⁺ astrocytes were significantly increased at the lesion borders in both groups (ANOVA, p < 0.005; **Figure 8D**), although SCI animals showed a reduction in astrocyte density at the lesion site relative to the rostral interface, an effect not observed in COLCHI-treated rats.

**Figure 8.**
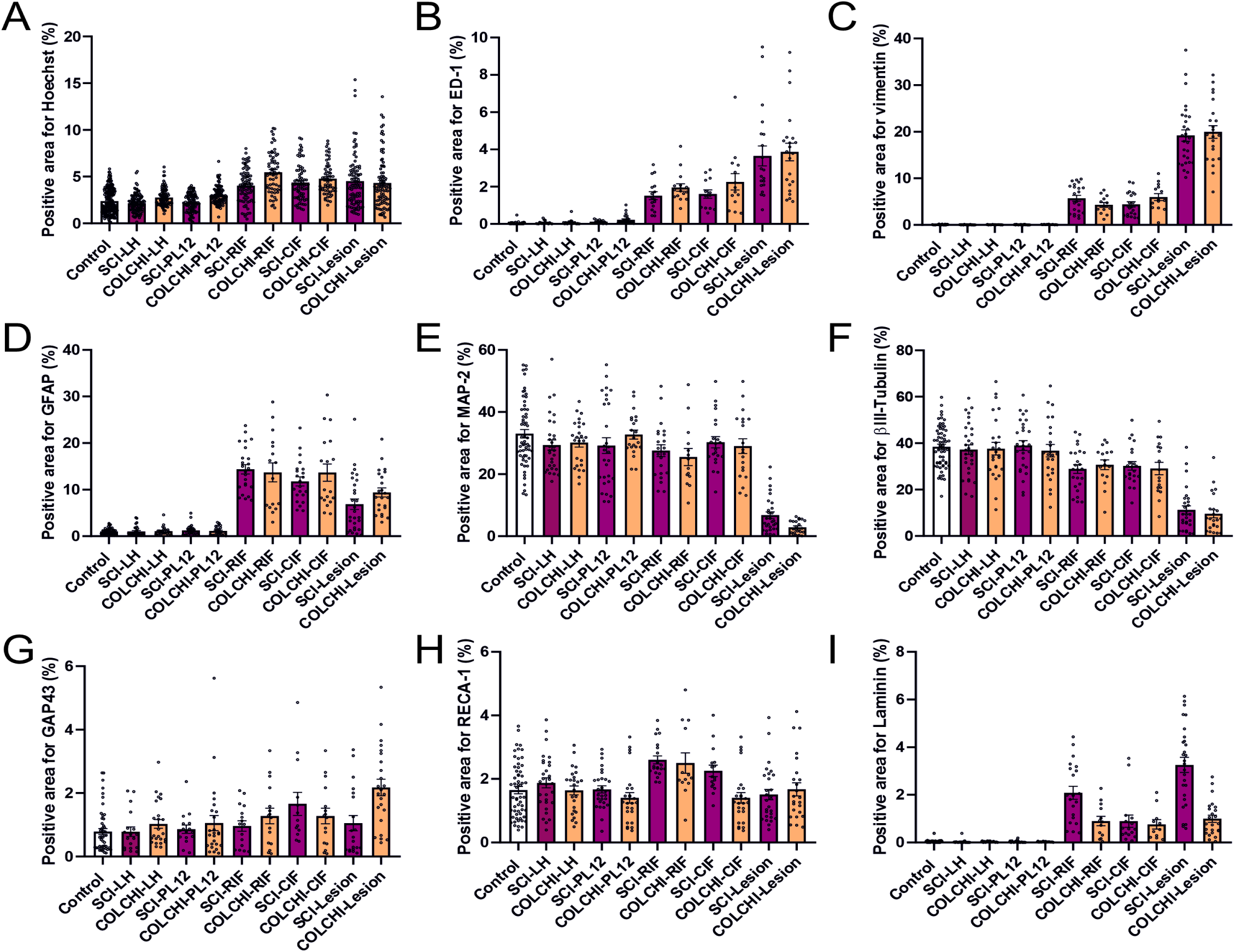
Immunofluorescence studies of the spinal cord tissue of COLCHI-implanted paraplegic rats. Selected markers included cell nuclei (Hoechst; **A**), macrophages (ED-1; **B**), non-neuronal cells (vimentin; **C**), astrocytes (GFAP; **D**), neurons (MAP-2 and βIII-tubulin; **E** and **F**, respectively), growth cones (GAP43; **G**), and vascular structures (RECA-1 and laminin; **H** and **I**, respectively). Statistically significant differences among treatment groups are described in the text for the good of graph clarity. Representative confocal images of all markers, treatment groups and areas are included in the supporting information (**Sup** Figure 7).

We next examined neuronal markers. MAP-2 (**Figure 8E**) and βIII-tubulin (**Figure 8F**) were similarly reduced at the lesion site in all injured animals (ANOVA, p < 0.005), with no differences in other regions compared with controls (ANOVA, p > 0.05). In contrast, COLCHI-implanted rats showed a significant increase in GAP43 at the lesion site relative to controls and to LH and PL12 regions in SCI rats (ANOVA, p < 0.005; **Figure 8G**), indicating enhanced activation of axonal growth in the presence of the hybrid hydrogel. Finally, vascular markers indicated substantial revascularization in both injured groups, with a selective increase in RECA-1 at the rostral interface in SCI animals (ANOVA, p < 0.005; **Figure 8H**). Laminin remained elevated at the lesion site and interfaces in both injured groups but was significantly higher in SCI than in COLCHI animals at the lesion and rostral interface. Overall, these results indicate that COLCHI scaffolds promote neurite growth within the lesion site while limiting excessive perilesional inflammation, thereby fostering a permissive microenvironment that supports regenerative responses and facilitates functional recovery.

Finally, we inspected the systemic toxicity of these hybrid collagen hydrogels after chronic implantation at the injured spinal cord. Hematoxylin-eosin staining revealed no signs of toxicity in major organs such as the spleen, the liver, the kidney and the lung (**Sup. Figure 8A**). When focused on their weight percentages, no significant changes were observed for any of the organs investigated except for the liver, which experienced a significant decrease in weight in SCI rats in comparison to COLCHI rats (ANOVA, p = 0.008; **Sup. Figure 8B**). The examination of the peripheral muscles (*triceps braquiae* and *triceps surae* of FL and HL, respectively) did not reveal remarkable changes in their weight percentage (ANOVA, p > 0.05; **Sup. Figure 8C**). Interestingly, the volume of the brachial triceps of the ipsilateral (affected) FL was significantly reduced in injured rats without implant (ANOVA, p = 0.012 with respect to control), while preserved in rats with the COLCHI hydrogel.

## DISCUSSION

Recent efforts in SCI research have increasingly focused on the use of biocompatible materials to promote neural repair and functional recovery. In this study, a hybrid collagen-based hydrogel incorporating chitosan-functionalized iron oxide nanoparticles (COLCHI) was investigated as a novel therapeutic scaffold in a rat model of incomplete SCI. *In vivo* implantation of COLCHI significantly enhanced the recovery of sensory and motor functions that were associated with increased neurite growth at the lesion site, reduced perilesional inflammation and adaptive reorganization of cortical circuits, indicating that COLCHI engages both local and supraspinal mechanisms of repair. Notably, although the COLCHI scaffold gradually degraded over time *in vivo*, its transient presence was sufficient to induce long-lasting neuroplastic and functional recovery, suggesting that further scaffold optimization could yield even greater therapeutic efficacy.

Our findings provide compelling evidence that COLCHI constitutes a suitable and promising biomaterial platform for SCI repair. At motor level, it provided long-term benefits for gross motor function and postural control, evidenced by improved performance in the regular spacing ladder test, asymmetry test and percentage of active time walking. These behavioral features primarily depend on the reticulospinal (RST) function modulating both lumbar central pattern generators to initiate basic patterns of locomotion, and propriospinal interneurons for the predicted interlimb coordination^35,36^. Additionally, RST is also the main driver of axial and proximal limb muscle tone providing stability and balance during locomotion^37^. The fact that most COLCHI beneficial effects on injured animals last until the end of the experimental protocol (120 DPI) indicates a robust and sustained circuit plasticity that is not affected by the eventual biomaterial degradation. Furthermore, COLCHI scaffolds also improved performance in grip strength, irregular ladder test and grooming, all requiring a high degree of fine, skilled and voluntary movements of distal muscles controlled by the corticospinal tract (CST)^38,39^. Circuit reorganization at the level of C3-C5, a critical anatomical and functional hub for forelimb motor control and recovery following SCI, could be involved in this functional recovery. For example, motor cortex plasticity may promote improved forelimb control through CST fibers reaching intact spinal cord segments at C3–C5, thereby enhancing control of proximal forelimb musculature in COLCHI animals^40,41^. Additionally, the preservation of motor neurons above the lesion level (C3–C5) may also facilitate axonal sprouting toward denervated muscle fibers in more proximal muscles (Jindal et al., 2026). Taken together, both mechanisms could contribute to the functional improvements observed. However, the parallel decline of the functional recovery in the grip strength test at more chronic time points suggests that the therapeutic effects of COLCHI on CST-mediated recovery are not sustained on time - an effect that may be related to the progressive biomaterial degradation. It is plausible that the COLCHI-induced new connections between spared CST fibers and propriospinal circuits may not be robust enough to maintain the initial functional improvement, leading to a loss of muscle strength and adaptive motor control at chronic time points^41,42^. Nonetheless, COLCHI rats retained functional improvement compared with SCI only, as evidenced by greater bilateral forelimb use during gripping. These results indicate that functional recovery preserved at 120 DPI is at least partly supported by spared CST fibers at C3-C4 within the injured hemicord, which continue to contribute to forelimb control. Moreover, the sustained improvement in complex locomotor tasks, such as the irregular ladder test, likely reflects compensatory reorganization of parallel descending fibers, including RST, contralateral CST projections and propriospinal networks capable of bridging spinal circuits above and below the lesion. Such circuit-level remodeling has been shown to support recovery when primary CST pathways are compromised (Cao et al., 2021; Freund et al., 2006) and may underlie the persistent sensorimotor improvements observed following COLCHI implantation.

COLCHI scaffolds also modulated the responsiveness of ascending sensory pathways, which are differentially affected in the intact versus injured hemicord. By assessing sensory responses from both HL in our hemisection SCI model, we were able to dissect the impact of COLCHI on distinct ascending systems. In this model, the injured ipsilateral hemicord directly disrupts STT fibers conveying sensory input from the contralateral paw, while DCML fibers originating from the ipsilateral paw are also compromised. In SCI animals, overall sensory responsiveness remained close to baseline over time, despite direct damage to these ascending tracts. This apparent preservation of responsiveness is unlikely to reflect true sensory recovery and instead is consistent with enhanced reflex response of distal limbs, a well-known consequence of SCI in both patients and experimental models^43^. In this case, the loss of descending inhibitory control from CST to the dorsal horn interneurons after SCI^3^ lead to increased excitability of ascending pathways, influencing the output behaviour and supporting aberrant responses^44^. In contrast, COLCHI-implanted animals exhibited a pattern of sensory responsiveness more consistent with the expected physiological consequences of tract injury, namely increased sensory thresholds reflecting altered tactile and thermal sensitivity. These findings suggest that COLCHI may be attenuating maladaptive plasticity within sensory-motor circuits, thereby limiting the development of hyperreflexia and preserving more physiological sensory responses. The spared ascending fibers traversing the intact hemicord could also contribute to the maintenance of responsiveness via compensatory mechanisms^45^; however, the bilateral nature of the effect argues against this being the primary driver. Nonetheless, COLCHI effects could be explained by its regenerative response promoting neurite growth within the lesion site and reducing perilesional inflammation, which creates a more permissive microenvironment that allows for heightened plasticity and reorganization of spared circuits at the spinal cord. Possible sprouting of CST acting on STT and DCML tracts to establish new relay connections or strengthening existing ones could also play an important role as they have been shown to promote functional recovery after SCI^3,46^.

Systems-level analysis revealed that COLCHI scaffolds restore coordinated integration of motor, sensory, and cortical networks disrupted by SCI, rather than merely improving isolated tasks. PCA demonstrated that SCI induces motor impairment, sensory/cortical dispersion, and sensorimotor fragmentation, mirroring multidimensional deficits observed in human patients. COLCHI repositioned animals toward uninjured controls and realigned variable structure, indicating restoration of integrated neural architecture. Consistently, correlation analyses showed that compensatory contralateral coupling induced by SCI was further strengthened by COLCHI, particularly between somatosensory evoked potentials and motor performance, implicating cortical re-engagement as a key recovery mechanism.

From a clinical perspective, these findings suggest some translational opportunities. First, multidimensional metrics such as PCA and cortical-sensorimotor coupling may serve as sensitive biomarkers for the early identification of treatment responders, while also providing more robust endpoints for clinical trials beyond traditional functional scales. In addition, approaches that integrate multiple types of neurophysiological data and brain network assessment (EEG/fMRI) could offer a more comprehensive way to evaluate the therapeutic efficacy in both preclinical and clinical settings. Second, the enhancement in cortical-sensorimotor coupling observed in the correlation analysis (**Figure 7**) suggest a target for combination treatments where COLCHI scaffolds with motor rehabilitation, brain stimulation (TMS, tDCS), or drugs that promote plasticity (e.g., serotonergic modulators) may enhance recovery by augmenting beneficial changes in brain networks, even in later stages of SCI. Third, strengthened contralateral pathways suggest COLCHI may be particularly effective when paired with bilateral coordination rehabilitation, offering a rational basis for regenerative rehabilitation strategies. Limitations include the inferential nature of covariance analyses (associative, not causal) and unresolved anatomical substrates of circuit nodes (*i.e.,* corticospinal *vs.* propriospinal relays) underlying the observed network shifts. Future work combining longitudinal electrophysiology, pathway-specific circuit dissection, and regenerative rehabilitation principles should validate these mechanistic predictions and build up on the pathway toward first-in-human studies.

## MATERIALS AND METHODS

### Sex as a biological variable

All rats in our study were male, and sex was not considered as a biological variable. It is unknown whether the findings are relevant for female mice.

### NPCHI IONPs and COLCHI hydrogels fabrication and characterization

IONPs with ∼8 nm of core diameter were synthesized via a modified Massart’s co-precipitation method^47^. Briefly, aqueous ferric (0.09 mol) and ferrous (0.054 mol) chloride salts were mixed under stirring, with quick addition of NH₄OH (25 %, 75 mL). The precipitate was washed thrice by magnetic decantation with ultrapure water. For improved colloidal stability, particles were acid-treated with HNO₃ (2M, 300 mL) for 15 min, followed by addition of Fe(NO₃)₃ (1M, 75 mL) and water (130 mL), then heating up to 90 °C for 30 min, and further washing with HNO₃ (2M) and ultrapure water. Resulting IONPs were stored at room temperature in ultrapure water until use. To enhance stability at physiological pH, the resulting bare IONPs were coated with chitosan. Commercial chitosan (Merck, > 75% deacetylated, reference C3646) was dissolved in acetic acid (10 mM) and added dropwise (1_CHI_:5_IONPs_ w/w) to 10 mg of IONPs under acidic conditions. The mixture was incubated for 15 min in sonication and crosslinked with sodium tripolyphosphate (0.5 mg in 0.5 mL water; final volume 10 mL) during 30 min in sonication. IONP core sizes were determined by TEM on a 200 kV JEOL-2000FXII microscope. Sample preparation consisted in depositing dilute aqueous IONP suspensions on carbon-coated copper grids and air-dried prior to imaging, with over 150 particles measured per sample. Colloidal stability was assessed by DLS using a Malvern Zetasizer, recording Z-average hydrodynamic diameters and zeta potentials at pH 7. For thermal analysis, suspensions were freeze-dried at −50 °C and 0.2 mbar for 48 h, then examined by TGA on a Seiko TG/DTA EXSTAR 6000 thermobalance from 25 to 900 °C at 5 °C min⁻¹ under 100 mL min⁻¹ air flow. Magnetic properties were evaluated on lyophilized powders via vibrating sample magnetometry from 5–300 K up to 50 kOe. Magnetic heating capacity of aqueous suspensions was tested in a water-cooled 50 mm coil (Five Celes AC inductor) with temperature monitored by using an optical fiber probe (OSENSA). SAR values were computed as previously reported^48^.

For COLCHI hydrogels preparation, the NPCHI-to-collagen mass ratio was approximately 1:1 as previously reported^49^. Each hydrogel was prepared by pipetting 100 µL of the collagen–IONP solution into 0.5 mL Eppendorf tube caps, rapidly frozen at –80 °C for 5–6 h and lyophilized for 48 h to form randomly porous 3D freeze-casted scaffolds. The so-obtained scaffolds were then crosslinked by exposure to paraformaldehyde (PFA, 4 %) vapors for 2 h under continuous stirring. Morphological characterization was performed using an FEI VERIOS 460 ultrahigh-resolution field-emission microscope and a JEM1400 Flash (Jeol) TEM microscope.

### Neural cell culture studies

Embryonic neural progenitor cells (ENPCs) were isolated from the cerebral cortices of E17 rat embryos following protocols previously established in our laboratory^50^. ENPCs were seeded at a density of 50 x 10^3^ cells/cm^2^ and maintained in complete Neurobasal^TM^ media containing B-27 supplement (2%), streptomycin (100 UI mL^-1^), penicillin (100 UI mL^-1^), and GlutaMAX® (1 %) in a sterile incubator at 37 °C under 5% CO₂, half-replacing media every 3-4 days. At 14 DIV, cells were exposed to IONPs in suspension. After 24 h, cells were gently washed with warm PBS (37 °C) and thereafter incubated in fresh culture media for 24 h before corresponding analysis. Collagen hydrogels were seeded with ENPCs (50 x 10^4^ cells/ scaffold) and maintained for up to 14 DIV. Cell viability in both cultures exposed to NPCHI and collagen hydrogels seeded with ENPCs was determined using the Live/Dead® assay kit according to the manufacturer’s instructions. Samples were incubated with the probes for 5 min at 37 °C before imaging with a Leica SP5 confocal laser scanning microscope. Fluorescence excitation was achieved with a 488 nm Argon laser, and emission was collected using a triple dichroic filter (488/561/633 nm). Green fluorescence from calcein indicated live cells (detected at 505–570 nm), while red fluorescence from ethidium homodimer 1 (EthD-1) marked dead cells (630–750 nm). Viability was quantified by analyzing the fluorescence images with ImageJ software, calculating the percentage area of live (green) and dead (red) cells relative to the total cell area in each image.

### *In vitro* immunofluorescence staining and microscopy imaging of neural cell cultures

For differentiation studies with hybrid hydrogels, cell cultures were fixed with paraformaldehyde (4% in PBS) for 15 min at room temperature (RT), permeabilized with saponin and incubated with primary antibodies as follows: anti-MAP-2 for labelling neurons and anti-vimentin for tagging non-neuronal cells including glial cells. The secondary antibodies used were Alexa Fluor®488 anti-mouse in goat IgG (H + L) and Alexa Fluor®594 anti-rabbit in goat IgG (H + L). Cell nuclei were stained in blue with Hoechst. After staining, samples were visualized under a SP5 confocal laser scanning microscope. Fluorochromes were excited and measured as follows: Alexa Fluor®488 excitation at 488 nm with an argon laser and detection at 507-576 nm, Alexa Fluor®594 excitation at 594 nm with a helium-neon laser and detection at 625-689 nm and Hoechst excitation at 405 nm with a diode UV laser and detection at 423-476 nm.

For quantification of excitatory or inhibitory neurons and synapse quantification in ENPCs exposed to NPCHI, cell cultures were fixed with PFA (4% in PBS, RT, 15 min), then washed 3x with PBS for 5 min and subsequently blocked with PBS solution containing 5% goat serum (Sigma-Aldrich, ref. S26-100ML), 5% donkey serum (Milipore, ref. S30-100ML) and 0.3% Triton X-100 (Sigma-Aldrich, ref. T9284-100ML) for 15 min. Cells were then incubated for 1 h at RT with the following primary antibodies dissolved in the blocking solution: guinea pig monoclonal anti-vGlut1 (1:2500, SYSY, #135304), mouse monoclonal anti-GAD67 (1:1000, Millipore, #MAB5406), and rabbit polyclonal anti-MAP2 (1:500, Termofisher, #PA5-85756). After that, cells were washed 5x in PBS and further incubated for 1 h at RT in darkness with the respective secondary antibodies: goat anti-guinea pig Alexa Fluor 555 (1:500, ABCAM, #AB150186), donkey anti-mouse Alexa Fluor 488 (1:1000, Termofisher, #A-21202), goat anti-rabbit Alexa Fluor 647 (1:1000, ABCAM, #AB150079), goat anti-rabbit Alexa Fluor 488 (1:1000, ABCAM, #AB150077), and goat anti-mouse Alexa Fluor 647 (1:1000, ABCAM, #AB150115). Finally, cells were washed 5x in PBS for 5 min and mounted onto glass slides using Fluoromount medium with DAPI (Sigma-Aldrich, #F6057-20ML). Fluorescence images for VGlut1/GAD67/MAP2 were acquired using an Olympus IX83 epifluorescence microscopy with a 10x objective controlled by CellSens Dimension software. For quantification of the percentage of mature inhibitory and excitatory neurons and their primary branches, three random ∼900 × 900 nm regions were selected in QuPath and transferred to ImageJ. The number of MAP2⁺/VGlut1⁺ and MAP2⁺/GAD67⁺ neurons were then manually counted and the primary branches originating within 50 μm of the soma were quantified. Measurements were performed on two independent coverslips per culture, and averaged values were used for analysis. The mature inhibitory and excitatory neurons in NPCHI conditions were normalized by the control values of the cultures not exposed to IONPs.

For synapse quantification in ENPCs exposed to NPCHI, cell cultures were fixed and immunostained as indicated before using the following antibodies: rabbit monoclonal anti-Homer (1:1000, ABCAM, #AB184955). After incubation, cells were washed as before and incubated with the respective secondary antibodies: goat anti-rabbit Alexa Fluor 488 (1:1000, ABCAM, #AB150077), and goat anti-mouse Alexa Fluor 647 (1:1000, ABCAM, #AB150115). A confocal LEICA SP5 with a 63× oil immersion objective (NA = 1.35) was used to acquire images containing individual neurites with visible synaptic proteins. The number of colocalized synaptic puncta (Homer/Bassoon) was quantified as previously described^51,52^. Briefly, 0.35 µm thick confocal Z-stacks (optical section depth of 0.33 µm; 7 sections/Z-stacks; image area of 2,441 μm^2^) of the neurites were imaged. Maximum projections of 5-7 consecutive optical sections were generated from the original Z-stack. The number of pre-, post-, and colocalized synaptic dots was quantified by using the Puncta Analyser plugin (developed by Barry Wark and provided by Cagla Eroglu) for ImageJ software (RRID:SCR_003070). Images were analyzed by a blind observer. A total of 12 neurons per cell culture were analyzed, averaged and counted as 1. The neurite length used to quantify the synaptic number was measured using the NeuronJ plugin of ImageJ software, and then the number of colocalized synaptic puncta normalized per length (punctas/µm).

### Calcium imaging acquisition and analysis

For calcium imaging, Fluo-4AM (Invitrogen F-14201) was dissolved in dimethyl sulfoxide (DMSO, 2 mM) and 0.01% pluronic acid F-127 (Sigma-Aldrich cat#P2443), which was further diluted in cultured medium to achieve a final concentration of 5 μM. Coverslips containing ENPCs were incubated with the probe at RT for 20–25 min. After that, coverslips were rinsed gently in culture media and incubated with sulforhodamine 101 (SR101; 2 μM) for 5 min to label glial cells. Coverslip were then transferred to an immersion recording chamber and superfused with gassed ACSF (95% O_2_ + 5% CO_2_, 2 mL/min) in an epifluorescence Olympus BX50WI microscope. Ca^2+^ transients were monitored using an CMOS Orca Fusion camera (Hammamatsu) attached to an Olympus BX51WI microscope and illuminated with a CooLED pE-300 fluorescent excitation system. Images were acquired every 200 ms (5 Hz) using the HCImage (Hamamatsu Inc) software under the control of pCLAMP10 (Molecular Devices, Inc). Videos were analyzed offline using SARFIA (freely available on http://www.igorexchange.com/project/SARFIA), a suite of macros and custom-made scripts running in Igor Pro (Wavemetrics, Portland, OR)^53,54^. Prior to analysis, bleaching was corrected using exponential fitting in ImageJ and registered to correct movements in the *X* and *Y* directions if needed. Regions of activity (ROIs) containing both soma and neurites were chosen using a filtering algorithm based on a Laplace operator and segmented by applying a threshold, as described in detail elsewhere. This algorithm defined most or all of the ROIs that an experienced observer would recognize by eye. Individual ROI responses were then normalized as the relative change in fluorescence (*ΔF/F*), smoothed by binomial Gaussian filtering, and analyzed to detect activity using custom-made scripts based on a first derivative detection algorithm. A threshold set at ∼2 times the standard deviation of the time derivative trace was used to detect changes in fluorescence within the ROIs. The reliability of this algorithm to detect calcium activity on ENPCs was first tested by comparing the results with manual activity detection. In this work, the fluorescent intensity of ROIs is reported as the average intensity across all pixels within its area. Fluorescent responses are reported as normalized increases as follows:

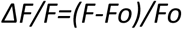

where *F* is the instantaneous fluorescence induced by a spontaneous activity and *Fo* is the baseline fluorescence. Binary raster plots were then constructed from the detected events to analyze event rate. The time points in which calcium transients reach the threshold of the first derivative were also used to overlap individual calcium events used for the kinetics analysis. To study cell network connectivity, we used cross-correlation coefficients obtained by calculating the zero-lag pairwise Pearson correlation coefficients of neuron i and neuron j using the Matlab *corrcoef* routine (MathWorks, Inc). The resulting matrix of cross-correlation coefficients for spontaneous activity was clustered using hierarchical clustering with maximum or complete-linkage clustering. Cross-correlation matrices were plotted without rearranging the neurons and the average of the mean correlation coefficient obtained.

### Spinal cord injury model in rats

Animals were anesthetized with an intraperitoneal (i.p.) injection of sodium pentobarbital (50 mg/kg) mixed with atropine (0.5 mg/kg) and an injection of xylazine (10 mg/kg). If needed, supplementary doses of 30% were administered intramuscularly after 90 min. Eyes were protected with ophthalmologic gel (Lubrithal, Dechra) and body temperature kept constant at a temperature of 36.5 °C using an automatically controlled heating pad (Cibertec SL, Madrid, Spain). Once rats were anesthetized, the skin of the surgical area was shaved and disinfected with povidone iodine. Next, a midline incision was made on the dorsal region from the base of the skull to the T3 backbone level. Then, muscle layers were carefully dissected and retracted, keeping a majority of muscles and main blood vessels intact. Once the vertebral bones were exposed, the lamina of C5 vertebra was removed and a small drop of tetracaine applied on top of the meninges for 2 min to further anesthetize the area. The dura mater was next cut and retracted allowing direct exposure of the C6 spinal cord segment. By using surgical micro-scissors, three cuts (caudal, rostral and sagittal to the midline) were then carried out to achieve a right hemisection of approximately 8 mm^3^. Hemisected animals were randomly distributed into two experimental groups: 1) right hemisection without scaffold (SCI; n = 6), and 2) right hemisection with COLCHI scaffold implantation at the injury site (COLCHI; n = 6). In implanted rats, a cylindrical and porous COLCHI scaffold (2 x 2 x 2 mm^3^) was carefully accommodated inside the injury site. A portion of a microporous patch (Neuro-patch®, Braun) was placed on top to protect the spinal cord. Superficial and deep muscles were then approximated with a bioabsorbable suture, and the skin closed with a similar suture. Immediately after the surgery and during the first three days, animals received injections of meloxicam (5 mg/mL, 0.1 mL/kg every 12 h, subcutaneous; Boehringer Ingelheim), enrofloxacin (2.5%, 25 mg/mL, 0.3 mL/kg every 24 h, subcutaneous; Bayer), and saline solution (5 mL, B. Braun Medical, S.A.). At 120 DPI, animals were sacrificed by using a standard perfusion-fixation protocol. Major organs (spleen, liver, kidneys, and lungs), peripheral muscles (ipsilateral and contralateral brachial and sural triceps) and spinal cords were extracted, weighted and macroscopically examined to discard any signs of gross damage before histological processing. The weight of each main organ and the four groups of peripheral muscles was measured and expressed normalized by the total body weight of each animal.

### Behavioral assessments

Animals underwent longitudinal motor and sensory testing from one week before the injury (pre-SCI) to 120 DPI. Prior testing, all rats were habituated to the experimenter through two daily handling sessions of 5-10 min each for five consecutive days. Habituation was performed in the same room and at the same time of the day as behavioral tests.

#### Ladder walking test

Animals were tested pre-injury and at 30 and 120 DPI, completing 5 trials per session (2 in RP, and 3 in IP). Trials were recorded at 1920 x 1080 pixels, 50 fps, using a ventrally angled camera (Panasonic HC-V785) and analyzed with Kinovea (v0.9.5). The apparatus consisted of two transparent Plexiglas walls (20 x 117 cm^2^) with 47 metal rungs (8 cm), elevated 22 cm above the ground, leading to a home cage. Rungs were arranged either in a regular pattern (RP, 2.5 cm spacing) or in an irregular pattern (IP, 1.25-5 cm spacing). Paw placements were scored using an adapted system from^55^ (**Sup. Figure 2A**). The parameters represented were: a) time to cross, b) errors/total steps, and c) frequency of foot placement (from 0 to 6).

#### Global mobility and spontaneous behavior

Rats were placed in an open field arena consisting of an 80 x 130 cm^2^ table limited by clear Plexiglass walls with 30 cm in height and a non-slippery floor. Behavior was recorded using three digital cameras strategically located around the table to cover 360°. For each session and rat, two videos of 4 min were recorded and used to quantify the relative percentage of active behavior, walking and grooming (stereotyped grooming sequences included forelimb licking and face washing, forelimb face grooming, repetitive body licking, and hind paw scratching^56^.

#### Forelimb asymmetry

To assess forelimb asymmetry, spontaneous exploratory behavior was recorded for over 5 min in their own cage as in^57^. Selected variables were monitored: the number of times that the animal leant on the cage walls by using their right forelimb (ipsilateral forelimb), left forelimb (contralateral) or both (lean on both). When animals were found dramatically inactive, stimuli such as water, paper, food pellets, and tricks (cereal flakes) were used to get them back to an active exploratory behavior. Videos with more than 60 s of animal inactivity were discarded and recorded periods of inactivity longer than 30 s excluded from analyses.

#### Forelimb grooming

Forelimb grooming function was assessed by using a scoring system adapted from previously described protocols^58^. Briefly, tap water was applied to the animal’s head and cervical back area with a soft gauze. Stereotypical grooming sequences of at least 50 s of duration were recorded for each animal in their own cage with a video camera. Forelimb scoring was defined as follows: grooming of the area between the eyes and the ears without reaching ears (position 3), grooming of the ears area (position 4), and grooming of the back area behind the ears (position 5). Slow motion video playback (speed = 0.20x) was used to score each forelimb independently. Values were expressed as a function of time.

#### Grip strength test

Following protocol described in^59^, rats were gently held by the ventral body part and tail and allowed to grasp a horizontal metal bar attached to a force transducer with their forelimbs (Ugo Basile, SRL). Once a firm grip was established, the animal was steadily pulled backward in a horizontal plane until it released the bar. The maximum exerted pulling force was then calculated by the spike amplifier coupled to the transducer. Each rat performed five consecutive trials with an inter-trial interval of 3-min to avoid fatigue. The highest and lowest values were excluded, and the mean of the three intermediate measurements was used for analysis. Animals’ ability to grip the bar with both forelimbs was also analyzed, differentiating between: gripping with both paws (both paws gripping), gripping with the unaffected paw and support with the affected paw (one gripping + one leaning), and grip with only the unaffected paw (one paw gripping).

#### Dynamic Von Frey

Dynamic Plantar Aesthesiometer (Ugo Basile) was used to measure the withdrawal threshold to a single, un-bending filament (0.5 g/sec, 50 seconds) perpendicularly applied to the plantar hindlimb surface^60^. Each animal was tested five times with a 3-min interval between. Trials were repeated if paw withdrawals due to locomotion or weight shifting were observed. Trials in which the animal did not withdraw the paw within 50 seconds were excluded from the analysis. Maximum and minimum readings were rejected for analysis and the three middle measurements were averaged per animal to obtain the mechanical threshold. All animals were acclimatized to the plexiglass chamber with a wire mesh floor for 30 minutes prior to the test.

#### Plantar test

Animals were placed individually in small enclosures with a glass floor (Hargreaves test)^61,62^. An infrared heat source was then focused on the plantar surface of the hindlimb and the time taken to withdraw from the heat stimulus recorded. The intensity of the light source was set to 80 units, corresponding to about 38°C. The maximum duration of the infrared beam was limited to 20 s to prevent burn injuries if the paw was not withdrawn. Each animal was tested 5 times, and the maximum and minimum readings were rejected for analysis.

#### Randall-Selitto assay

Mechanical nociceptive thresholds were assessed using a bench-top Analgesy-Meter (Ugo-Basile)^63^. Rats were gently restrained into a soft cotton cloth, the tested paw held and the medial portion of the plantar hindlimb surface placed into the tip of the device. An increasing mechanical force was then applied until a withdrawal or vocalization response was observed. To avoid skin damage, a maximum force was limited to 250 g. Each hindlimb was tested five times with at least 2-min between trials, and the average threshold after rejecting the maximum and minimum values was calculated for each animal. Animals were acclimated to handling and the testing environment for 5 min prior to measurements to minimize stress-related variability.

### Cranial electrode implant and *in vivo* electrophysiology

Animals were anesthetized by inhalation using a mixture of isoflurane in oxygen (1.5-2%, 2L/ min) and then placed in a stereotaxic frame (SR-6 Narishige Scientific Instruments, Tokio, Japan) with the body temperature kept constant (36.5°C) using an automatically controlled heating pad. Eyes were protected with ophthalmic gel and anesthesia level confirmed by the absence of spinal and corneal reflexes. The skin over the head was shaved and lidocaine (2%) applied subcutaneously to locally anesthetized skin and muscles over the skull. A scalpel was used to perform a longitudinal incision in the midline of the skull and all tissues above the cranial bone were removed. Once the cranial surface was exposed and carefully dried, small holes were drilled bilaterally over CxHL (AP -1.5 mm; ML 2.5 mm) according to Paxinos and Watson (2007). An additional hole was performed located above the cerebellum. Stainless steel screws (Precision Technologies Supplies, M1 X 4 DIN 84 A2; 1mm diameter; 4 mm length) were placed in every location using a mini screwdriver and used as extracranial electrodes to obtain electroencephalography recordings (EEG) from CxHP or reference. After electrodes implantation, anesthesia was adjusted to produce an EEG showing cortical slow-wave activity (SWA), which is characterized by alternating silent and active cortical periods with main frequency <1Hz. Continuous EEG activity from both hemispheres was recorded in DC configuration and filtered up to 3 KHz and amplified (x500). Analog signals were digitized at 20 KHz sampling rate using a CED power 1401 (Cambridge Electronics Design) and amplified by a modular system (Neurolog System; Digitimer Ltd., UK) composed of a preamplifier, filter and amplifier. Signals were converted into digital signals with an A/D converter (1401 CED Cambridge Electronic Designs, UK) and Spike2 software was used for acquisition and posterior analysis.

Electrical stimulation of increasing intensities was applied using bipolar needle electrodes, placed subcutaneously on each side of the hindlimb wrist. Stimulation protocol started only when the EEG remained stable for more than 10 min under SWA. The stimulation protocol (from 0.2, to 8 mA) consisted of 3-min recording periods during which stimulus pulses (1 ms duration) were delivered at a frequency of 0.2 Hz, using a digital stimulator CS-420 (Cibertec S.A.) and an ISO-Flex Flexible Stimulus Isolator (A.M.P.I.). Signals were filtered offline (Spike2 v7, Cambridge Electronics Design, Cambridge, UK), and SEPs analyzed as changes in field potential within the 5–30 ms time window following sensory stimulation. The maximum amplitude (in mV, peak-to-peak) of individual SEPs occurring during down-states was averaged to obtain overall responses. To avoid inter-animal variability, data from individual animals were normalized to its maximum and minimum to obtain an input:output curve. The generated curves were fitted with a Hill-fitting equation to obtain the I_half_ values that were used to measure cortical recruitment. Only animals showing a clear SEP recruitment curve were included in the analysis.

### Histological processing

Spinal cords were placed in PFA 4% (0.1 M in phosphate buffer) at 4 °C for 24 h and organs for one week. All tissue samples were then submersed for 3 days or until sunk in sucrose (30% in 0.1 M phosphate buffer) at 4 °C for cryo-protection. Spinal cord tissue pieces (the entire C5-C7 fragment), mounted on plastic containers with OCT, were cut in sagittal sections of 10-μm from right to left by using a Leica CM1900 cryostat with an angle of 10°. Organs were cut in horizontal sections of 20 µm with the cryostat and stained with conventional hematoxylin/eosin (H&E) stain. Spinal cords were examined after Masson’s trichrome for visualizing collagen. In all cases, panoramic images at low magnification were collected by using an Olympus BX61 microscope. Spinal cord samples were examined for the presence of the following markers: (1) MAP-2 and (2) βIII-tubulin for somas and dendrites in neurons, (3) vimentin for non-neuron cells including glial and connective tissue cells, (4) glial fibrillary acidic protein (GFAP) for astrocytes, (5) ED1 for macrophages, (6) GAP43 for axon growth cones, (7) RECA-1 for endothelial cells in blood vessels, and (8) laminin for basement membranes in blood vessels. Appropriate secondary antibodies were selected accordingly. In all cases, cell nuclei were visualized by labeling with Hoechst (1 mg mL^−1^). Fluorescence images were collected by using a Leica TCS SP5 microscope. Capture conditions were fixed by using sections from the three experimental groups incubated with the secondary antibodies but without the primary ones. All images were thereafter captured under these conditions. All fluorescence images were automatically quantified by using a customized macro in Fiji software as the number of pixels (correspondent μm^2^) positively stained for each particular fluorescence marker after the definition of the correspondent threshold of positive labeling. Control spinal cords and contralateral hemicords in injured animals served as reference values to define correspondent threshold values for each marker. At least three non-overlapping images per animal were acquired in each position of interest (n ≥ 9 per fluorescence marker, study region and group from 6 different animals per group). Areas under study were: LH: left hemicord LH, PL12: perilesional areas at 1–2mm from the lesion site, CIF: caudal interface of the lesion, RIF: rostral interface of the lesion, lesion site without scaffold, and lesion site with scaffold.

### Statistics

Data are presented as the mean ± SEM; box-plot whiskers indicate one standard deviation. For in vitro studies, “N” denotes independent cultures and “n” technical replicates. For in vivo longitudinal studies, values were normalized to the mean of pre-injury baseline. No data was excluded. Normality was assessed with Shapiro-Wilk. Parametric data were analysed using Student’s t-test (paired/unpaired), one-way or two-way ANOVA with Tukey’s post-hoc tests. Non-parametric data used Mann-Whitney or Kruskall-Wallis with Dunn’s correction. Repeated measures (behaviour assessments) were analysed with mixed-effects models with time as a within-subject factor and treatment as a between-subject factor followed by Sidak’s post-hoc test. Geisser–Greenhouse correction was applied when violations of sphericity were found. In all graphs, except when denoted, only the interaction effect was shown for clarity. The rest of statistical values are indicated in figure legends, main text or supplementary tables. Analyses were performed using GraphPad or SPSS (v23.0), with significance set at p < 0.05 (*p < 0.05, **p < 0.01, ***p < 0.001. Principal component analysis (PCA) was used to assess group distribution and variable contribution using behavioural and electrophysiology end-point values (120 DPI). In cases such as grooming or asymmetry tests, only the most significant variable from each test was used (*i.e.* grooming at position 3 and leaning on both forelimbs, respectively). Component number was determined by scree plot and correlation analyses were visualized with heatmaps. Multivariate analyses were conducted in the R/Rstudio (Posit team, 2025; Version 2024.12.1+563). Additional statistical details are provided in supplementary materials.

### Study approval

Experiments were approved by the Ethical Committee for Animal Research at the *Hospital Nacional de Parapléjicos* (Toledo, Spain, 55-OH/2022) and carried out in accordance with the ARRIVE guidelines, the International Council for Laboratory Animal Science and the European Union 2010/63/EU Guidelines. Pregnant adult females and adult male Wistar rats (*ca.* 3 months, 200-300 g) were used in the present study. Pregnant females were used for the collection of E17 embryos (N = 5). Male subjects were selected for the experimental model of SCI to minimize variability arising from hormonal fluctuations associated with the estrous cycle, as the study follows a longitudinal design extending for up to four months post-injury. The total number of male animals (N = 18) was randomly distributed into control (CTR), injured (SCI) and injured receiving COLCHI hydrogel implant (COLCHI). The injury model of selection was a right C6 hemisection. All animals were maintained until 120 DPI, with behavioral tests performed at different time points (basal, 7-10, 14-15, 30, 60, and 120 DPI) to investigate their longitudinal behavioral evolution over time. Animals were group-housed in a controlled temperature environment under a 12 h light/dark cycle, with access to food and water *ad libitum*. All efforts were made to minimize the number of animals used and their suffering and to apply the 3Rs principles.

## Data availability

Data points can be accessed from the Supporting Data Values file. Data for study are available upon request from the corresponding authors.

## Author contributions

EL-D, JA, MCS and JMR designed research study; JM-R fabricated and characterized the biomaterials; ML-A, YH-M, JM-R and EB performed and analysed in vitro data; EL-D, RM-M and MCS performed in vivo spinal cord injuries; VB-M, ML-A, YH-M and CR acquired in vivo behavioral data; VB-M, JM-R, ML-A, YH-M, MS-P, EB, VC and CR performed in vivo behavioral analysis; VB-M and EA-C performed and analysed electrophysiological data; MCS and JMR wrote the manuscript; all authors revised the manuscript; EL-D, JA, MCS and JMR provided fundings. VB-M and JM-R contributed equally to this manuscript and their contribution outstands above that from the rest of co-authors participating in the experimental part of the work. The order between them both was defined based on the impact of the PCA studies carried out by VB-M in the deep discussion of the overall data.

## Funding Support

This work has received funding from the European Union’s Horizon Europe research and Innovation Programme under grant agreement No. 101098597 (Piezo4Spine). It has been also supported by grant PID 2023 - 150170 OB- I 00 (funded by MICIU/AEI/ 10.13039/501100011033 and FEDER, UE), Grant SBPLY/23/180225/000115 (co-funded by “ERDF A way of making Europe”, and JCCM through INNOCAM to JA and JMR).

## Acknowledgements

Schematics representing experimental paradigms or spinal cord injury shown on Figures 2-6 were created with BioRender.com. Animal drawings representing grooming positions shown on panel D, Figure 4, were manually made with a graphic tablet and then texturized using ChatGPT (OpenAI). After the writing, authors used CoPilot (Microsoft, personal account) in Introduction and Discussion sections with the prompt “indicate typos and text redundancy” to improve readability. After using this tool, authors reviewed and edited the content if needed and take full responsibility for the content included in this publication.

The Service of Microscopy and Image Analysis and the Animal Facilities at the *Hospital Nacional de Parapléjicos* are acknowledged for assistance with confocal and rat studies, respectively. Pablo Ruiz-Amezcua and Manuel Nieto-Díaz are acknowledged for technical assistance in the multivariate analysis. Marta Toldos and Andrea Ferreras are acknowledged for technical assistance in early stages of this work. EB acknowledges *Ministerio de Ciencia, Innovación y Universidades* of Spain for an FPI fellowship to grant PID2020-113480RB-I00. ICMM-CSIC acknowledges the Severo Ochoa Centres of Excellence program through Grant CEX2024-001445-S. JMR acknowledges funding from the Spanish Ministry of Science and Innovation MCIN/AEI/10.13039/501100011033 grant RYC2019-026870-I, co-funded by ‘‘ESF Investing in your future’’. MSP acknowledges funding from the Spanish Ministry of Science and Innovation grant JDC2022-049856-I, co-funded by “European Union NextGenerationEU/ PRTR”.

## Notes

**Conflict of Interest** The authors have declared that no conflict of interest exists

### Competing Interest Statement

The authors have declared no competing interest.

