## Supplementary material for "Collagen-based scaffolds loaded with iron oxide nanoparticles promote functional sensorimotor recovery in spinal cord injury": Suplemental material

**1. SUPPLEMENTAL FIGURE LEGENDS**

**2. SUPPLEMENTAL FIGURES**

**3. SUPPLEMENTAL TABLES**

**Sup. Figure 1. COLCHI hydrogels exhibit homogeneous dispersion of IONPs and support the growth of viable and differentiated neural cultures. A)** Representative SEM images. Scale bars: 20 µm (left) and 500 nm (right). **B)** Representative TEM image by cross-section (dark-field mode; IONPs appear as black dots due to higher contrast). Scale bar: 200 nm. **C)** Representative viability images of ENPCs cultured on magnetic hydrogels. Scale bars: 50 (left) and 25 (right) µm. **D)** Representative differentiation images of ENPCs on hybrid hydrogels. Immunolabeling for MAP-2 (green; neurons) and vimentin (red; non-neuronal cells). Scale bar: 150 µm.

**Sup. Figure 2. Step errors in the ladder test. A)** Representative intra-surgical images illustrating the lesion cavity created at C6 (left) and the COLCHI hydrogel implanted (right). Yellow arrow indicates the COLCHI scaffold implanted at the right C6 hemicord. Scale bars: 2 mm. **B)** Experimental details on animal groups, type of paw placement categories and irregular patterns of ladder rungs. **C)** % of error/step obtained from contra/ipsi FL/HL at different time points when using the regular pattern. **D)** % of error/step obtained from contra/ipsi FL/HL at different time points when using the irregular pattern. (n = 6 for SCI, n = 6 COLCHI). Mixed-effects analysis with *post hoc* Tukey's test. * p < 0.05; ** p < 0.01.

**Sup. Figure 3. Ladder scoring in aging/healthy animals. A)** Time to cross the ladder when using regular and irregular patterns at day 0 (white column) and in the same animals at 120 days later (dashed columns). Percentage of error/step and distribution of paw placement categories for both FL and HL when using either regular **(B)** or irregular **(C)** patterns. (n = 6).

**Sup. Figure 4. SCI limits the use of the affected forelimb for grooming tasks. A)** Schematic representation of grooming positions where the first image reflects position 3 (eyes' area), second image is position 4 (ears' area) and third image is position 5 (beyond ears' area). **B)** Normalized grooming assessment with focus on the use of the ipsilateral forelimb to reach positions 3, 4 and 5. (n = 6 per group). Statistics: mixed-effects analysis with Tukey's post-hoc test. * p < 0.05; ** p < 0.01; *** p < 0.001.

**Sup. Figure 5. COLCHI reverses the systemic decoupling induced by SCI. A)** Scree plot of the different principal components to determine the number of components that capture the greatest variance in the data. **B)** Percentage of the contribution that each variable makes to the first 3 main components. **C, D** and **E)** PCA contribution graphs for each group: control (healthy animals), SCI (injured animals without scaffold) and COLCHI (injured animals with COLCHI hydrogel implant).

**Sup. Figure 6. Gross histological examination of the lesion site in COLCHI-implanted paraplegic rats. A)** Masson trichrome staining of the C5-C7 segment in control (untreated) and SCI rats and COLCHI rats **(B)**. Scale bars: 2 mm (left; spinal segments) and 50 µm (spinal tissue zoom-in images, middle and right). Collagen fibers deposited are labelled with gray arrows and macrophage-like cells illustrating the internalization of NPCHI (brownish color) with yellow arrows.

**Sup. Figure 7. Confocal images from immunofluorescence studies of the spinal cord tissue from COLCHI-implanted rats.** Selected markers included neurons (MAP-2 and βIII-tubulin), non-neuronal cells (vimentin), astrocytes (GFAP), macrophages (ED-1), growth cones (GAP43), and vascular structures (RECA-1 and laminin). A) Confocal images from LH: left hemicord and lesion site. B) Confocal images from perilesional area (PL12), caudal interface of the lesion (CIF) and rostral interface of the lesion (RIF).

**Sup. Figure 8. Systemic toxicity at pivotal organs in COLCHI-implanted paraplegic rats**. **A)** Representative hematoxylin/eosin microscopy images for the spleen, liver, kidney, and lung. **B)** Weight percentage with respect to total body weight for each specific organ. **C)** Weight percentage (top) and volume (bottom) of peripheral limb muscles including the ipsilateral and contralateral brachial triceps (RBT and LBT, respectively) and the ipsilateral and contralateral sural triceps (RST and LST, respectively). Treatment groups: control (healthy rats; white), SCI (injured rats without COLCHI scaffold; dark purple) and COLCHI (injured rats with COLCHI scaffold; pale orange). Statistics: one-way ANOVA followed by corresponding *post hoc* tests. Statistical significance: $p < 0.05$* and $p < 0.01$**).

Supplemental Figure 1

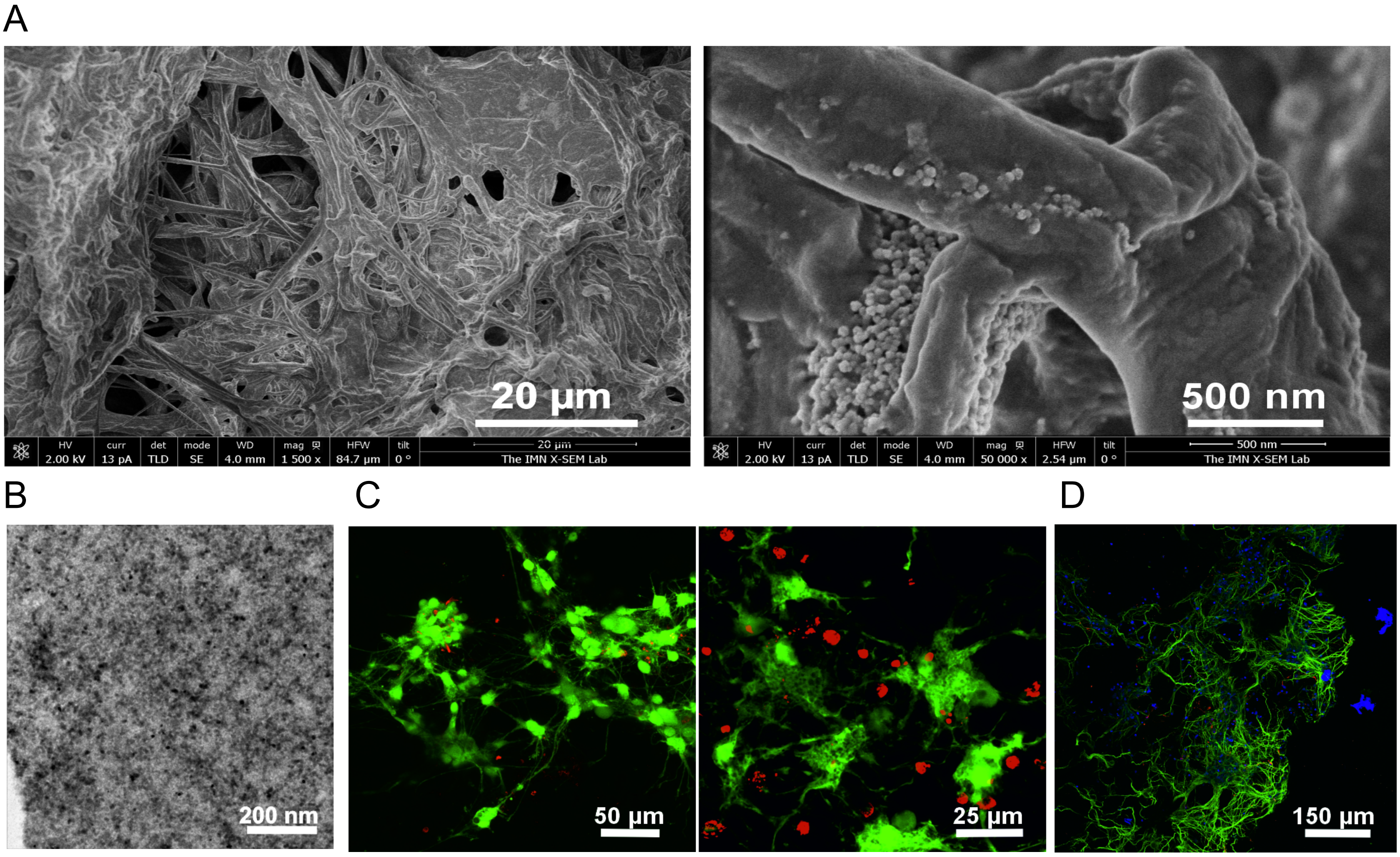

Supplemental Figure 2

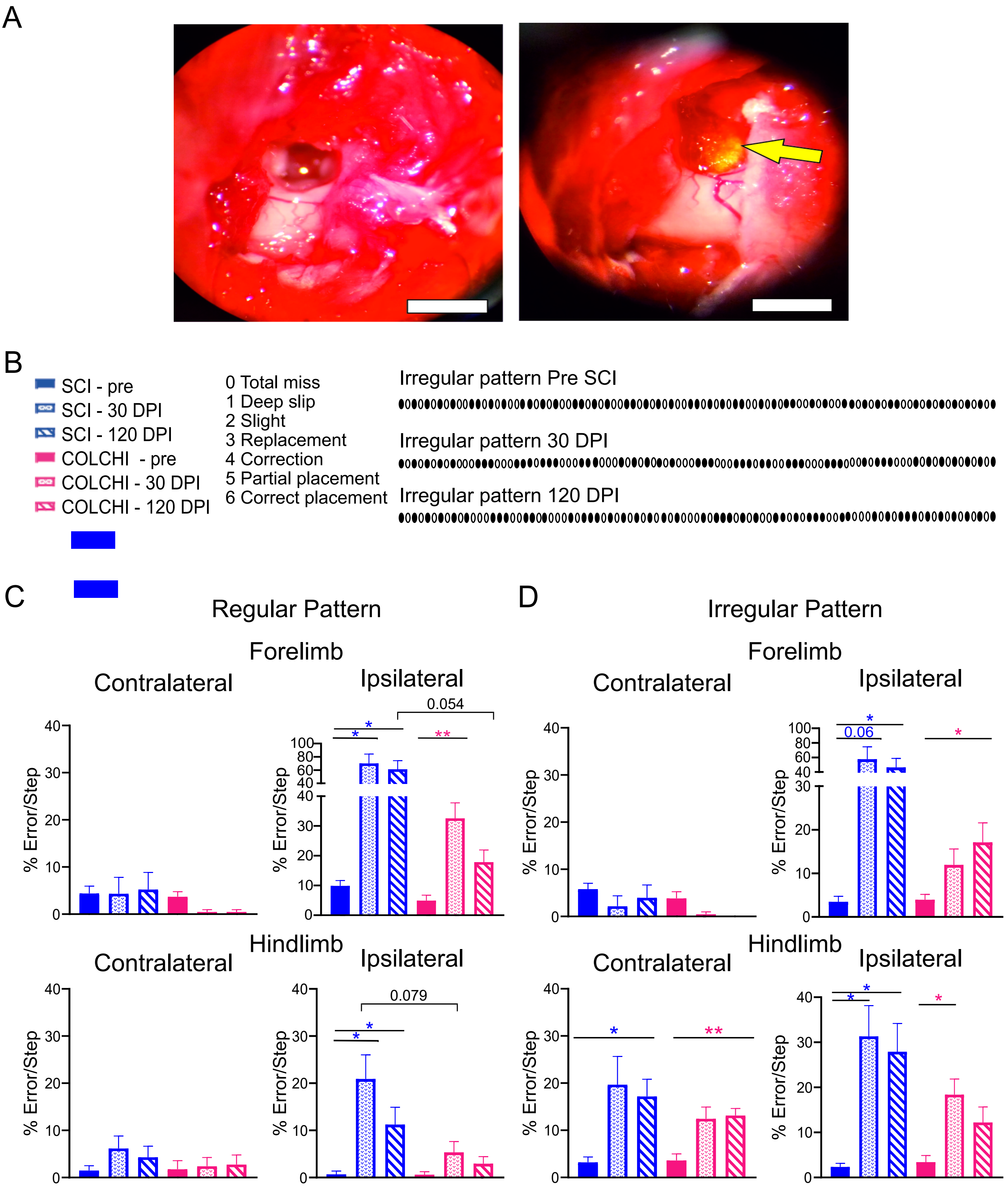

Supplemental Figure 3

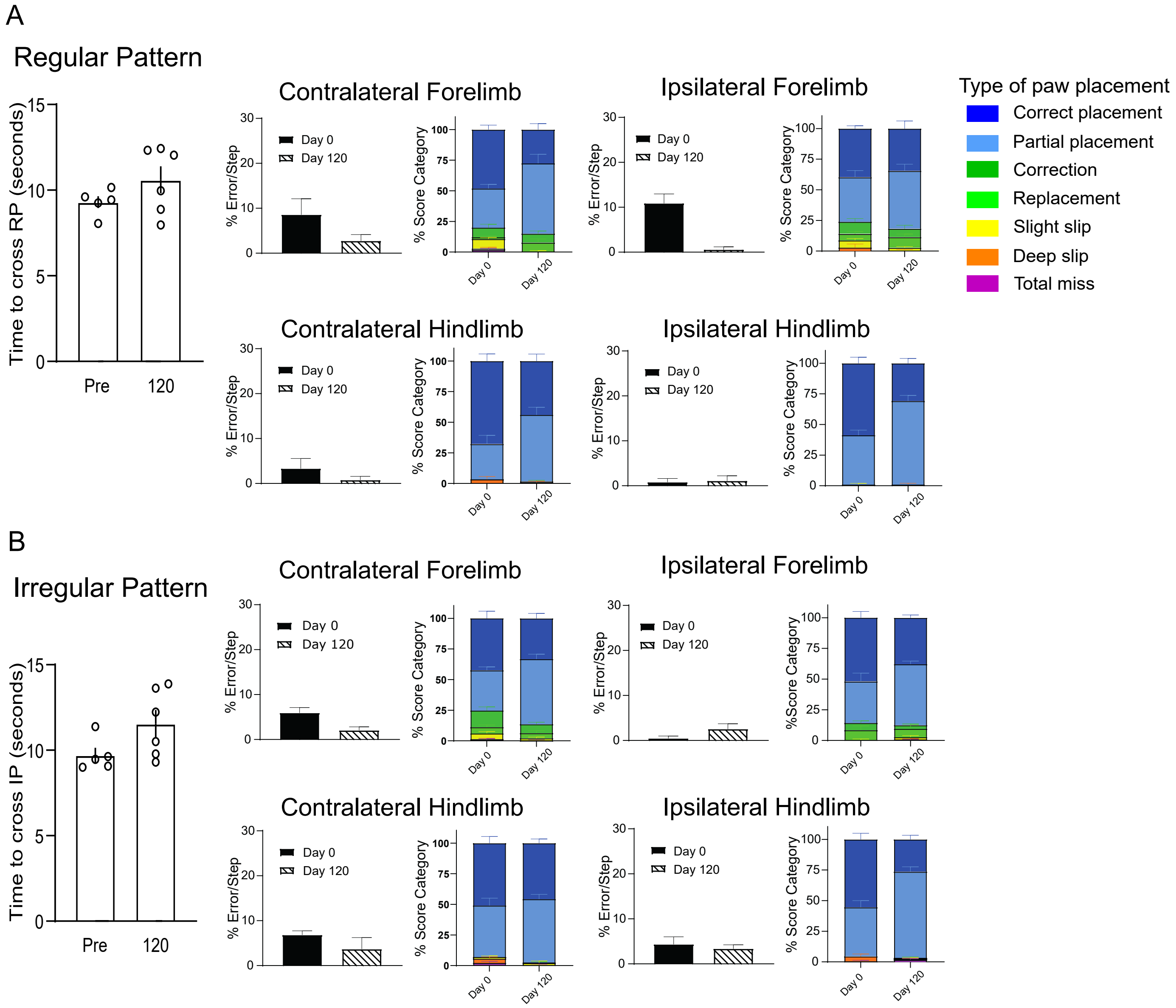

Supplemental Figure 4

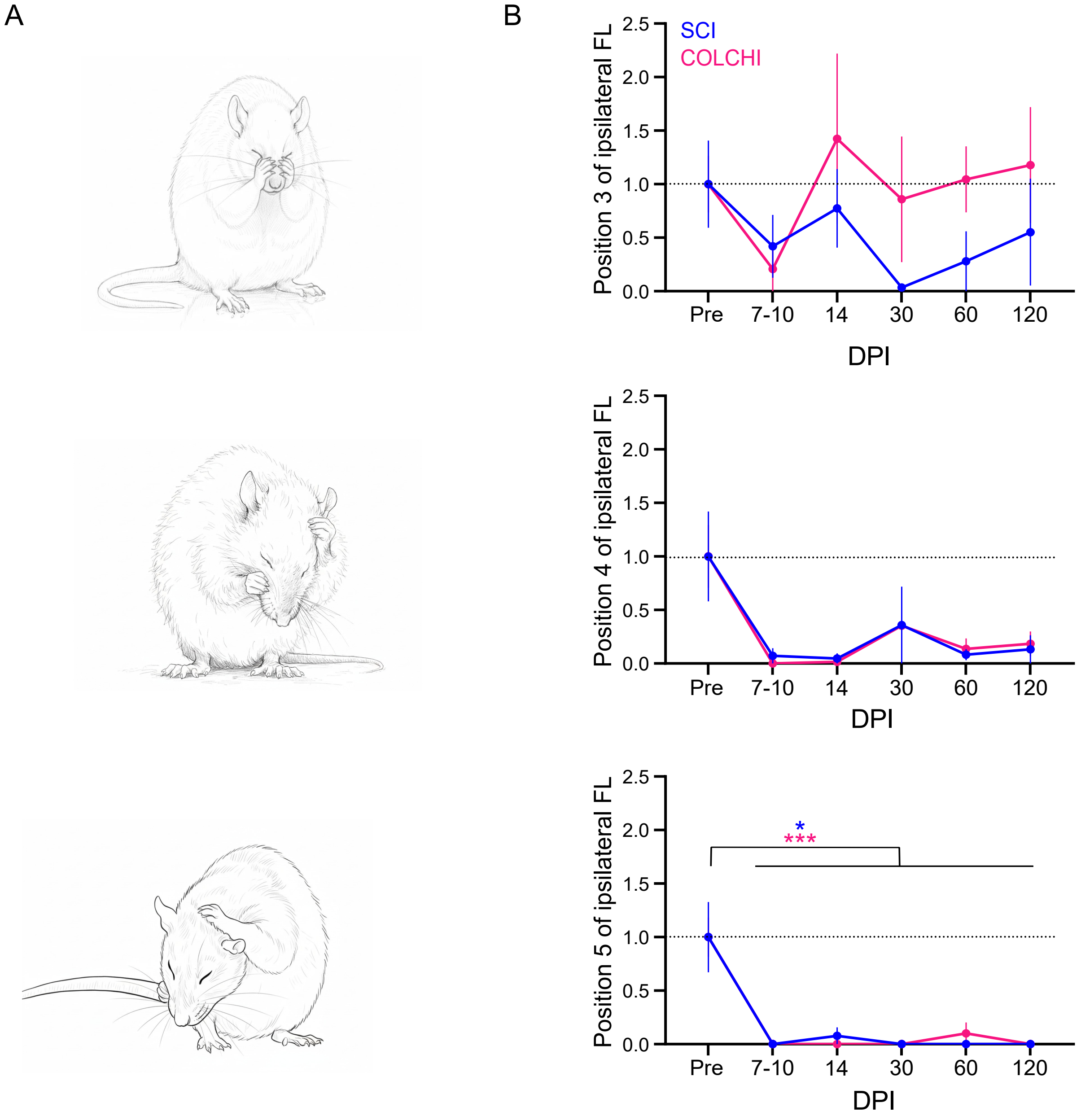

### Supplemental Figure 5

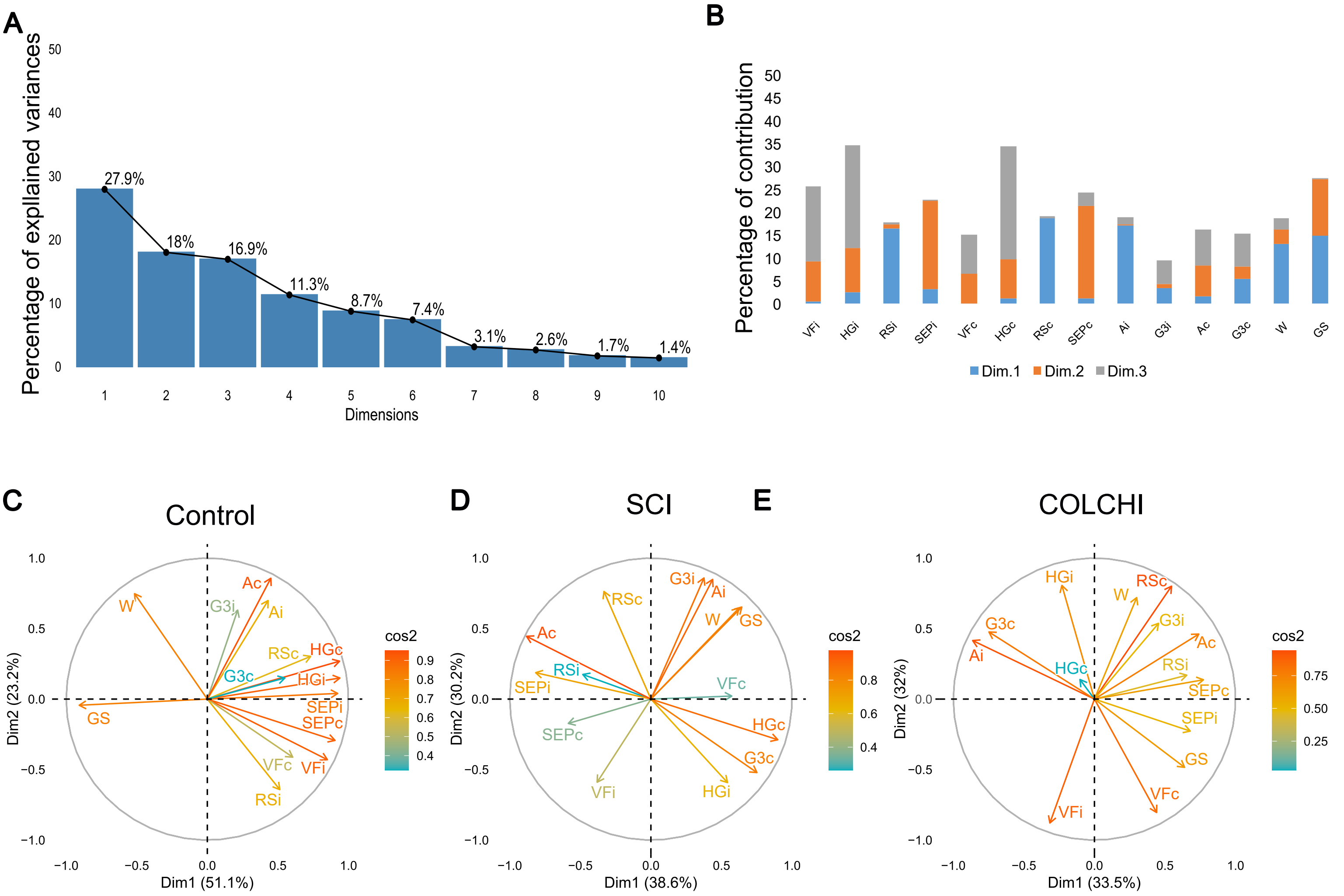

Supplemental Figure 6

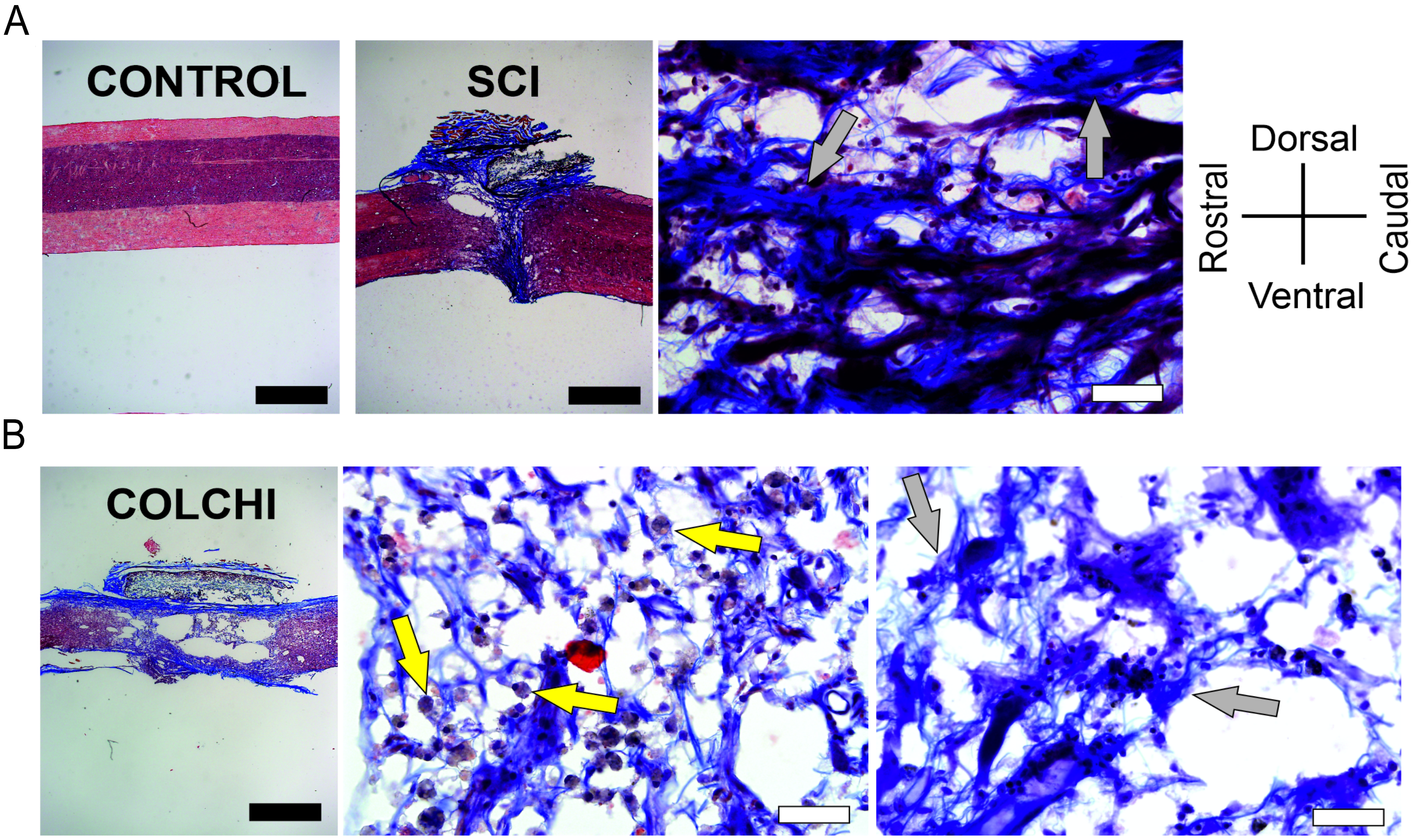

Supplemental figure 7

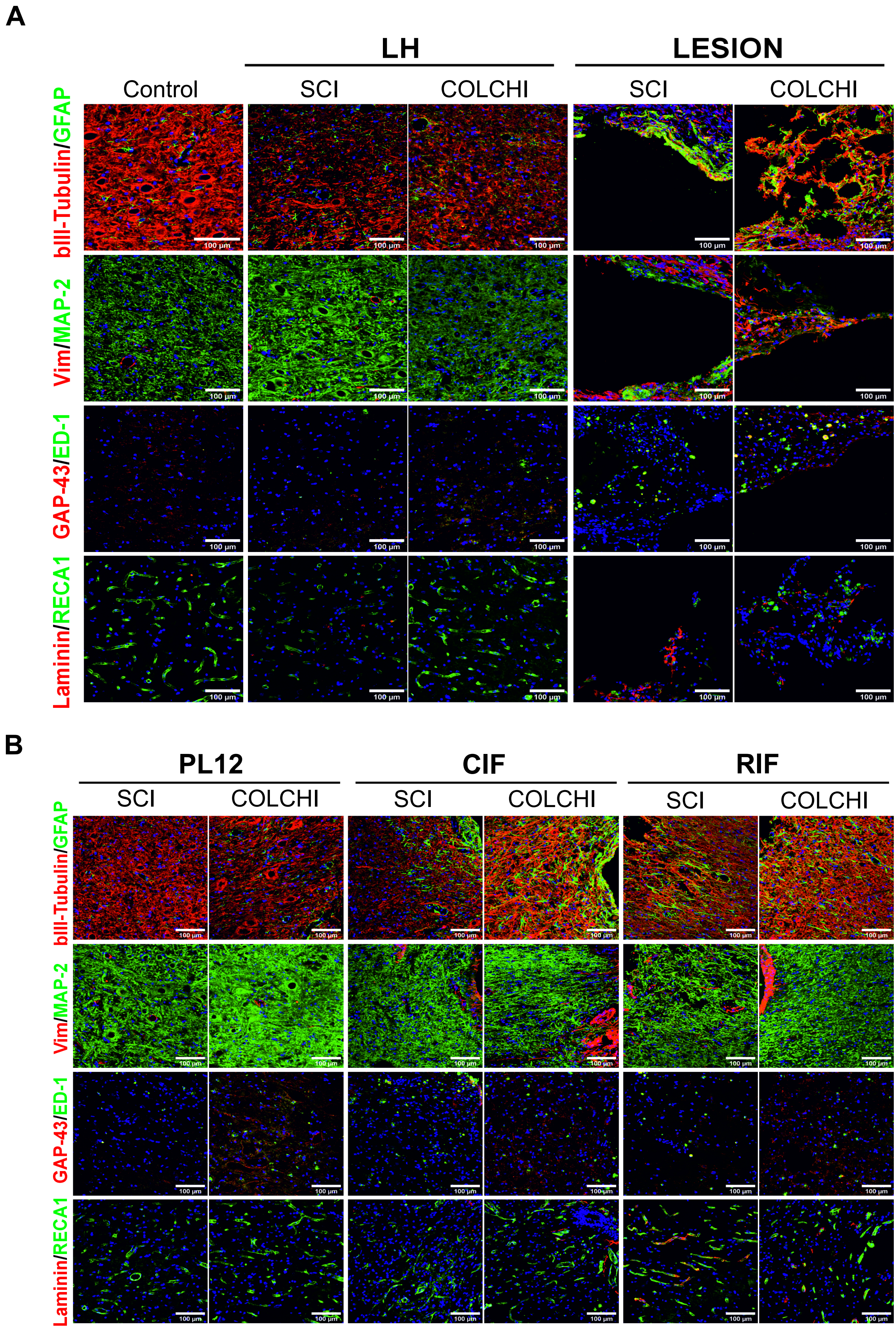

Supplemental Figure 8

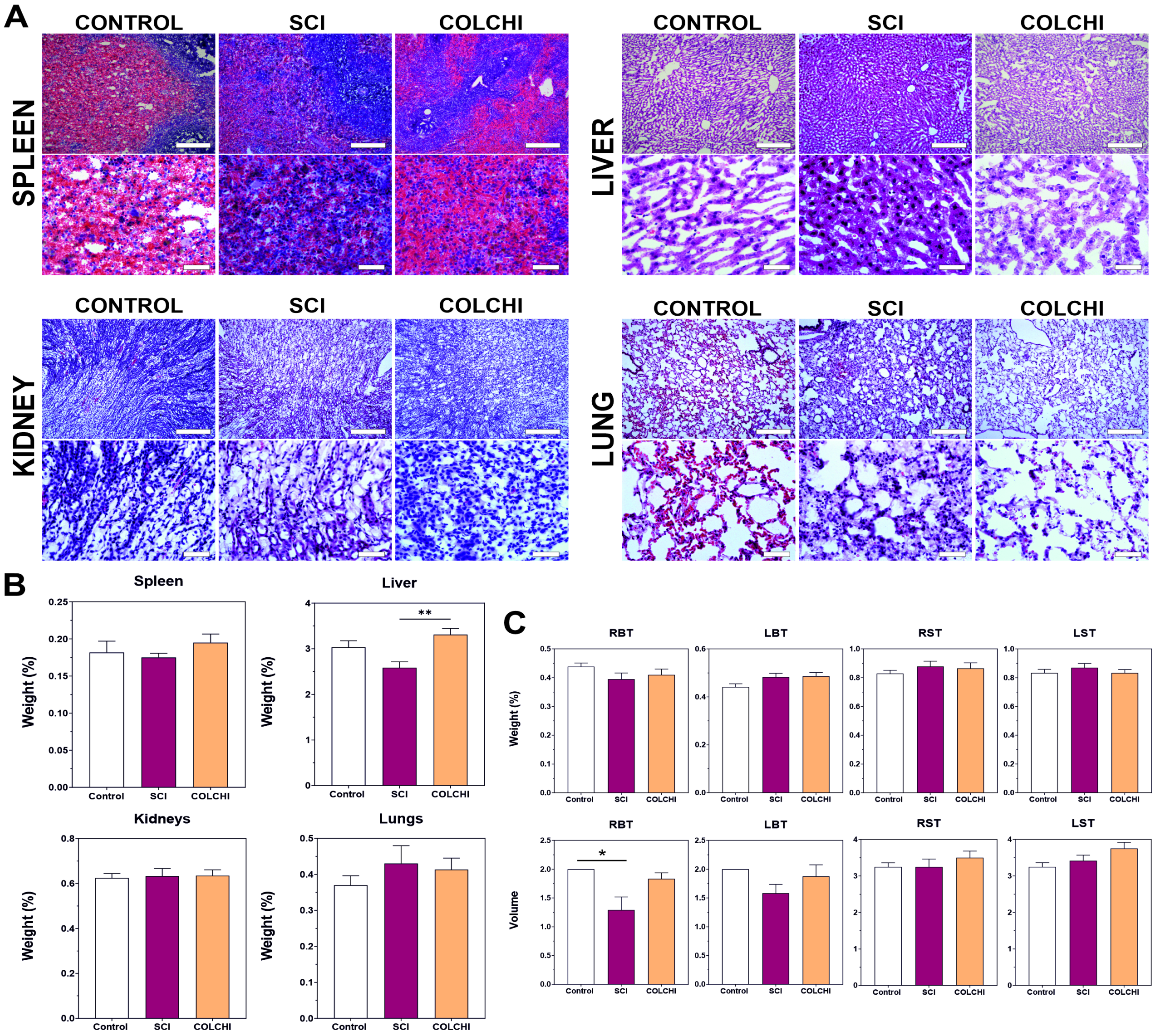

**Supplemental Tables**

**Supplemental Table 1: Statistics for Fig. 3 C-H: Time to cross and % Score Category**

| **Figure 3C: Time to cross of Regular pattern** | | |
|---|---|---|
| **Kruskal-Wallis test** | | |
| **SCI** | **COLCHI** | |
| **p-value** | **p-value** | |
| 0.0007 | 0.3527 | |
| **Dunn's *post hoc* (SCI)** | | |
| | **Mean rank diff** | **p-value** |
| Pre vs 30 | -9.833 | 0.0028 |
| Pre vs 120 | -7.667 | 0.0257 |

| **Figure 3D: %Score category of contralateral forelimb (Regular pattern)** | | | | |
|---|---|---|---|---|
| **Total miss** | **Mixed-effects model (REML)** | | | |
| | | **F (DFn, DFd)** | **p-value** | **Geisser-Greenhouse's ε** |
| | Time | F (1.321, 19.81) = 0.5525 | 0.5133 | 0.6604 |
| | Column Factor | F (1, 30) = 1.895 | 0.1788 | |
| | Time x Column Factor | F (2, 30) = 0.5525 | 0.5812 | |
| **Deep slip** | **Mixed-effects model (REML)** | | | |
| | | **F (DFn, DFd)** | **p-value** | **Geisser-Greenhouse's ε** |
| | Time | F (2, 30) = 1.008 | 0.3384 | 0.5505 |
| | Column Factor | F (1, 30) = 0.7992 | 0.3784 | |
| | Time x Column Factor | F (2, 30) = 2.388 | 0.1091 | |
| **Slight slip** | **Mixed-effects model (REML)** | | | |
| | | **F (DFn, DFd)** | **p-value** | **Geisser-Greenhouse's ε** |
| | Time | F (1.433, 14.33) = 1.193 | 0.3149 | 0.7163 |

| | | | | | | |
|---|---|---|---|---|---|---|
| | Column Factor | F (1, 10) = 1.872 | | 0.2012 | | |
| | Time x Column Factor | F (2, 20) = 0.4374 | | 0.6517 | | |
| **Replacement** | **Mixed-effects model (REML)** | | | | | |
| | | **F (DFn, DFd)** | | **p-value** | **Geisser-Greenhouse's ε** | |
| | Time | F (1.471, 22.06) = 3.026 | | 0.0821 | 0.7355 | |
| | Column Factor | F (1, 30) = 5.586 | | 0.0248 | | |
| | Time x Column Factor | F (2, 30) = 1.912 | | 0.1654 | | |
| **Correction** | **Mixed-effects model (REML)** | | | | | |
| | | **F (DFn, DFd)** | | **p-value** | **Geisser-Greenhouse's ε** | |
| | Time | F (1.324, 19.86) = 0.2170 | | 0.7141 | 0.662 | |
| | Column Factor | F (1, 30) = 1.588 | | 0.2174 | | |
| | Time x Column Factor | F (2, 30) = 2.314 | | 0.1163 | | |
| **Partial placement** | **Mixed-effects model (REML)** | | | | | |
| | | **F (DFn, DFd)** | | **p-value** | **Geisser-Greenhouse's ε** | |
| | Time | F (1.633, 24.50) = 0.05454 | | 0.9181 | 0.8167 | |
| | Column Factor | F (1, 30) = 3.033 | | 0.0919 | | |
| | Time x Column Factor | F (2, 30) = 0.5896 | | 0.5608 | | |
| **Correct placement** | **Two-way ANOVA** | | | | | |
| | | **SS** | **DF** | **MS** | **F (DFn, DFd)** | **p-value** |
| | Interaction | 91.31 | 2 | 45.66 | F (2, 20) = 0.2050 | 0.8163 |
| | Time | 243.7 | 2 | 121.8 | F (2, 20) = 0.5470 | 0.5871 |
| | Column Factor | 132.4 | 1 | 132.4 | F (1, 10) = 0.2753 | 0.6112 |

| | | | | | | |
|---|---|---|---|---|---|---|
| | Subject | 4811 | 10 | 481.1 | F (10, 20) = 2.160 | 0.0685 |
| | Residual | 4454 | 20 | 222.7 | | |

| **Figure 3D: %Score category of ipsilateral forelimb (Regular pattern)** | | | | | |
|---|---|---|---|---|---|
| **Total miss** | **Mixed-effects model (REML)** | | | | |
| | | **F (DFn, DFd)** | **p-value** | **Geisser-Greenhouse's ε** | |
| | Time | F (1.313, 13.13) = 19.52 | 0.0003 | 0.6565 | |
| | Column Factor | F (1, 10) = 6.503 | 0.0289 | | |
| | Time x Column Factor | F (2, 20) = 5.105 | 0.0162 | | |
| | **Sidak's *post hoc* (SCI-COLCHI) (SCI-COLCHI)** | | | | |
| | | **Mean diff,** | **95,00% CI of diff,** | **p-value** | |
| | Pre | 0.641 | -1.614 to 2.896 | 0.7418 | |
| | 30 | 31.49 | -18.05 to 81.03 | 0.2247 | |
| | 120 | 38.02 | -8.731 to 84.76 | 0.104 | |
| | **Tukey's *post hoc*** | | | | |
| | | | **Mean diff,** | **95,00% CI of diff,** | **p-value** |
| | **SCI** | Pre vs. 30 | -53.38 | -101.1 to -5.710 | 0.0333 |
| | | Pre vs. 120 | -45.55 | -90.03 to -1.068 | 0.046 |
| | | 30 vs. 120 | 7.831 | -5.284 to 20.95 | 0.2214 |
| | **COLCHI** | Pre vs. 30 | -22.53 | -35.79 to -9.276 | 0.0061 |
| | | Pre vs. 120 | -8.174 | -15.28 to -1.069 | 0.0301 |
| | | 30 vs. 120 | 14.36 | -3.250 to 31.96 | 0.0972 |
| **Deep slip** | **Mixed-effects model (REML)** | | | | |
| | | **F (DFn, DFd)** | **p-value** | **Geisser-Greenhouse's ε** | |
| | Time | F (1.761, 17.61) = 1.508 | 0.2476 | 0.8807 | |
| | Column Factor | F (1, 10) = 2,984 | 0.1148 | | |
| | Time x Column Factor | F (2, 20) = 1.050 | 0.3686 | | |

| | | | | | |
|---|---|---|---|---|---|
| **Slight slip** | **Mixed-effects model (REML)** | | | | |
| | | **F (DFn, DFd)** | **p-value** | **Geisser-Greenhouse's ε** | |
| | Time | F (1.415, 14.15) = 1.178 | 0.318 | 0.7077 | |
| | Column Factor | F (1, 10) = 0.1218 | 0.7344 | | |
| | Time x Column Factor | F (2, 20) = 0.4684 | 0.6327 | | |
| **Replacement** | **Mixed-effects model (REML)** | | | | |
| | | **F (DFn, DFd)** | **p-value** | **Geisser-Greenhouse's ε** | |
| | Time | F (1.498, 14.98) = 3.267 | 0.0775 | 0.749 | |
| | Column Factor | F (1, 10) = 1.103 | 0.3183 | | |
| | Time x Column Factor | F (2, 20) = 1.610 | 0.2247 | | |
| **Correction** | **Mixed-effects model (REML)** | | | | |
| | | **F (DFn, DFd)** | **p-value** | **Geisser-Greenhouse's ε** | |
| | Time | F (1.704, 25.55) = 15.33 | <0.0001 | 0.8518 | |
| | Column Factor | F (1, 30) = 3.903 | 0.0575 | | |
| | Time x Column Factor | F (2, 30) = 0.8143 | 0.4525 | | |
| | **Tukey's *post hoc*** | | | | |
| | | | **Mean diff,** | **95,00% CI of diff,** | **p-value** |
| | **SCI** | Pre vs. 30 | 8.027 | -0.6078 to 16.66 | 0.0641 |
| | | Pre vs. 120 | 7.945 | 0.5463 to 15.34 | 0.0388 |
| | | 30 vs. 120 | -0.08167 | -5.130 to 4.966 | 0.9985 |
| | **COLCHI** | Pre vs. 30 | 10.24 | 1.455 to 19.02 | 0.0286 |
| | | Pre vs. 120 | 5.774 | -6.608 to 18.16 | 0.3595 |
| | | 30 vs. 120 | -4.465 | -12.97 to 4.043 | 0.2904 |
| [illegible] | **Two-way ANOVA** | | | | |

**Partial placement**

| | SS | DF | MS | F (DFn, DFd) | p-value |
|---|---|---|---|---|---|
| Interaction | 692 | 2 | 346 | F (2, 20) = 4.493 | 0.0245 |
| Time | 326.2 | 2 | 163.1 | F (2, 20) = 2.118 | 0.1464 |
| Column Factor | 973.7 | 1 | 973.7 | F (1, 10) = 2.444 | 0.1491 |
| Subject | 3984 | 10 | 398.4 | F (10, 20) = 5.174 | 0.0009 |
| Residual | 1540 | 20 | 77.01 | | |

**Sidak's *post hoc* (SCI-COLCHI) (SCI-COLCHI)**

| | Mean diff, | 95,00% CI of diff, | p-value |
|---|---|---|---|
| Pre | 1.679 | -18.13 to 21.49 | 0.9952 |
| 30 | -18.87 | -38.68 to 0.9435 | 0.0657 |
| 120 | -14.02 | -33.83 to 5.793 | 0.2307 |

**Tukey's *post hoc***

| | | Mean diff, | 95,00% CI of diff, | p-value |
|---|---|---|---|---|
| **SCI** | Pre vs. 30 | 16.61 | 8.465 to 24.75 | 0.0027 |
| | Pre vs. 120 | 7.748 | -6.949 to 22.45 | 0.2878 |
| | 30 vs. 120 | -8.86 | -19.63 to 1.911 | 0.0947 |
| **COLCHI** | Pre vs. 30 | -3.937 | -21.21 to 13.33 | 0.7513 |
| | Pre vs. 120 | -7.947 | -32.56 to 16.67 | 0.5808 |
| | 30 vs. 120 | -4.01 | -22.13 to 14.11 | 0.7631 |

**Correct placement**

**Mixed-effects model (REML)**

| | F (DFn, DFd) | p-value | Geisser-Greenhouse's ε |
|---|---|---|---|
| Time | F (1.032, 10.32) = 13.98 | 0.0035 | 0.5161 |
| Column Factor | F (1, 10) = 4.774 | 0.0538 | |
| Time x Column Factor | F (2, 20) = 3.018 | 0.0715 | |

**Tukey's *post hoc***

| | | Mean diff, | 95,00% CI of diff, | p-value |
|---|---|---|---|---|
| **SCI** | Pre vs. 30 | 33.77 | -3.637 to 71.17 | 0.0706 |

| | | | | | |
|---|---|---|---|---|---|
| | | Pre vs. 120 | 34.31 | -1.610 to 70.24 | 0.0585 |
| | | 30 vs. 120 | 0.5492 | -0.9966 to 2.095 | 0.5252 |
| | **COLCHI** | Pre vs. 30 | 15.71 | -0.6441 to 32.07 | 0.0574 |
| | | Pre vs. 120 | 9.973 | -9.207 to 29.15 | 0.2956 |
| | | 30 vs. 120 | -5.74 | -11.50 to 0.01921 | 0.0506 |

| **Figure 3E: %Score category of contralateral hindlimb (Regular pattern)** | | | | |
|---|---|---|---|---|
| **Total miss** | **Mixed-effects model (REML)** | | | |
| | | **F (DFn, DFd)** | **p-value** | **Geisser-Greenhouse's ε** |
| | Time | F (1.000, 10.00) = 1.000 | 0.3409 | 0.5 |
| | Column Factor | F (1, 10) = 1.000 | 0.3409 | |
| | Time x Column Factor | F (2, 20) = 1.000 | 0.3855 | |
| **Deep slip** | **Mixed-effects model (REML)** | | | |
| | | **F (DFn, DFd)** | **p-value** | **Geisser-Greenhouse's ε** |
| | Time | F (1.582, 15.82) = 0.3491 | 0.6615 | 0.7909 |
| | Column Factor | F (1, 10) = 0.04948 | 0.8285 | |
| | Time x Column Factor | F (2, 20) = 0.7466 | 0.4867 | |
| **Slight slip** | **Mixed-effects model (REML)** | | | |
| | | **F (DFn, DFd)** | **p-value** | **Geisser-Greenhouse's ε** |
| | Time | F (1.713, 17.13) = 0.3686 | 0.6652 | 0.8567 |
| | Column Factor | F (1, 10) = 0.8651 | 0.3742 | |
| | Time x Column Factor | F (2, 20) = 0.8712 | 0.4338 | |
| **Replacement** | **Mixed-effects model (REML)** | | | |
| | | **F (DFn, DFd)** | **p-value** | **Geisser-Greenhouse's ε** |
| | Time | F (1.000, 15.00) = 4.746 | 0.0457 | 0.5 |

| | | | | |
|---|---|---|---|---|
| | Column Factor | F (1, 30) = 4.746 | 0.0374 | |
| | Time x Column Factor | F (2, 30) = 4.746 | 0.0162 | |
| | **Sidak's *post hoc* (SCI-COLCHI) (SCI-COLCHI)** | | | |
| | | **Mean diff,** | **95,00% CI of diff,** | **p-value** |
| | 30 dpi | 2.083 | -0.3750 to 4.542 | 0.0813 |
| | **Tukey's *post hoc*** | | | |
| | | | **Mean diff,** | **95,00% CI of diff,** | **p-value** |
| | **SCI** | Pre vs. 30 | -2.083 | -4.542 to 0.3750 | 0.0813 |
| | | 30 vs. 120 | 2.083 | -0.3750 to 4.542 | 0.0813 |
| **Correction** | **Mixed-effects model (REML)** | | | |
| | | **F (DFn, DFd)** | **p-value** | **Geisser-Greenhouse's ε** |
| | Time | F (1.583, 23.74) = 0.5045 | 0.5676 | 0.7914 |
| | Column Factor | F (1, 30) = 0.009061 | 0.9248 | |
| | Time x Column Factor | F (2, 30) = 1.495 | 0.2404 | |
| **Partial placement** | **Mixed-effects model (REML)** | | | |
| | | **F (DFn, DFd)** | **p-value** | **Geisser-Greenhouse's ε** |
| | Time | F (1.769, 17.69) = 13.30 | 0.0004 | 0.8844 |
| | Column Factor | F (1, 10) = 0.2310 | 0.6411 | |
| | Time x Column Factor | F (2, 20) = 3.921 | 0.0366 | |
| | **Sidak's *post hoc* (SCI-COLCHI)** | | | |
| | | **Mean diff,** | **95,00% CI of diff,** | **p-value** |
| | Pre | -10.84 | -33.70 to 12.02 | 0.4956 |
| | 30 | 2.65 | -22.27 to 27.57 | 0.9874 |
| | 120 | 15.53 | 0.2495 to 30.82 | 0.0463 |
| | **Tukey's *post hoc*** | | | |

| | | | | |
|---|---|---|---|---|
| | | Mean diff, | 95,00% CI of diff, | p-value |
| **SCI** | Pre vs. 30 | -25.01 | -41.46 to -8.565 | 0.0099 |
| | Pre vs. 120 | -36.18 | -56.35 to -16.00 | 0.0049 |
| | 30 vs. 120 | -11.17 | -34.15 to 11.81 | 0.335 |
| **COLCHI** | Pre vs. 30 | -11.52 | -40.74 to 17.69 | 0.4618 |
| | Pre vs. 120 | -9.807 | -36.00 to 16.39 | 0.4937 |
| | 30 vs. 120 | 1.717 | -6.764 to 10.20 | 0.7959 |

**Correct placement**

**Two-way ANOVA**

| | SS | DF | MS | F (DFn, DFd) | p-value |
|---|---|---|---|---|---|
| Interaction | 1188 | 2 | 594.2 | F (2, 20) = 6.078 | 0.0087 |
| Time | 4560 | 2 | 2280 | F (2, 20) = 23.32 | <0.0001 |
| Column Factor | 209.2 | 1 | 209.2 | F (1, 10) = 0.7193 | 0.4162 |
| Subject | 2909 | 10 | 290.9 | F (10, 20) = 2.975 | 0.0182 |
| Residual | 1955 | 20 | 97.77 | | 0.0087 |

**Sidak's *post hoc* (SCI-COLCHI)**

| | Mean diff, | 95,00% CI of diff, | p-value |
|---|---|---|---|
| Pre | 11 | -7.585 to 29.59 | 0.3748 |
| 30 | -9.532 | -28.12 to 9.056 | 0.4969 |
| 120 | -15.94 | -34.52 to 2.652 | 0.1104 |

**Tukey's *post hoc***

| | | Mean diff, | 95,00% CI of diff, | p-value |
|---|---|---|---|---|
| **SCI** | Pre vs. 30 | 32.18 | 17.73 to 46.62 | <0.0001 |
| | Pre vs. 120 | 38.91 | 24.47 to 53.36 | <0.0001 |
| | 30 vs. 120 | 6.737 | -7.705 to 21.18 | 0.4782 |
| **COLCHI** | Pre vs. 30 | 11.64 | -2.800 to 26.09 | 0.1286 |
| | Pre vs. 120 | 11.98 | -2.466 to 26.42 | 0.1156 |
| | 30 vs. 120 | 0.3337 | -14.11 to 14.78 | 0.9981 |

**Figure 3E: %Score category of ipsilateral hindlimb (Regular pattern)**

**Total miss**

**Mixed-effects model (REML)**

| | F (DFn, DFd) | p-value | Geisser-Greenhouse's ε |
|---|---|---|---|

| | | | | | |
|---|---|---|---|---|---|
| | Time | | F (1.665, 16.65) = 5.645 | 0.0171 | 0.8326 |
| | Column Factor | | F (1, 10) = 1.221 | 0.295 | |
| | Time x Column Factor | | F (2, 20) = 1.062 | 0.3646 | |
| | **Tukey's *post hoc*** | | | | |
| | | | **Mean diff,** | **95,00% CI of diff,** | **p-value** |
| | **SCI** | Pre vs. 30 | -5.483 | -12.96 to 1.993 | 0.1321 |
| | | Pre vs. 120 | -2.543 | -8.384 to 3.297 | 0.4012 |
| | | 30 vs. 120 | 2.94 | -2.965 to 8.845 | 0.3206 |
| | **COLCHI** | Pre vs. 30 | -2.165 | -7.030 to 2.700 | 0.3879 |
| | | Pre vs. 120 | -0.9804 | -4.171 to 2.210 | 0.6083 |
| | | 30 vs. 120 | 1.185 | -1.308 to 3.677 | 0.3481 |
| **Deep slip** | **Mixed-effects model (REML)** | | | | |
| | | | **F (DFn, DFd)** | **p-value** | **Geisser-Greenhouse's ε** |
| | Time | | F (1.191, 10.72) = 0.8674 | 0.3919 | 0.5957 |
| | Column Factor | | F (1, 10) = 3.093 | 0.1091 | |
| | Time x Column Factor | | F (2, 18) = 0.2379 | 0.7907 | |
| **Slight slip** | **Mixed-effects model (REML)** | | | | |
| | | | **F (DFn, DFd)** | **p-value** | **Geisser-Greenhouse's ε** |
| | Time | | F (1.322, 19.84) = 6.729 | 0.0118 | 0.6612 |
| | Column Factor | | F (1, 30) = 15.05 | 0.0005 | |
| | Time x Column Factor | | F (2, 30) = 4.534 | 0.019 | |
| | **Sidak's *post hoc* (SCI-COLCHI)** | | | | |
| | | | **Mean diff,** | **95,00% CI of diff,** | **p-value** |
| | Pre | | - | - | - |
| | 30 | | 6.777 | 1.164 to 12.39 | 0.0256 |

| | | | | |
|---|---|---|---|---|
| | 120 | 3.968 | 0.04396 to 7.893 | 0.0483 |
| | **Tukey's *post hoc*** | | | |
| | | **Mean diff,** | **95,00% CI of diff,** | **p-value** |
| | **SCI** Pre vs. 30 | -7.535 | -14.64 to -0.4320 | 0.0406 |
| | Pre vs. 120 | -3.968 | -8.936 to 0.9992 | 0.1034 |
| | 30 vs. 120 | 3.567 | -6.064 to 13.20 | 0.5002 |
| | **COLCHI** Pre vs. 30 | -0.7583 | -2.708 to 1.191 | 0.3632 |
| | Pre vs. 120 | - | - | - |
| | 30 vs. 120 | 0.7583 | -1.191 to 2.708 | 0.3632 |
| **Replacement** | **Mixed-effects model (REML)** | | | |
| | | **F (DFn, DFd)** | **p-value** | **Geisser-Greenhouse's ε** |
| | Time | F (1.000, 15.00) = 4.542 | 0.05 | 0.5 |
| | Column Factor | F (1, 30) = 4.542 | 0.0414 | |
| | Time x Column Factor | F (2, 30) = 4.542 | 0.0189 | |
| | **Sidak's *post hoc* (SCI-COLCHI)** | | | |
| | | **Mean diff,** | **95,00% CI of diff,** | **p-value** |
| | 30 dpi | -1.918 | -4.232 to 0,3956 | 0.0863 |
| | **Tukey's *post hoc*** | | | |
| | | **Mean diff,** | **95,00% CI of diff,** | **p-value** |
| | **COLCHI** Pre vs. 30 | -1.918 | -4.232 to 0.3956 | 0.0863 |
| | 30 vs. 120 | 1.918 | -0.3956 to 4.232 | 0.0863 |
| **Correction** | **Mixed-effects model (REML)** | | | |
| | | **F (DFn, DFd)** | **p-value** | **Geisser-Greenhouse's ε** |
| | Time | F (1.583, 23.74) = 0.5045 | 0.5676 | 0.7914 |
| | Column Factor | F (1, 30) = 0.009061 | 0.9248 | |
| | Time x Column Factor | F (2, 30) = 1.495 | 0.2404 | |

**Sidak's *post hoc* (SCI-COLCHI)**

| | Mean diff, | 95,00% CI of diff, | p-value |
|---|---|---|---|
| Pre | 1.042 | -1.636 to 3.719 | 0.0776 |
| 120 | -2.441 | -5.274 to 0.3918 | 0.0776 |

**Tukey's *post hoc***

| | | Mean diff, | 95,00% CI of diff, | p-value |
|---|---|---|---|---|
| SCI | Pre vs. 30 | 1.042 | -1.636 to 3.719 | 0.3632 |
| | 30 vs. 120 | 1.042 | -1.636 to 3.719 | 0.3632 |
| COLCHI | Pre vs. 30 | -2.441 | -5.274 to 0.3918 | 0.0776 |
| | 30 vs. 120 | -2.441 | -5.274 to 0.3918 | 0.0776 |

**Partial placement**

**Two-way ANOVA**

| | SS | DF | MS | F (DFn, DFd) | p-value |
|---|---|---|---|---|---|
| Interaction | 176 | 2 | 88.01 | F (2, 20) = 0.6603 | 0.5276 |
| Time | 4713 | 2 | 2357 | F (2, 20) = 17.68 | <0.0001 |
| Column Factor | 112.5 | 1 | 112.5 | F (1, 10) = 1.418 | 0.2612 |
| Subject | 793.3 | 10 | 79.33 | F (10, 20) = 0.5952 | 0.7994 |
| Residual | 2666 | 20 | 133.3 | | |

**Tukey's *post hoc***

| | | Mean diff, | 95,00% CI of diff, | p-value |
|---|---|---|---|---|
| **SCI** | Pre vs. 30 | -29.4 | -46.26 to -12.53 | 0.0008 |
| | Pre vs. 120 | -27.47 | -44.33 to -10.60 | 0.0015 |
| | 30 vs. 120 | 1.931 | -14.93 to 18.79 | 0.9549 |
| **COLCHI** | Pre vs. 30 | -18.58 | -35.44 to -1.713 | 0.0293 |
| | Pre vs. 120 | -21.63 | -38.49 to -4.767 | 0.0108 |
| | 30 vs. 120 | -3.054 | -19.92 to 13.81 | 0.8913 |

**Correct placement**

**Two-way ANOVA**

| | SS | DF | MS | F (DFn, DFd) | p-value |
|---|---|---|---|---|---|
| Interaction | 891.9 | 2 | 445.9 | F (2, 20) = 4.064 | 0.033 |
| Time | 9498 | 2 | 4749 | F (2, 20) = 43.28 | <0.0001 |
| Column Factor | 120.3 | 1 | 120.3 | F (1, 10) = 0.8366 | 0.3819 |

| | | | | | | |
|---|---|---|---|---|---|---|
| | Subject | 1437 | 10 | 143.7 | F (10, 20) = 1.310 | 0.2902 |
| | Residual | 2194 | 20 | 109.7 | | 0.033 |
| | **Sidak's *post hoc* (SCI-COLCHI)** | | | | | |
| | | **Mean diff,** | | **95,00% CI of diff,** | | **p-value** |
| | Pre | 7.93 | | -8.132 to 23.99 | | 0.5283 |
| | 30 | -16.37 | | -32.44 to -0.3129 | | 0.0446 |
| | 120 | -2.521 | | -18.58 to 13.54 | | 0.9714 |
| | **Tukey's *post hoc*** | | | | | |
| | | | **Mean diff,** | **95,00% CI of diff,** | | **p-value** |
| | **SCI** | Pre vs. 30 | 48.81 | 33.51 to 64.11 | | <0.0001 |
| | | Pre vs. 120 | 36.94 | 21.64 to 52.24 | | <0.0001 |
| | | 30 vs. 120 | -11.87 | -27.17 to 3.431 | | 0.1476 |
| | **COLCHI** | Pre vs. 30 | 24.51 | 9.208 to 39.81 | | 0.0017 |
| | | Pre vs. 120 | 26.49 | 11.19 to 41.79 | | 0.0008 |
| | | 30 vs. 120 | 1.985 | -13.32 to 17.28 | | 0.9425 |

**Supplemental Table 3: Statistics for Fig. 3 G- % Score Category**

| | | | | | | | |
|---|---|---|---|---|---|---|---|
| **Figure 3F: Time to cross of Irregular pattern** | | | | | | | |
| **One way ANOVA** | | | | | | | |
| **SCI** | | | | **COLCHI** | | | |
| **F** | **R square** | | **p-value** | **F** | **R square** | | **p-value** |
| 7.059 | 0.5021 | | 0.0076 | 3.997 | 0.3477 | | 0.0406 |
| **Dunnett's *post hoc*** | | | | **Dunnett's *post hoc*** | | | |
| | **Mean diff** | **95,00% CI of diff** | **p-value** | | **Mean diff** | **95,00% CI of diff** | **p-value** |
| Pre vs 30 | -13.4 | -22.98 to -3.817 | 0.0075 | Pre vs 30 | -13.84 | -27.70 to 0.01185 | 0.0502 |
| Pre vs 120 | -11.02 | -20.15 to -1.880 | 0.0189 | Pre vs 120 | -13.97 | -27.83 to -0.1176 | 0.0481 |

| | | | | |
|---|---|---|---|---|
| **Figure 3G: %Score category of contralateral forelimb (Irregular pattern)** | | | | |
| **Total miss** | **Mixed-effects model (REML)** | | | |
| | | **F (DFn, DFd)** | **p-value** | **Geisser-Greenhouse's ε** |

| | | | | |
|---|---|---|---|---|
| | Time | F (1.445, 20.95) = 0.9556 | 0.3732 | 0.7224 |
| | Column Factor | F (1, 29) = 0.006887 | 0.9344 | |
| | Time x Column Factor | F (2, 29) = 0.5026 | 0.6101 | |
| **Deep slip** | **Mixed-effects model (REML)** | | | |
| | | **F (DFn, DFd)** | **p-value** | **Geisser-Greenhouse's ε** |
| | Time | F (1.223, 11.62) = 0.9527 | 0.3688 | 0.6117 |
| | Column Factor | F (1, 10) = 3.368 | 0.0964 | |
| | Time x Column Factor | F (2, 19) = 0.2460 | 0.7844 | |
| **Slight slip** | **Mixed-effects model (REML)** | | | |
| | | **F (DFn, DFd)** | **p-value** | **Geisser-Greenhouse's ε** |
| | Time | F (1.287, 12.22) = 3.456 | 0.0798 | 0.6434 |
| | Column Factor | F (1, 10) = 1.964 | 0.1914 | |
| | Time x Column Factor | F (2, 19) = 0.2757 | 0.762 | |
| **Replacement** | **Mixed-effects model (REML)** | | | |
| | | **F (DFn, DFd)** | **p-value** | **Geisser-Greenhouse's ε** |
| | Time | F (1.747, 16.59) = 0.02489 | 0.9636 | 0.8734 |
| | Column Factor | F (1, 10) = 0.1419 | 0.7143 | |
| | Time x Column Factor | F (2, 19) = 0.4642 | 0.6356 | |
| **Correction** | **Mixed-effects model (REML)** | | | |
| | | **F (DFn, DFd)** | **p-value** | **Geisser-Greenhouse's ε** |
| | Time | F (1.594, 15.15) = 1.817 | 0.1989 | 0.7971 |
| | Column Factor | F (1, 10) = 0.003323 | 0.9552 | |

| | | | | |
|---|---|---|---|---|
| | Time x Column Factor | F (2, 19) = 0.2001 | 0.8203 | |
| **Partial placement** | **Mixed-effects model (REML)** | | | |
| | | **F (DFn, DFd)** | **p-value** | **Geisser-Greenhouse's ε** |
| | Time | F (1.688, 16.03) = 4.777 | 0.0282 | 0.8439 |
| | Column Factor | F (1, 10) = 1.652 | 0.2276 | |
| | Time x Column Factor | F (2, 19) = 2.858 | 0.0822 | |

| | | | | | |
|---|---|---|---|---|---|
| **Partial placement** | **Tukey's *post hoc*** | | | | |
| | | | **Mean diff,** | **95,00% CI of diff,** | **p-value** |
| | **SCI** | Pre vs. 30 | -19.05 | -39.22 to 1.129 | 0.0596 |
| | | Pre vs. 120 | -5.274 | -34.66 to 24.11 | 0.8343 |
| | | 30 vs. 120 | 13.77 | -0.4700 to 28.01 | 0.0554 |
| | **COLCHI** | Pre vs. 30 | -3.836 | -29.22 to 21.55 | 0.8576 |
| | | Pre vs. 120 | -20.55 | -35.92 to -5.184 | 0.0167 |
| | | 30 vs. 120 | -16.72 | -36.62 to 3.189 | 0.084 |

| | | | | |
|---|---|---|---|---|
| **Correct placement** | **Mixed-effects model (REML)** | | | |
| | | **F (DFn, DFd)** | **p-value** | **Geisser-Greenhouse's ε** |
| | Time | F (1.916, 18.20) = 1.977 | 0.1683 | 0.958 |
| | Column Factor | F (1, 10) = 2.005 | 0.1872 | |
| | Time x Column Factor | F (2, 19) = 1.331 | 0.2878 | |

| | | | | |
|---|---|---|---|---|
| **Figure 3G: %Score category of Ipsilateral forelimb (Irregular pattern)** | | | | |
| **Total miss** | **Mixed-effects model (REML)** | | | |
| | | **F (DFn, DFd)** | **p-value** | **Geisser-Greenhouse's ε** |
| | Time | F (1.142, 10.85) = 11.65 | 0.0049 | 0.5709 |
| | Column Factor | F (1, 10) = 6.362 | 0.0303 | |

| | | | | |
|---|---|---|---|---|
| | Time x Column Factor | F (2, 19) = 5.432 | 0.0136 | |
| | **Sidak's *post hoc* (SCI-COLCHI)** | | | |
| | | **Mean diff,** | **95,00% CI of diff,** | **p-value** |
| | Pre | 0.0777 | -3.052 to 3.207 | 0.9998 |
| | 30 | 41.06 | -24.58 to 106.7 | 0.2003 |
| | 120 | 28.03 | -18.73 to 74.79 | 0.251 |
| | **Tukey's *post hoc*** | | | |
| | | | **Mean diff,** | **95,00% CI of diff,** | **p-value** |
| | **SCI** | Pre vs. 30 | -47.6 | -106.8 to 11.58 | 0.0948 |
| | | Pre vs. 120 | -39.15 | -84.07 to 5.760 | 0.079 |
| | | 30 vs. 120 | 8.446 | -14.69 to 31.58 | 0.466 |
| | **COLCHI** | Pre vs. 30 | -6.619 | -15.27 to 2.027 | 0.1171 |
| | | Pre vs. 120 | -11.2 | -18.32 to -4.092 | 0.0085 |
| | | 30 vs. 120 | -4.584 | -10.51 to 1.340 | 0.1135 |
| **Deep slip** | **Mixed-effects model (REML)** | | | |
| | | **F (DFn, DFd)** | **p-value** | **Geisser-Greenhouse's ε** |
| | Time | F (1.894, 18.00) = 2.124 | 0.1503 | 0.9472 |
| | Column Factor | F (1, 10) = 2.773 | 0.1269 | |
| | Time x Column Factor | F (2, 19) = 1.919 | 0.1741 | |
| **Slight slip** | **Mixed-effects model (REML)** | | | |
| | | **F (DFn, DFd)** | **p-value** | **Geisser-Greenhouse's ε** |
| | Time | F (1.653, 15.71) = 0.5792 | 0.5407 | 0.8267 |
| | Column Factor | F (1, 10) = 0.003809 | 0.952 | |
| | Time x Column Factor | F (2, 19) = 0.1160 | 0.8911 | |
| **Replacement** | **Mixed-effects model (REML)** | | | |
| | | **F (DFn, DFd)** | **p-value** | **Geisser-Greenhouse's ε** |

| | | | | | |
|---|---|---|---|---|---|
| | Time | F (1.712, 16.26) = 6.255 | 0.0121 | 0.8559 | |
| | Column Factor | F (1, 10) = 1.103 | 0.3184 | | |
| | Time x Column Factor | F (2, 19) = 0.4085 | 0.6703 | | |
| | **Tukey's *post hoc*** | | | | |
| | | | **Mean diff,** | **95,00% CI of diff,** | **p-value** |
| | **SCI** | Pre vs. 30 | 2.832 | -0.2155 to 5.880 | 0.0625 |
| | | Pre vs. 120 | 3.215 | -2.531 to 8.961 | 0.2551 |
| | | 30 vs. 120 | 0.3828 | -4.092 to 4.858 | 0.9507 |
| | **COLCHI** | Pre vs. 30 | 1.593 | -3.775 to 6.962 | 0.6271 |
| | | Pre vs. 120 | 3.106 | -0.4847 to 6.697 | 0.081 |
| | | 30 vs. 120 | 1.513 | -1.802 to 4.828 | 0.3725 |
| **Correction** | **Mixed-effects model (REML)** | | | | |
| | | **F (DFn, DFd)** | **p-value** | **Geisser-Greenhouse's ε** | |
| | Time | F (1.119, 16.22) = 13.73 | 0.0015 | 0.5594 | |
| | Column Factor | F (1, 29) = 13.27 | 0.001 | | |
| | Time x Column Factor | F (2, 29) = 0.7169 | 0.4967 | | |
| | **Tukey's *post hoc*** | | | | |
| | | | **Mean diff,** | **95,00% CI of diff,** | **p-value** |
| | **SCI** | Pre vs. 30 | 5.639 | -3.196 to 14.47 | 0.1711 |
| | | Pre vs. 120 | 5.268 | -2.036 to 12.57 | 0.1383 |
| | | 30 vs. 120 | -0.3704 | -1.892 to 1.151 | 0.6864 |
| | **COLCHI** | Pre vs. 30 | 7.824 | -0.4435 to 16.09 | 0.0604 |
| | | Pre vs. 120 | 8.862 | -2.248 to 19.97 | 0.1038 |
| | | 30 vs. 120 | 1.037 | -2.938 to 5.013 | 0.692 |
| **Partial placement** | **Mixed-effects model (REML)** | | | | |
| | | **F (DFn, DFd)** | **p-value** | **Geisser-Greenhouse's ε** | |
| | Time | F (1.517, 14.41) = 0.2262 | 0.7399 | 0.7583 | |

| | | | |
|---|---|---|---|
| | Column Factor | F (1, 10) = 0.01275 | 0.9123 |
| | Time x Column Factor | F (2, 19) = 0.4709 | 0.6316 |

| | | | | | |
|---|---|---|---|---|---|
| **Correct placement** | **Mixed-effects model (REML)** | | | | |
| | | | **F (DFn, DFd)** | **p-value** | **Geisser-Greenhouse's ε** |
| | Time | | F (1.972, 28.60) = 12.10 | 0.0002 | 0.9861 |
| | Column Factor | | F (1, 29) = 16.74 | 0.0003 | |
| | Time x Column Factor | | F (2, 29) = 8.475 | 0.0013 | |
| | **Sidak's *post hoc* (SCI-COLCHI)** | | | | |
| | | | **Mean diff,** | **95,00% CI of diff,** | **p-value** |
| | Pre | | 7.334 | -8.174 to 22.84 | 0.4987 |
| | 30 | | -30.36 | -58.36 to -2.371 | 0.0338 |
| | 120 | | -30.52 | -51.24 to -9.802 | 0.0057 |
| | **Tukey's *post hoc*** | | | | |
| | | | **Mean diff,** | **95,00% CI of diff,** | **p-value** |
| | **SCI** | Pre vs. 30 | 41.55 | 8.152 to 74.95 | 0.0246 |
| | | Pre vs. 120 | 41.38 | 11.85 to 70.92 | 0.0139 |
| | | 30 vs. 120 | -0.1666 | -21.57 to 21.24 | 0.9996 |
| | **COLCHI** | Pre vs. 30 | 3.854 | -21.99 to 29.70 | 0.8812 |
| | | Pre vs. 120 | 3.528 | -13.73 to 20.79 | 0.7925 |
| | | 30 vs. 120 | -0.3258 | -31.31 to 30.66 | 0.9994 |

| **Figure 3H: %Score category of contralateral hindlimb (Irregular pattern)** | | | | |
|---|---|---|---|---|
| **Total miss** | **Mixed-effects model (REML)** | | | |
| | | **F (DFn, DFd)** | **p-value** | **Geisser-Greenhouse's ε** |
| | Time | F (1.632, 15.50) = 28.82 | <0.0001 | 0.8158 |
| | Column Factor | F (1, 10) = 5.267 | 0.0446 | |

| | | | | | |
|---|---|---|---|---|---|
| | Time x Column Factor | F (2, 19) = 2.965 | 0.0757 | | |
| | **Sidak's *post hoc* (SCI-COLCHI)** | | | | |
| | | **Mean diff,** | **95,00% CI of diff,** | **p-value** | |
| | Pre | -0.6914 | -3.411 to 2.028 | 0.8496 | |
| | 30 | 5.381 | -2.369 to 13.13 | 0.1522 | |
| | 120 | 3.737 | -3.931 to 11.41 | 0.4682 | |
| | **Tukey's *post hoc*** | | | | |
| | | | **Mean diff,** | **95,00% CI of diff,** | **p-value** |
| | **SCI** | Pre vs. 30 | -11.77 | -18.09 to -5.457 | 0.0059 |
| | | Pre vs. 120 | -10.36 | -15.65 to -5.079 | 0.0033 |
| | | 30 vs. 120 | 1.411 | -5.712 to 8.534 | 0.7732 |
| | **COLCHI** | Pre vs. 30 | -5.701 | -9.052 to -2.350 | 0.0061 |
| | | Pre vs. 120 | -5.934 | -13.99 to 2.126 | 0.1307 |
| | | 30 vs. 120 | -0.2326 | -6.350 to 5.885 | 0.9916 |
| **Deep slip** | **Mixed-effects model (REML)** | | | | |
| | | **F (DFn, DFd)** | **p-value** | **Geisser-Greenhouse's ε** | |
| | Time | F (1.875, 17.81) = 0.3414 | 0.7018 | 0.9375 | |
| | Column Factor | F (1, 10) = 0.03465 | 0.8561 | | |
| | Time x Column Factor | F (2, 19) = 1.038 | 0.3735 | | |
| **Slight slip** | **Mixed-effects model (REML)** | | | | |
| | | **F (DFn, DFd)** | **p-value** | **Geisser-Greenhouse's ε** | |
| | Time | F (1.676, 15.92) = 4.12 | 0.0332 | 0.838 | |
| | Column Factor | F (1, 10) = 0.003538 | 0.9537 | | |
| | Time x Column Factor | F (2, 19) = 0.04048 | 0.9604 | | |
| | **Tukey's *post hoc*** | | | | |

| | | | | | |
|---|---|---|---|---|---|
| | | | **Mean diff,** | **95,00% CI of diff,** | **p-value** |
| | **SCI** | Pre vs. 30 | -2.657 | -8.446 to 3.131 | 0.3324 |
| | | Pre vs. 120 | -3.046 | -8.516 to 2.423 | 0.2576 |
| | | 30 vs. 120 | -0.3891 | -9.093 to 8.315 | 0.9862 |
| | **COLCHI** | Pre vs. 30 | -2.876 | -6.464 to 0.7117 | 0.1023 |
| | | Pre vs. 120 | -2.655 | -5.841 to 0.5298 | 0.0909 |
| | | 30 vs. 120 | 0.2207 | -4.421 to 4.863 | 0.9869 |
| **Replacement** | **Mixed-effects model (REML)** | | | | |
| | | **F (DFn, DFd)** | **p-value** | **Geisser-Greenhouse's ε** | |
| | Time | F (1.285, 12.21) = 0.5476 | 0.5164 | 0.6427 | |
| | Column Factor | F (1, 10) = 0.4338 | 0.525 | | |
| | Time x Column Factor | F (2, 19) = 1.140 | 0.3407 | | |
| **Correction** | **Mixed-effects model (REML)** | | | | |
| | | **F (DFn, DFd)** | **p-value** | **Geisser-Greenhouse's ε** | |
| | Time | F (1.205, 11.45) = 3.193 | 0.0956 | 0.6025 | |
| | Column Factor | F (1, 10) = 0.01253 | 0.9131 | | |
| | Time x Column Factor | F (2, 19) = 0.2274 | 0.7988 | | |
| **Partial placement** | **Mixed-effects model (REML)** | | | | |
| | | **F (DFn, DFd)** | **p-value** | **Geisser-Greenhouse's ε** | |
| | Time | F (1.544, 22.39) = 8.182 | 0.0039 | 0.772 | |
| | Column Factor | F (1, 29) = 0.1794 | 0.675 | | |
| | Time x Column Factor | F (2, 29) = 0.1254 | 0.8826 | | |
| | **Tukey's *post hoc*** | | | | |
| | | | **Mean diff,** | **95,00% CI of diff,** | **p-value** |
| | **SCI** | Pre vs. 30 | -22.94 | -69.88 to 24.00 | 0.2979 |

| | | | | | |
|---|---|---|---|---|---|
| | | Pre vs. 120 | -19.23 | -55.98 to 17.52 | 0.2922 |
| | | 30 vs. 120 | 3.712 | -24.16 to 31.59 | 0.8865 |
| | **COLCHI** | Pre vs. 30 | -22.37 | -49.07 to 4.336 | 0.0895 |
| | | Pre vs. 120 | -13.99 | -30.55 to 2.570 | 0.0872 |
| | | 30 vs. 120 | 8.379 | -6.015 to 22.77 | 0.2344 |
| **Correct placement** | **Mixed-effects model (REML)** | | | | |
| | | **F (DFn, DFd)** | **p-value** | **Geisser-Greenhouse's ε** | |
| | Time | F (1.527, 14.51) = 25.93 | <0.0001 | 0.7635 | |
| | Column Factor | F (1, 10) = 0.9271 | 0.3583 | | |
| | Time x Column Factor | F (2, 19) = 0.3319 | 0.7217 | | |
| | **Tukey's *post hoc*** | | | | |
| | | | **Mean diff,** | **95,00% CI of diff,** | **p-value** |
| | **SCI** | Pre vs. 30 | 39.4 | 6.224 to 72.57 | 0.0288 |
| | | Pre vs. 120 | 32.62 | 2.308 to 62.92 | 0.0385 |
| | | 30 vs. 120 | -6.782 | -21.75 to 8.183 | 0.3397 |
| | **COLCHI** | Pre vs. 30 | 31.84 | 2.778 to 60.90 | 0.0361 |
| | | Pre vs. 120 | 25.48 | 7.577 to 43.39 | 0.013 |
| | | 30 vs. 120 | -6.355 | -25.23 to 12.52 | 0.5568 |

| **Figure 3H: %Score category of ipsilateral hindlimb (Irregular pattern)** | | | | |
|---|---|---|---|---|
| **Total miss** | **Mixed-effects model (REML)** | | | |
| | | **F (DFn, DFd)** | **p-value** | **Geisser-Greenhouse's ε** |
| | Time | F (1.496, 21.69) = 28.75 | <0.0001 | 0.748 |
| | Column Factor | F (1, 29) = 7.439 | 0.0107 | |
| | Time x Column Factor | F (2, 29) = 3.075 | 0.0615 | |
| | **Sidak's *post hoc* (SCI-COLCHI)** | | | |
| | | **Mean diff,** | **95,00% CI of diff,** | **p-value** |
| | Pre | -1.061 | -3.425 to 1.303 | 0.4391 |

| | | | | | |
|---|---|---|---|---|---|
| | 30 | 9.249 | -5.461 to 23.96 | 0.235 | |
| | 120 | 5.991 | -3.063 to 15.05 | 0.2395 | |
| | **Tukey's *post hoc*** | | | | |
| | | | **Mean diff,** | **95,00% CI of diff,** | **p-value** |
| | **SCI** | Pre vs. 30 | -21.35 | -35.03 to -7.674 | 0.0112 |
| | | Pre vs. 120 | -12.03 | -19.79 to -4.260 | 0.0091 |
| | | 30 vs. 120 | 9.325 | -4.413 to 23.06 | 0.1477 |
| | **COLCHI** | Pre vs. 30 | -11.04 | -16.85 to -5.237 | 0.0037 |
| | | Pre vs. 120 | -4.975 | -12.88 to 2.933 | 0.1962 |
| | | 30 vs. 120 | 6.068 | -5.321 to 17.46 | 0.2819 |
| **Deep slip** | **Mixed-effects model (REML)** | | | | |
| | | **F (DFn, DFd)** | **p-value** | **Geisser-Greenhouse's ε** | |
| | Time | F (1.779, 16.90) = 1.611 | 0.2291 | 0.8894 | |
| | Column Factor | F (1, 10) = 6.030 | 0.0339 | | |
| | Time x Column Factor | F (2, 19) = 1.966 | 0.1674 | | |
| **Slight slip** | **Mixed-effects model (REML)** | | | | |
| | | **F (DFn, DFd)** | **p-value** | **Geisser-Greenhouse's ε** | |
| | Time | F (1.419, 20.58) = 4.441 | 0.0358 | 0.7096 | |
| | Column Factor | F (1, 29) = 0.01130 | 0.9161 | | |
| | Time x Column Factor | F (2, 29) = 0.2610 | 0.772 | | |
| | **Tukey's *post hoc*** | | | | |
| | | | **Mean diff,** | **95,00% CI of diff,** | **p-value** |
| | **SCI** | Pre vs. 30 | -3.688 | -9.529 to 2.153 | 0.1755 |
| | | Pre vs. 120 | -5.739 | -11.93 to 0.4533 | 0.0648 |
| | | 30 vs. 120 | -2.051 | -7.979 to 3.877 | 0.4975 |
| | **COLCHI** | Pre vs. 30 | -4.764 | -13.70 to 4.175 | 0.2817 |
| | | Pre vs. 120 | -4.189 | -11.79 to 3.409 | 0.2631 |
| | | 30 vs. 120 | 0.5756 | -14.03 to 15.18 | 0.991 |

| Replacement | Mixed-effects model (REML) | | | | |
|---|---|---|---|---|---|
| | | F (DFn, DFd) | p-value | Geisser-Greenhouse's ε | |
| | Time | F (1.189, 11.29) = 1.658 | 0.2278 | 0.5944 | |
| | Column Factor | F (1, 10) = 0.4980 | 0.4965 | | |
| | Time x Column Factor | F (2, 19) = 0.6173 | 0.5499 | | |
| Correction | Mixed-effects model (REML) | | | | |
| | | F (DFn, DFd) | p-value | Geisser-Greenhouse's ε | |
| | Time | F (1.000, 14.50) = 5.678 | 0.0314 | 0.5 | |
| | Column Factor | F (1, 29) = 0,8971 | 0.3514 | | |
| | Time x Column Factor | F (2, 29) = 0,8691 | 0.43 | | |
| | Tukey's *post hoc* | | | | |
| | | | Mean diff, | 95,00% CI of diff, | p-value |
| | SCI | Pre vs. 30 | -0.6667 | -2.439 to 1.105 | 0.6713 |
| | | 30 vs. 120 | 0.6667 | -1.105 to 2.439 | 0.6713 |
| | COLCHI | Pre vs. 30 | -1.524 | -3.285 to 0.2376 | 0.0964 |
| | | 30 vs. 120 | 1.524 | -0.2376 to 3.285 | 0.0964 |
| Partial placement | Mixed-effects model (REML) | | | | |
| | | F (DFn, DFd) | p-value | Geisser-Greenhouse's ε | |
| | Time | F (1.801, 26.11) = 3.188 | 0.0624 | 0.9003 | |
| | Column Factor | F (1, 29) = 8.316 | 0.0073 | | |
| | Time x Column Factor | F (2, 29) = 0.1006 | 0.9046 | | |
| Correct placement | Mixed-effects model (REML) | | | | |
| | | F (DFn, DFd) | p-value | Geisser-Greenhouse's ε | |
| | Time | F (1.786, 16,97) = 63.67 | <0.0001 | 0.8929 | |

| | | | |
|---|---|---|---|
| | Column Factor | F (1, 10) = 1.645 | 0.2286 |
| | Time x Column Factor | F (2, 19) = 3.468 | 0.052 |

| | | | | |
|---|---|---|---|---|
| **Tukey's *post hoc*** | | | | |
| | | **Mean diff,** | **95,00% CI of diff,** | **p-value** |
| **SCI** | Pre vs. 30 | 43.66 | 29.72 to 57.61 | 0.0008 |
| | Pre vs. 120 | 38.33 | 26.74 to 49.93 | 0.0003 |
| | 30 vs. 120 | -5.331 | -20.92 to 10.26 | 0.5044 |
| **COLCHI** | Pre vs. 30 | 28.71 | 8.224 to 49.20 | 0.0138 |
| | Pre vs. 120 | 22.19 | 1.432 to 42.94 | 0.0395 |
| | 30 vs. 120 | -6.524 | -18.68 to 5.628 | 0.2777 |

**Supplemental Table 2: Statistics for Fig. 4B-F: Motor behaviour test**

| | | | |
|---|---|---|---|
| **Figure 4B(i): % of walking** | | | |
| **Mixed-effects model (REML)** | | | |
| | **F (DFn, DFd)** | **p-value** | **Geisser-Greenhouse's ε** |
| Time | F (2.964, 29.64) = 7.939 | 0.0005 | 0.5928 |
| Column Factor | F (1, 10) = 5.332 | 0.0436 | |
| Time x Column Factor | F (5, 50) = 3.114 | 0.0159 | |
| **Sidak's *post hoc* (SCI-COLCHI)** | | | |
| | **Mean diff,** | **95,00% CI of diff,** | **p-value** |
| Pre | 0 | -0.5675 to 0.5675 | >0.9999 |
| 7-10 | -0.3049 | -0.8891 to 0.2793 | 0.5303 |
| 14 | -0.5287 | -1.039 to -0.01891 | 0.0415 |

| | | | |
|---|---|---|---|
| 30 | -0.2851 | -0.7946 to 0.2243 | 0.4598 |
| 60 | -0.6259 | -1.592 to 0.3405 | 0.2836 |
| 120 | -0.1526 | -0.6341 to 0.3290 | 0.8857 |

| **Tukey's *post hoc*** | | | | | | |
|---|---|---|---|---|---|---|
| **SCI** | | | | **COLCHI** | | |
| | **Mean diff,** | **95,00% CI of diff,** | **p-value** | **Mean diff,** | **95,00% CI of diff,** | **p-value** |
| Pre vs. 7-10 | 0.6189 | -0.03515 to 1.273 | 0.0615 | 0.314 | -0.2090 to 0.8371 | 0.2593 |
| Pre vs. 14 | 0.5414 | -0.1040 to 1.187 | 0.0946 | 0.01267 | -0.5196 to 0.5449 | >0.9999 |
| Pre vs. 30 | 0.4245 | -0.1705 to 1.019 | 0.1599 | 0.1394 | -0.1990 to 0.4778 | 0.5551 |
| Pre vs. 60 | 0.5309 | 0.05511 to 1.007 | 0.0326 | -0.09504 | -0.9817 to 0.7916 | 0.9959 |
| Pre vs. 120 | 0.5916 | 0.1645 to 1.019 | 0.0132 | 0.439 | -0.1426 to 1.021 | 0.1341 |
| 7-10 vs. 14 | -0.07753 | -0.4340 to 0.2789 | 0.923 | -0.3014 | -0.8424 to 0.2397 | 0.3117 |
| 7-10 vs. 30 | -0.1944 | -0.6342 to 0.2453 | 0.496 | -0.1747 | -0.4838 to 0.1345 | 0.3014 |
| 7-10 vs. 60 | -0.08808 | -0.8011 to 0.6249 | 0.9922 | -0.4091 | -1.019 to 0.2010 | 0.192 |
| 7-10 vs. 120 | -0.02733 | -0.6382 to 0.5835 | >0.9999 | 0.125 | -0.4314 to 0.6814 | 0.9137 |
| 14 vs. 30 | -0.1169 | -0.3712 to 0.1374 | 0.4635 | 0.1267 | -0.2904 to 0.5438 | 0.7786 |
| 14 vs. 60 | -0.01054 | -0.5246 to 0.5035 | >0.9999 | -0.1077 | -1.118 to 0.9029 | 0.996 |
| 14 vs. 120 | 0.0502 | -0.3996 to 0.5000 | 0.9951 | 0.4264 | 0.2993 to 0.5535 | 0.0002 |
| 30 vs. 60 | 0.1064 | -0.3028 to 0.5156 | 0.8599 | -0.2344 | -0.9033 to 0.4344 | 0.6823 |
| 30 vs. 120 | 0.1671 | -0.2231 to 0.5573 | 0.5227 | 0.2997 | -0.1124 to 0.7118 | 0.1508 |
| 60 vs. 120 | 0.06075 | -0.2301 to 0.3516 | 0.9335 | 0.5341 | -0.4610 to 1.529 | 0.3393 |

| Figure 4B(ii): Active behaviour | | | | | | |
|---|---|---|---|---|---|---|
| **Mixed-effects model (REML)** | | | | | | |
| | **F (DFn, DFd)** | | **p-value** | | **Geisser-Greenhouse's ε** | |
| Time | F (1.901, 19.01) = 17.97 | | <0.0001 | | 0.3803 | |
| Column Factor | F (1, 10) = 2.419 | | 0.151 | | | |
| Time x Column Factor | F (5, 50) = 2.659 | | 0.033 | | | |
| **Sidak's *post hoc* (SCI-COLCHI)** | | | | | | |
| | **Mean diff,** | | **95,00% CI of diff,** | | **p-value** | |
| Pre | 0 | | -0.1519 to 0.1519 | | >0.9999 | |
| 7-10 | -0.1397 | | -0.4097 to 0.1304 | | 0.5291 | |
| 14 | -0.2715 | | -0.6940 to 0.1511 | | 0.2321 | |
| 30 | -0.01655 | | -0.2873 to 0.2542 | | >0.9999 | |
| 60 | 0.1349 | | -0.1810 to 0.4508 | | 0.5902 | |
| 120 | -0.2018 | | -0.7656 to 0.3619 | | 0.8477 | |
| **Tukey's *post hoc*** | | | | | | |
| **SCI** | | | **COLCHI** | | | |
| | **Mean diff,** | **95,00% CI of diff,** | **p-value** | **Mean diff,** | **95,00% CI of diff,** | **p-value** |
| Pre vs. 7-10 | 0.1182 | -0.07266 to 0.3091 | 0.2391 | -0.02144 | -0.3665 to 0.3236 | 0.9997 |
| Pre vs. 14 | 0.1588 | -0.2628 to 0.5804 | 0.6273 | -0.1127 | -0.2675 to 0.04214 | 0.1504 |
| Pre vs. 30 | 0.0259 | -0.06266 to 0.1145 | 0.8007 | 0.009357 | -0.3472 to 0.3659 | >0.9999 |
| Pre vs. 60 | 0.01223 | -0.02714 to 0.05160 | 0.7648 | 0.1471 | -0.09877 to 0.3930 | 0.2616 |
| Pre vs. 120 | 0.6056 | 0.1117 to 1.099 | 0.0222 | 0.4038 | -0.08181 to 0.8893 | 0.0975 |
| 7-10 vs. 14 | 0.04059 | -0.2107 to 0.2919 | 0.9755 | -0.09123 | -0.3545 to 0.1720 | 0.6905 |
| 7-10 vs. 30 | -0.09231 | -0.2199 to 0.03529 | 0.1532 | 0.0308 | -0.06503 to 0.1266 | 0.743 |
| 7-10 vs. 60 | -0.106 | -0.2720 to 0.06004 | 0.2204 | 0.1686 | -0.3579 to 0.6950 | 0.7454 |

| | | | | | | |
|---|---|---|---|---|---|---|
| 7-10 vs. 120 | 0.4874 | 0.09311 to 0.8816 | 0.0214 | 0.4252 | -0.2877 to 1.138 | 0.2637 |
| 14 vs. 30 | -0.1329 | -0.4720 to 0.2062 | 0.5956 | 0.122 | -0.1496 to 0.3936 | 0.4827 |
| 14 vs. 60 | -0.1466 | -0.5374 to 0.2443 | 0.6308 | 0.2598 | -0.07293 to 0.5925 | 0.1203 |
| 14 vs. 120 | 0.4468 | -0.01814 to 0.9117 | 0.0581 | 0.5164 | -0.03113 to 1.064 | 0.0623 |
| 30 vs. 60 | -0.01367 | -0.06678 to 0.03944 | 0.8642 | 0.1378 | -0.4283 to 0.7039 | 0.8867 |
| 30 vs. 120 | 0.5797 | 0.1003 to 1.059 | 0.0235 | 0.3944 | -0.3767 to 1.165 | 0.3767 |
| 60 vs. 120 | 0.5934 | 0.09958 to 1.087 | 0.0241 | 0.2566 | 0.003339 to 0.5099 | 0.0476 |

| **Figure 4B(iii): % of Grooming** | | | |
|---|---|---|---|
| **Mixed-effects model (REML)** | | | |
| | **F (DFn, DFd)** | **p-value** | **Geisser-Greenhouse's ε** |
| Time | F (2.449, 24.49) = 2.920 | 0.0635 | 0.4899 |
| Column Factor | F (1, 10) = 14.06 | 0.0038 | |
| Time x Column Factor | F (5, 50) = 3.285 | 0.0121 | |
| **Sidak's *post hoc* (SCI-COLCHI)** | | | |
| | **Mean diff,** | **95,00% CI of diff,** | **p-value** |
| Pre | -1.667E-09 | -2.231 to 2.231 | >0.9999 |
| 7-10 | 5.935 | -2.472 to 14.34 | 0.1806 |
| 14 | 1.806 | -3.306 to 6.918 | 0.8423 |
| 30 | 7.095 | -3.386 to 17.58 | 0.2021 |
| 60 | 1.485 | -3.132 to 6.102 | 0.8636 |
| 120 | 4.162 | -0.7219 to 9.046 | 0.0963 |

| **Tukey's *post hoc*** | | | | | | |
|---|---|---|---|---|---|---|
| **SCI** | | | **COLCHI** | | | |
| | **Mean diff,** | **95,00% CI of diff,** | **p-value** | **Mean diff,** | **95,00% CI of diff,** | **p-value** |

| | | | | | | |
|---|---|---|---|---|---|---|
| Pre vs. 7-10 | -6.347 | -16.54 to 3.844 | 0.2356 | -0.4112 | -2.268 to 1.445 | 0.9179 |
| Pre vs. 14 | -2.656 | -6.133 to 0.8206 | 0.1291 | -0.8505 | -3.005 to 1.305 | 0.59 |
| Pre vs. 30 | -6.749 | -17.20 to 3.698 | 0.2133 | 0.3458 | -0.7301 to 1.422 | 0.7429 |
| Pre vs. 60 | -2.02 | -7.207 to 3.168 | 0.6009 | -0.535 | -3.371 to 2.301 | 0.9546 |
| Pre vs. 120 | -4.882 | -10.29 to 0.5222 | 0.0729 | -0.7197 | -2.464 to 1.025 | 0.5536 |
| 7-10 vs. 14 | 3.69 | -8.560 to 15.94 | 0.7836 | -0.4393 | -3.801 to 2.922 | 0.9901 |
| 7-10 vs. 30 | -0.4025 | -13.28 to 12.48 | >0.9999 | 0.757 | -0.6056 to 2.120 | 0.3136 |
| 7-10 vs. 60 | 4.327 | -2.857 to 11.51 | 0.2572 | -0.1238 | -1.965 to 1.718 | 0.9995 |
| 7-10 vs. 120 | 1.465 | -6.800 to 9.730 | 0.9644 | -0.3085 | -2.186 to 1.569 | 0.9738 |
| 14 vs. 30 | -4.093 | -14.84 to 6.654 | 0.6186 | 1.196 | -1.612 to 4.004 | 0.527 |
| 14 vs. 60 | 0.6368 | -5.676 to 6.949 | 0.9969 | 0.3155 | -3.475 to 4.106 | 0.9987 |
| 14 vs. 120 | -2.225 | -9.494 to 5.043 | 0.7739 | 0.1308 | -3.011 to 3.273 | >0.9999 |
| 30 vs. 60 | 4.73 | -2.764 to 12.22 | 0.2273 | -0.8808 | -3.369 to 1.608 | 0.675 |
| 30 vs. 120 | 1.868 | -11.16 to 14.90 | 0.9852 | -1.065 | -2.164 to 0.03342 | 0.0562 |
| 60 vs. 120 | -2.862 | -9.049 to 3.325 | 0.4585 | -0.1847 | -2.996 to 2.627 | 0.9996 |

| **Figure 4C(i): Asymmetry (leaning on both forelimbs)** | | | |
|---|---|---|---|
| **Mixed-effects model (REML)** | | | |
| | **F (DFn, DFd)** | **p-value** | **Geisser-Greenhouse's ε** |
| Time | F (1.555, 15.55) = 7.264 | 0.0089 | 0.311 |
| Column Factor | F (1, 10) = 2.885 | 0.1202 | |
| Time x Column Factor | F (5, 50) = 1.245 | 0.3024 | |
| **Tukey's *post hoc*** | | | |
| **SCI** | | **COLCHI** | |

| | Mean diff, | 95,00% CI of diff, | p-value | Mean diff, | 95,00% CI of diff, | p-value |
|---|---|---|---|---|---|---|
| Pre vs. 7-10 | 1 | 0.6151 to 1.385 | 0.000002 | 0.7007 | -0.03759 to 1.439 | 0.0608 |
| Pre vs. 14 | 0.9814 | 0.5873 to 1.375 | 0.000003 | 0.4669 | -1.418 to 2.352 | 0.88 |
| Pre vs. 30 | 0.9807 | 0.5861 to 1.375 | 0.000003 | 0.5955 | -0.8033 to 1.994 | 0.5276 |
| Pre vs. 60 | 1 | 0.6151 to 1.385 | 0.000002 | 0.3633 | -1.566 to 2.293 | 0.9549 |
| Pre vs. 120 | 1 | 0.6151 to 1.385 | 0.000002 | 0.2941 | -1.863 to 2.452 | 0.988 |
| 7-10 vs. 14 | -0.01861 | -0.06645 to 0.02923 | 0.86541 | -0.2338 | -1.841 to 1.374 | 0.9842 |
| 7-10 vs. 30 | -0.0193 | -0.06892 to 0.03031 | 0.8521 | -0.1052 | -1.019 to 0.8081 | 0.9943 |
| 7-10 vs. 60 | - | - | - | -0.3374 | -1.741 to 1.067 | 0.8914 |
| 7-10 vs. 120 | - | - | - | -0.4066 | -2.099 to 1.285 | 0.8914 |
| 14 vs. 30 | -0.0006893 | -0.002461 to 0.001083 | 0.99999 | 0.1286 | -0.7801 to 1.037 | 0.986 |
| 14 vs. 60 | 0.01861 | -0.02923 to 0.06645 | 0.86541 | -0.1036 | -1.213 to 1.006 | 0.9978 |
| 14 vs. 120 | 0.01861 | -0.02923 to 0.06645 | 0.86541 | -0.1728 | -1.093 to 0.7473 | 0.9554 |
| 30 vs. 60 | 0.0193 | -0.03031 to 0.06892 | 0.8521 | -0.2322 | -0.8248 to 0.3604 | 0.5959 |
| 30 vs. 120 | 0.0193 | -0.03031 to 0.06892 | 0.8521 | -0.3014 | -1.122 to 0.5189 | 0.6467 |
| 60 vs. 120 | - | - | - | -0.06918 | -0.4998 to 0.3615 | 0.9761 |

| **Figure 4C(ii): Asymmetry (leaning on ipsilateral forelimb)** | | | |
|---|---|---|---|
| **Mixed-effects model (REML)** | | | |
| | **F (DFn, DFd)** | **p-value** | **Geisser-Greenhouse's ε** |
| Time | F (2.067, 20.67) = 96.28 | <0.0001 | 0.4134 |
| Column Factor | F (1, 10) = 0.7159 | 0.4173 | |
| Time x Column Factor | F (5, 50) = 1.120 | 0.3621 | |
| **Tukey's *post hoc*** | | | |

| SCI | | | | COLCHI | | |
|---|---|---|---|---|---|---|
| | Mean diff, | 95,00% CI of diff, | p-value | Mean diff, | 95,00% CI of diff, | p-value |
| Pre vs. 7-10 | 0.9012 | 0.5482 to 1.254 | 0.0008 | 0.8035 | 0.5918 to 1.015 | 0.0001 |
| Pre vs. 14 | 0.8482 | 0.3125 to 1.384 | 0.0074 | 0.6812 | 0.3020 to 1.060 | 0.0042 |
| Pre vs. 30 | 0.9173 | 0.6035 to 1.231 | 0.0004 | 0.7185 | 0.4880 to 0.9490 | 0.0003 |
| Pre vs. 60 | 0.8186 | 0.2439 to 1.393 | 0.0117 | 0.7325 | 0.5082 to 0.9567 | 0.0002 |
| Pre vs. 120 | 0.8786 | 0.4528 to 1.304 | 0.0022 | 0.8161 | 0.4077 to 1.224 | 0.0025 |
| 7-10 vs. 14 | -0.05292 | -0.2787 to 0.1728 | 0.9001 | -0.1223 | -0.3423 to 0.09772 | 0.3131 |
| 7-10 vs. 30 | 0.01619 | -0.08924 to 0.1216 | 0.9802 | -0.08499 | -0.2561 to 0.08615 | 0.4003 |
| 7-10 vs. 60 | -0.08251 | -0.3062 to 0.1412 | 0.6436 | -0.07104 | -0.2485 to 0.1064 | 0.5782 |
| 7-10 vs. 120 | -0.02253 | -0.1187 to 0.07360 | 0.9001 | 0.01254 | -0.3242 to 0.3493 | >0.9999 |
| 14 vs. 30 | 0.0691 | -0.2597 to 0.3979 | 0.9319 | 0.03733 | -0.2479 to 0.3225 | 0.99 |
| 14 vs. 60 | -0.02959 | -0.2021 to 0.1429 | 0.9688 | 0.05127 | -0.1992 to 0.3018 | 0.9382 |
| 14 vs. 120 | 0.03038 | -0.09922 to 0.1600 | 0.9001 | 0.1349 | -0.2442 to 0.5139 | 0.6712 |
| 30 vs. 60 | -0.0987 | -0.3899 to 0.1925 | 0.7065 | 0.01394 | -0.07682 to 0.1047 | 0.9801 |
| 30 vs. 120 | -0.03872 | -0.2386 to 0.1612 | 0.9497 | 0.09753 | -0.1130 to 0.3080 | 0.4571 |
| 60 vs. 120 | 0.05998 | -0.1102 to 0.2301 | 0.678 | 0.08359 | -0.1309 to 0.2981 | 0.6003 |

| Figure 4C(iii): Asymmetry (leaning on contralateral forelimb) | | | | | |
|---|---|---|---|---|---|
| Two-way ANOVA | | | | | |
| | SS | DF | MS | F (DFn, DFd) | p-value |

| | | | | | |
|---|---|---|---|---|---|
| Interaction | 1.285 | 5 | 0.2569 | F (5, 50) = 1.368 | 0.2521 |
| Time | 3.649 | 5 | 0.7298 | F (5, 50) = 3.887 | 0.0047 |
| Column Factor | 4.294 | 1 | 4.294 | F (1, 10) = 3.169 | 0.1054 |
| Subject | 13.55 | 10 | 1.355 | F (10, 50) = 7.216 | <0.0001 |
| Residual | 9.388 | 50 | 0.1878 | | |

| **Tukey's *post hoc*** | | | | | | |
|---|---|---|---|---|---|---|
| | **SCI** | | | **COLCHI** | | |
| | **Mean diff,** | **95,00% CI of diff,** | **p-value** | **Mean diff,** | **95,00% CI of diff,** | **p-value** |
| Pre vs. 7-10 | -1.058 | -1.799 to -0.3164 | 0.0013 | -0.3462 | -1.088 to 0.3950 | 0.7362 |
| Pre vs. 14 | -0.8478 | -1.589 to -0.1065 | 0.0164 | -0.232 | -0.9732 to 0.5093 | 0.9376 |
| Pre vs. 30 | -0.5204 | -1.262 to 0.2209 | 0.3141 | -0.2036 | -0.9448 to 0.5377 | 0.9637 |
| Pre vs. 60 | -0.7839 | -1.525 to -0.04261 | 0.0325 | 0.01149 | -0.7298 to 0.7528 | >0.9999 |
| Pre vs. 120 | -0.8386 | -1.580 to -0.09737 | 0.0181 | -0.3475 | -1.089 to 0.3938 | 0.7333 |
| 7-10 vs. 14 | 0.2099 | -0.5314 to 0.9512 | 0.9587 | 0.1143 | -0.6270 to 0.8555 | 0.9974 |
| 7-10 vs. 30 | 0.5373 | -0.2040 to 1.279 | 0.2805 | 0.1427 | -0.5986 to 0.8840 | 0.9925 |
| 7-10 vs. 60 | 0.2738 | -0.4675 to 1.015 | 0.8813 | 0.3577 | -0.3835 to 1.099 | 0.7089 |
| 7-10 vs. 120 | 0.219 | -0.5222 to 0.9603 | 0.9506 | -0.001228 | -0.7425 to 0.7400 | >0.9999 |
| 14 vs. 30 | 0.3274 | -0.4139 to 1.069 | 0.7789 | 0.02842 | -0.7128 to 0.7697 | >0.9999 |
| 14 vs. 60 | 0.06391 | -0.6774 to 0.8052 | 0.9998 | 0.2435 | -0.4978 to 0.9847 | 0.9243 |
| 14 vs. 120 | 0.009145 | -0.7321 to 0.7504 | >0.9999 | -0.1155 | -0.8568 to 0.6258 | 0.9972 |
| 30 vs. 60 | -0.2635 | -1.005 to 0.4778 | 0.8972 | 0.215 | -0.5262 to 0.9563 | 0.9542 |
| 30 vs. 120 | -0.3183 | -1.060 to 0.4230 | 0.7985 | -0.1439 | -0.8852 to 0.5973 | 0.9922 |
| 60 vs. 120 | -0.05476 | -0.7960 to 0.6865 | >0.9999 | -0.359 | -1.100 to 0.3823 | 0.7059 |

| Figure 4D(i): Position 3 of contralateral forelimb (grooming) | | | | | | |
|---|---|---|---|---|---|---|
| **Mixed-effects model (REML)** | | | | | | |
| | **F (DFn, DFd)** | **p-value** | **Geisser-Greenhouse's ε** | | | |
| Time | F (3.273, 32.73) = 3.447 | 0.0247 | 0.6546 | | | |
| Column Factor | F (1, 10) = 3.280 | 0.1002 | | | | |
| Time x Column Factor | F (5, 50) = 3.393 | 0.0102 | | | | |
| **Sidak's *post hoc* (SCI-COLCHI)** | | | | | | |
| | **Mean diff,** | **95,00% CI of diff,** | **p-value** | | | |
| Pre | -1.667E-09 | -2.719 to 2.719 | >0.9999 | | | |
| 7-10 | -4.322 | -9.697 to 1.053 | 0.1214 | | | |
| 14 | -1.376 | -5.170 to 2.417 | 0.8385 | | | |
| 30 | -0.6337 | -4.160 to 2.893 | 0.9924 | | | |
| 60 | -1.671 | -5.007 to 1.665 | 0.5583 | | | |
| 120 | -0.2634 | -4.043 to 3.517 | >0.9999 | | | |
| **Tukey's *post hoc*** | | | | | | |
| **SCI** | | | | **COLCHI** | | |
| | **Mean diff,** | **95,00% CI of diff,** | **p-value** | **Mean diff,** | **95,00% CI of diff,** | **p-value** |
| Pre vs. 7-10 | 0.3003 | -3.686 to 4.286 | 0.9992 | -4.021 | -9.844 to 1.801 | 0.1762 |
| Pre vs. 14 | -1.428 | -6.602 to 3.745 | 0.8315 | -2.805 | -6.502 to 0.8923 | 0.132 |
| Pre vs. 30 | -0.2963 | -3.782 to 3.190 | 0.9986 | -0.9299 | -4.808 to 2.948 | 0.8922 |
| Pre vs. 60 | -0.5666 | -3.288 to 2.155 | 0.9343 | -2.237 | -6.211 to 1.736 | 0.3037 |
| Pre vs. 120 | -1.334 | -6.788 to 4.120 | 0.8849 | -1.597 | -3.446 to 0.2512 | 0.0853 |
| 7-10 vs. 14 | -1.729 | -3.599 to 0.1415 | 0.0671 | 1.217 | -3.334 to 5.767 | 0.8471 |
| 7-10 vs. 30 | -0.5966 | -2.389 to 1.196 | 0.7193 | 3.091 | -0.3954 to 6.578 | 0.0779 |
| 7-10 vs. 60 | -0.8669 | -3.070 to 1.336 | 0.5924 | 1.784 | -4.512 to 8.079 | 0.8181 |
| 7-10 vs. 120 | -1.634 | -4.256 to 0.9876 | 0.2351 | 2.424 | -2.773 to 7.621 | 0.4516 |

| | | | | | | |
|---|---|---|---|---|---|---|
| 14 vs. 30 | 1.132 | -2.284 to 4.548 | 0.7223 | 1.875 | -0.8176 to 4.567 | 0.172 |
| 14 vs. 60 | 0.8616 | -2.676 to 4.400 | 0.8864 | 0.5672 | -2.365 to 3.499 | 0.95 |
| 14 vs. 120 | 0.09425 | -3.255 to 3.444 | >0.9999 | 1.207 | -1.356 to 3.770 | 0.4435 |
| 30 vs. 60 | -0.2703 | -1.876 to 1.336 | 0.971 | -1.308 | -5.122 to 2.507 | 0.6984 |
| 30 vs. 120 | -1.038 | -3.520 to 1.444 | 0.5426 | -0.6675 | -3.323 to 1.988 | 0.8741 |
| 60 vs. 120 | -0.7674 | -4.115 to 2.580 | 0.9074 | 0.6401 | -2.420 to 3.700 | 0.9331 |

| **Figure 4D(ii): Position 4 of contralateral forelimb (grooming)** | | | | | |
|---|---|---|---|---|---|
| **Two-way ANOVA** | | | | | |
| | **SS** | **DF** | **MS** | **F (DFn, DFd)** | **p-value** |
| Interaction | 9.767 | 5 | 1.953 | F (5, 50) = 0.9773 | 0.4408 |
| Time | 24.02 | 5 | 4.804 | F (5, 50) = 2.404 | 0.0497 |
| Column Factor | 26.69 | 1 | 26.69 | F (1, 10) = 7.658 | 0.0199 |
| Subject | 34.86 | 10 | 3.486 | F (10, 50) = 1.744 | 0.0966 |
| Residual | 99.94 | 50 | 1.999 | | |

| **Tukey's *post hoc*** | | | | | | |
|---|---|---|---|---|---|---|
| **SCI** | | | **COLCHI** | | | |
| | **Mean diff,** | **95,00% CI of diff,** | **p-value** | **Mean diff,** | **95,00% CI of diff,** | **p-value** |
| Pre vs. 7-10 | -0.7597 | -3.178 to 1.659 | 0.9366 | -2.285 | -4.703 to 0.1340 | 0.0742 |
| Pre vs. 14 | -0.3022 | -2.721 to 2.116 | 0.999 | -2.586 | -5.005 to -0.1677 | 0.0296 |
| Pre vs. 30 | -0.06421 | -2.483 to 2.354 | >0.9999 | -1.629 | -4.048 to 0.7896 | 0.3592 |
| Pre vs. 60 | -0.4987 | -2.917 to 1.920 | 0.9898 | -1.849 | -4.268 to 0.5695 | 0.2275 |
| Pre vs. 120 | -1.464 | -3.882 to 0.9551 | 0.4794 | -2.046 | -4.465 to 0.3725 | 0.1417 |
| 7-10 vs. 14 | 0.4575 | -1.961 to 2.876 | 0.9931 | -0.3017 | -2.720 to 2.117 | 0.999 |

| | | | | | | |
|---|---|---|---|---|---|---|
| 7-10 vs. 30 | 0.6955 | -1.723 to 3.114 | 0.9559 | 0.6556 | -1.763 to 3.074 | 0.9656 |
| 7-10 vs. 60 | 0.261 | -2.158 to 2.680 | 0.9995 | 0.4355 | -1.983 to 2.854 | 0.9945 |
| 7-10 vs. 120 | -0.7038 | -3.122 to 1.715 | 0.9537 | 0.2385 | -2.180 to 2.657 | 0.9997 |
| 14 vs. 30 | 0.238 | -2.181 to 2.657 | 0.9997 | 0.9573 | -1.461 to 3.376 | 0.8475 |
| 14 vs. 60 | -0.1965 | -2.615 to 2.222 | 0.9999 | 0.7372 | -1.681 to 3.156 | 0.9439 |
| 14 vs. 120 | -1.161 | -3.580 to 1.257 | 0.7133 | 0.5402 | -1.878 to 2.959 | 0.9853 |
| 30 vs. 60 | -0.4345 | -2.853 to 1.984 | 0.9946 | -0.2201 | -2.639 to 2.198 | 0.9998 |
| 30 vs. 120 | -1.399 | -3.818 to 1.019 | 0.5291 | -0.4171 | -2.836 to 2.001 | 0.9955 |
| 60 vs. 120 | -0.9648 | -3.383 to 1.454 | 0.8433 | -0.197 | -2.616 to 2.222 | 0.9999 |

| **Figure 4D(iii): Position 5 of contralateral forelimb (grooming)** | | | |
|---|---|---|---|
| **Mixed-effects model (REML)** | | | |
| | **F (DFn, DFd)** | **p-value** | **Geisser-Greenhouse's ε** |
| Time | F (3.066, 30.66) = 0.3084 | 0.8232 | 0.6133 |
| Column Factor | F (1, 10) = 0.1140 | 0.7426 | |
| Time x Column Factor | F (5, 50) = 0.6585 | 0.6565 | |

| **Figure 4E: Grip Strength force** | | | |
|---|---|---|---|
| **Mixed-effects model (REML)** | | | |
| | **F (DFn, DFd)** | **p-value** | **Geisser-Greenhouse's ε** |
| Time | F (3.402, 34.02) = 40.36 | <0.0001 | 0.6803 |
| Column Factor | F (1, 10) = 2.447 | 0.1488 | |
| Time x Column Factor | F (5, 50) = 3.418 | 0.0098 | |
| **Sidak’s *post hoc* (SCI-COLCHI)** | | | |
| | **Mean diff,** | **95,00% CI of diff,** | **p-value** |

| | | | |
|---|---|---|---|
| Pre | 1.667E-09 | -0.2300 to 0.2300 | >0.9999 |
| 7-10 | -0.1332 | -0.3696 to 0.1032 | 0.4517 |
| 14 | -0.1574 | -0.4778 to 0.1630 | 0.5719 |
| 30 | -0.1867 | -0.4610 to 0.08772 | 0.2576 |
| 60 | -0.1948 | -0.5625 to 0.1730 | 0.4637 |
| 120 | 0.0721 | -0.1891 to 0.3333 | 0.9478 |

| **Tukey's *post hoc*** | | | | | | |
|---|---|---|---|---|---|---|
| **SCI** | | | | **COLCHI** | | |
| | **Mean diff,** | **95,00% CI of diff,** | **p-value** | **Mean diff,** | **95,00% CI of diff,** | **p-value** |
| Pre vs. 7-10 | 0.6064 | 0.2683 to 0.9444 | 0.0042 | 0.4731 | 0.2419 to 0.7044 | 0.0023 |
| Pre vs. 14 | 0.5382 | 0.2081 to 0.8683 | 0.0065 | 0.3808 | 0.2263 to 0.5352 | 0.0009 |
| Pre vs. 30 | 0.5293 | 0.2625 to 0.7962 | 0.0026 | 0.3427 | 0.07349 to 0.6118 | 0.019 |
| Pre vs. 60 | 0.4678 | -0.001640 to 0.9372 | 0.0507 | 0.273 | 0.04321 to 0.5028 | 0.0253 |
| Pre vs. 120 | 0.3588 | 0.2197 to 0.4979 | 0.0008 | 0.4309 | 0.1671 to 0.6946 | 0.0064 |
| 7-10 vs. 14 | -0.06819 | -0.3052 to 0.1689 | 0.81 | -0.09238 | -0.2665 to 0.08173 | 0.3481 |
| 7-10 vs. 30 | -0.07703 | -0.3177 to 0.1637 | 0.7457 | -0.1305 | -0.3819 to 0.1209 | 0.3652 |
| 7-10 vs. 60 | -0.1386 | -0.4606 to 0.1834 | 0.5186 | -0.2002 | -0.3376 to -0.06271 | 0.0107 |
| 7-10 vs. 120 | -0.2476 | -0.5138 to 0.01867 | 0.0656 | -0.04226 | -0.1803 to 0.09574 | 0.7737 |
| 14 vs. 30 | -0.008843 | -0.1961 to 0.1784 | >0.9999 | -0.0381 | -0.3746 to 0.2984 | 0.9948 |
| 14 vs. 60 | -0.0704 | -0.3158 to 0.1750 | 0.8115 | -0.1078 | -0.3558 to 0.1403 | 0.5105 |
| 14 vs. 120 | -0.1794 | -0.3968 to 0.03807 | 0.1001 | 0.05012 | -0.2110 to 0.3112 | 0.9514 |
| 30 vs. 60 | -0.06155 | -0.3193 to 0.1962 | 0.8936 | -0.06968 | -0.2996 to 0.1603 | 0.7801 |
| 30 vs. 120 | -0.1705 | -0.3206 to -0.02051 | 0.0302 | 0.08822 | -0.1534 to 0.3298 | 0.6514 |
| 60 vs. 120 | -0.109 | -0.4506 to 0.2327 | 0.7479 | 0.1579 | 0.09447 to 0.2213 | 0.0009 |

| Figure 4F: Use of forelimbs to gripping the bar in grip strength test | | | | |
|---|---|---|---|---|
| **Mixed-effects model (REML)** | | | | |
| | | **F (DFn. DFd)** | **p-value** | **Geisser-Greenhouse's ε** |
| **SCI** | Time | F (2.079. 31.19) = 1.134e-030 | >0.9999 | 0.5198 |
| | Column Factor | F (2. 15) = 20.47 | <0.0001 | |
| | Time x Column Factor | F (8. 60) = 1.420 | 0.207 | |
| **COLCHI** | Time | F (3.449. 51.73) = 1.174e-031 | >0.9999 | 0.8622 |
| | Column Factor | F (2. 15) = 7.773 | 0.0048 | |
| | Time x Column Factor | F (8. 60) = 8.459 | <0.0001 | |

**Tukey's *post hoc***

| | | Mean diff. | 95.00% CI of diff. | p-value |
|---|---|---|---|---|
| 7 DPI | **Both paws vs. One+One** | -3.333 | -38.49 to 31.82 | 0.9631 |
| | **Both paws vs. One paw** | -56.67 | -108.0 to -5.383 | 0.0319 |
| | **One+One vs. One** | -53.33 | -103.2 to -3.426 | 0.0377 |
| 14 DPI | **Both paws vs. One+One** | 70 | 25.53 to 114.5 | 0.0055 |
| | **Both paws vs. One paw** | 70 | 22.88 to 117.1 | 0.0062 |
| | **One+One vs. One** | 0 | -33.93 to 33.93 | >0.9999 |
| 30 DPI | **Both paws vs. One+One** | 10 | -54.67 to 74.67 | 0.9061 |
| | **Both paws vs. One paw** | 40 | -17.04 to 97.04 | 0.163 |
| | **One+One vs. One** | 30 | -20.09 to 80.09 | 0.2477 |
| 60 DPI | **Both paws vs. One+One** | 66.67 | 40.29 to 93.04 | 0.0003 |
| | **Both paws vs. One paw** | 63.33 | 28.18 to 98.49 | 0.0017 |
| | **One+One vs. One** | -3.333 | -35.34 to 28.67 | 0.9497 |
| 120 DPI | **Both paws vs. One+One** | -3.333 | -55.36 to 48.69 | 0.9831 |
| | **Both paws vs. One paw** | 43.33 | -2.813 to 89.48 | 0.0626 |
| | **One+One vs. One** | 46.67 | 6.758 to 86.58 | 0.0272 |

**Supplemental Table 3: Statistics for Fig. 5C-E: Sensory behaviour test**

| Figure 5C: Von frey contralateral | | | |
|---|---|---|---|
| **Mixed-effects model (REML)** | | | |
| | **F (DFn, DFd)** | **p-value** | **Geisser-Greenhouse's ε** |
| Time | F (3.607, 36.07) = 1.315 | 0.284 | 0.7215 |
| Column Factor | F (1, 10) = 24.33 | 0.0006 | |
| Time x Column Factor | F (5, 50) = 1.290 | 0.283 | |

| Figure 5C: Von frey ipsilateral | | | |
|---|---|---|---|
| **Mixed-effects model (REML)** | | | |
| | **F (DFn, DFd)** | **p-value** | **Geisser-Greenhouse's ε** |
| Time | F (3.018, 30.18) = 6.301 | 0.0019 | 0.6035 |
| Column Factor | F (1, 10) = 5.462 | 0.0415 | |
| Time x Column Factor | F (5, 50) = 2.057 | 0.0865 | |

| **Tukey's *post hoc*** | | | | | | |
|---|---|---|---|---|---|---|
| | **SCI** | | | **COLCHI** | | |
| | **Mean diff,** | **95,00% CI of diff,** | **p-value** | **Mean diff,** | **95,00% CI of diff,** | **p-value** |
| Pre vs. 7-10 | -0.06432 | -0.5930 to 0.4644 | 0.9928 | -0.4141 | -0.8647 to 0.03652 | 0.0685 |
| Pre vs. 14 | 0.01527 | -0.3768 to 0.4074 | >0.9999 | -0.1804 | -1.057 to 0.6967 | 0.9371 |

| | | | | | | |
|---|---|---|---|---|---|---|
| Pre vs. 30 | -0.141 | -0.6756 to 0.3936 | 0.8535 | -0.3849 | -1.049 to 0.2795 | 0.2836 |
| Pre vs. 60 | -0.1818 | -0.5768 to 0.2132 | 0.4627 | -0.6336 | -1.104 to -0.1632 | 0.0149 |
| Pre vs. 120 | -0.1025 | -0.4816 to 0.2766 | 0.8418 | -0.1946 | -0.9359 to 0.5467 | 0.8556 |
| 7-10 vs. 14 | 0.07959 | -0.1619 to 0.3211 | 0.7261 | 0.2337 | -0.4059 to 0.8733 | 0.651 |
| 7-10 vs. 30 | -0.07665 | -0.6041 to 0.4508 | 0.9843 | 0.02921 | -0.6680 to 0.7264 | >0.9999 |
| 7-10 vs. 60 | -0.1175 | -0.3503 to 0.1154 | 0.3877 | -0.2195 | -0.6721 to 0.2331 | 0.4194 |
| 7-10 vs. 120 | -0.03818 | -0.3976 to 0.3213 | 0.9961 | 0.2195 | -0.2408 to 0.6798 | 0.4332 |
| 14 vs. 30 | -0.1562 | -0.6069 to 0.2944 | 0.6902 | -0.2045 | -0.7304 to 0.3214 | 0.6021 |
| 14 vs. 60 | -0.1971 | -0.3650 to -0.02908 | 0.0266 | -0.4532 | -1.096 to 0.1894 | 0.1656 |
| 14 vs. 120 | -0.1178 | -0.4031 to 0.1676 | 0.5533 | -0.01418 | -0.2607 to 0.2324 | 0.9998 |
| 30 vs. 60 | -0.04082 | -0.3648 to 0.2832 | 0.9915 | -0.2487 | -0.7569 to 0.2595 | 0.4121 |
| 30 vs. 120 | 0.03847 | -0.4073 to 0.4843 | 0.9985 | 0.1903 | -0.3687 to 0.7493 | 0.7035 |
| 60 vs. 120 | 0.0793 | -0.1507 to 0.3093 | 0.6944 | 0.439 | -0.02648 to 0.9045 | 0.0623 |

| **Figure 5D: Hargreaves contralateral** | | | |
|---|---|---|---|
| **Mixed-effects model (REML)** | | | |
| | **F (DFn, DFd)** | **p-value** | **Geisser-Greenhouse's ε** |
| Time | F (3.547, 35.47) = 4.369 | 0.0073 | 0.7095 |
| Column Factor | F (1, 10) = 3.808 | 0.0796 | |
| Time x Column Factor | F (5, 50) = 2.233 | 0.0654 | |

| **Tukey's *post hoc*** | | | | | | |
|---|---|---|---|---|---|---|
| **SCI** | | | **COLCHI** | | | |
| | **Mean diff,** | **95,00% CI of diff,** | **p-value** | **Mean diff,** | **95,00% CI of diff,** | **p-value** |
| Pre vs. 7-10 | -0.2057 | -0.7930 to 0.3817 | 0.683 | -0.4155 | -0.8881 to 0.05715 | 0.0803 |

| | | | | | | |
|---|---|---|---|---|---|---|
| Pre vs. 14 | 0.04195 | -0.5361 to 0.6200 | 0.9993 | -0.1754 | -0.4210 to 0.07029 | 0.1596 |
| Pre vs. 30 | 0.05804 | -0.6606 to 0.7767 | 0.9989 | -0.3686 | -0.8101 to 0.07299 | 0.0962 |
| Pre vs. 60 | -0.02329 | -0.5996 to 0.5530 | >0.9999 | -0.1637 | -0.3731 to 0.04563 | 0.1197 |
| Pre vs. 120 | -0.08268 | -0.6197 to 0.4544 | 0.9799 | -0.1328 | -0.4890 to 0.2234 | 0.6351 |
| 7-10 vs. 14 | 0.2476 | -0.07745 to 0.5727 | 0.1303 | 0.2401 | -0.2475 to 0.7277 | 0.4072 |
| 7-10 vs. 30 | 0.2637 | -0.1521 to 0.6795 | 0.2244 | 0.0469 | -0.4029 to 0.4967 | 0.9964 |
| 7-10 vs. 60 | 0.1824 | -0.3753 to 0.7401 | 0.7313 | 0.2517 | -0.05943 to 0.5629 | 0.107 |
| 7-10 vs. 120 | 0.123 | -0.2212 to 0.4671 | 0.6679 | 0.2826 | -0.3100 to 0.8753 | 0.4333 |
| 14 vs. 30 | 0.01608 | -0.1561 to 0.1882 | 0.9978 | -0.1932 | -0.6914 to 0.3050 | 0.6042 |
| 14 vs. 60 | -0.06524 | -0.3338 to 0.2033 | 0.8874 | 0.01165 | -0.3202 to 0.3435 | >0.9999 |
| 14 vs. 120 | -0.1246 | -0.2444 to -0.004884 | 0.0429 | 0.04254 | -0.5287 to 0.6138 | 0.9993 |
| 30 vs. 60 | -0.08132 | -0.4395 to 0.2769 | 0.9105 | 0.2048 | -0.1423 to 0.5520 | 0.271 |
| 30 vs. 120 | -0.1407 | -0.4047 to 0.1233 | 0.3447 | 0.2357 | -0.2100 to 0.6815 | 0.3507 |
| 60 vs. 120 | -0.05939 | -0.2812 to 0.1625 | 0.8465 | 0.03089 | -0.3318 to 0.3935 | 0.9986 |

| **Figure 5D: Hargreaves ipsilateral** | | | |
|---|---|---|---|
| **Mixed-effects model (REML)** | | | |
| | **F (DFn, DFd)** | **p-value** | **Geisser-Greenhouse's ε** |
| Time | F (3.512, 35.12) = 2.922 | 0.0402 | 0.7023 |
| Column Factor | F (1, 10) = 1.775 | 0.2123 | |
| Time x Column Factor | F (5, 50) = 6.981 | <0.0001 | |
| **Sidak's *post hoc* (SCI-COLCHI)** | | | |
| | **Mean diff,** | **95,00% CI of diff,** | **p-value** |
| Pre | -0.003097 | -0.2920 to 0.2858 | >0.9999 |

| | | | |
|---|---|---|---|
| 7-10 | -0.01853 | -0.3638 to 0.3267 | >0.9999 |
| 14 | 0.09071 | -0.3140 to 0.4955 | 0.9783 |
| 30 | -0.5426 | -0.8998 to -0.1854 | 0.006 |
| 60 | -0.08963 | -0.5190 to 0.3397 | 0.9819 |
| 120 | 0.00946 | -0.3226 to 0.3415 | >0.9999 |

| **Tukey's *post hoc*** | | | | | | |
|---|---|---|---|---|---|---|
| **SCI** | | | | **COLCHI** | | |
| | **Mean diff,** | **95,00% CI of diff,** | **p-value** | **Mean diff,** | **95,00% CI of diff,** | **p-value** |
| Pre vs. 7-10 | -0.1952 | -0.5947 to 0.2044 | 0.4137 | -0.2106 | -0.4283 to 0.007132 | 0.0567 |
| Pre vs. 14 | -0.1017 | -0.5224 to 0.3190 | 0.8892 | -0.007893 | -0.1606 to 0.1448 | 0.9999 |
| Pre vs. 30 | 0.1392 | -0.04382 to 0.3221 | 0.131 | -0.4004 | -0.7099 to -0.09089 | 0.0177 |
| Pre vs. 60 | -0.03623 | -0.5362 to 0.4637 | 0.9993 | -0.1228 | -0.3650 to 0.1195 | 0.3841 |
| Pre vs. 120 | -0.03565 | -0.2919 to 0.2206 | 0.987 | -0.02309 | -0.3921 to 0.3459 | 0.9997 |
| 7-10 vs. 14 | 0.09347 | -0.5252 to 0.7121 | 0.9815 | 0.2027 | 0.007098 to 0.3983 | 0.0436 |
| 7-10 vs. 30 | 0.3343 | -0.1039 to 0.7726 | 0.1297 | -0.1898 | -0.5409 to 0.1613 | 0.334 |
| 7-10 vs. 60 | 0.1589 | -0.5272 to 0.8451 | 0.904 | 0.08784 | -0.1296 to 0.3053 | 0.5711 |
| 7-10 vs. 120 | 0.1595 | -0.3856 to 0.7046 | 0.8004 | 0.1875 | -0.04236 to 0.4174 | 0.104 |
| 14 vs. 30 | 0.2409 | -0.09919 to 0.5809 | 0.1634 | -0.3925 | -0.7100 to -0.07494 | 0.0215 |
| 14 vs. 60 | 0.06547 | -0.4766 to 0.6075 | 0.993 | -0.1149 | -0.2859 to 0.05613 | 0.191 |
| 14 vs. 120 | 0.06605 | -0.1536 to 0.2857 | 0.7847 | -0.0152 | -0.2928 to 0.2624 | 0.9998 |
| 30 vs. 60 | -0.1754 | -0.5454 to 0.1946 | 0.4382 | 0.2776 | -0.1562 to 0.7114 | 0.2189 |
| 30 vs. 120 | -0.1748 | -0.3380 to -0.01163 | 0.0383 | 0.3773 | 0.06456 to 0.6900 | 0.0237 |
| 60 vs. 120 | 0.0005847 | -0.4279 to 0.4291 | >0.9999 | 0.09968 | -0.2440 to 0.4434 | 0.8055 |

**Figure 5E: Randall-Selitto contralateral**

| Mixed-effects model (REML) | | | |
|---|---|---|---|
| | F (DFn, DFd) | p-value | Geisser-Greenhouse's ε |
| Time | F (2.167, 21.67) = 2.154 | 0.1371 | 0.7223 |
| Column Factor | F (1, 10) = 5.285 | 0.0443 | |
| Time x Column Factor | F (3, 30) = 2.177 | 0.1114 | |

| Figure 5E: Randall-Selitto ipsilateral | | | | | |
|---|---|---|---|---|---|
| Two-way ANOVA | | | | | |
| | SS | DF | MS | F (DFn, DFd) | p-value |
| Interaction | 0.1614 | 3 | 0.05379 | F (3, 30) = 4.928 | 0.0067 |
| Time | 0.1626 | 3 | 0.05421 | F (2.139, 21.39) = 4.967 | 0.0065 |
| Column Factor | 0.04843 | 1 | 0.04843 | F (1, 10) = 0.8265 | 0.3847 |
| Subject | 0.5859 | 10 | 0.05859 | F (10, 30) = 5.368 | 0.0002 |
| Residual | 0.3275 | 30 | 0.01092 | | |

| Sidak's *post hoc* | | | |
|---|---|---|---|
| | Mean diff, | 95,00% CI of diff, | p-value |
| Pre | 0 | -0.2275 to 0.2275 | >0.9999 |
| 30 | 0.2568 | 0.02923 to 0.4843 | 0.0214 |
| 60 | 0.04339 | -0.1841 to 0.2709 | 0.9795 |
| 120 | -0.04605 | -0.2736 to 0.1815 | 0.9745 |

| Tukey's *post hoc* | | | | | | |
|---|---|---|---|---|---|---|
| | SCI | | | COLCHI | | |
| | Mean diff, | 95,00% CI of diff, | p-value | Mean diff, | 95,00% CI of diff, | p-value |
| Pre vs. 30 | -0.07231 | -0.2363 to 0.09170 | 0.6323 | 0.1845 | 0.02044 to 0.3485 | 0.0228 |
| Pre vs. 60 | -0.04339 | -0.2074 to 0.1206 | 0.8886 | 1.667E-09 | -0.1640 to 0.1640 | >0.9999 |

| | | | | | | |
|---|---|---|---|---|---|---|
| Pre vs. 120 | 0.1519 | -0.01216 to 0.3159 | 0.0774 | 0.1058 | -0.05821 to 0.2698 | 0.3148 |
| 30 vs. 60 | 0.02893 | -0.1351 to 0.1929 | 0.963 | -0.1845 | -0.3485 to -0.02044 | 0.0228 |
| 30 vs. 120 | 0.2242 | 0.06016 to 0.3882 | 0.0044 | -0.07865 | -0.2427 to 0.08537 | 0.5677 |
| 60 vs. 120 | 0.1952 | 0.03123 to 0.3593 | 0.0148 | 0.1058 | -0.05821 to 0.2698 | 0.3148 |
